# Genomic surveillance of vancomycin-resistant *Enterococcus faecium* reveals spread of a linear plasmid conferring a nutrient utilization advantage

**DOI:** 10.1101/2021.05.07.442932

**Authors:** Mathilde Boumasmoud, Vanina Dengler Haunreiter, Tiziano A. Schweizer, Lilly Meyer, Bhavya Chakrakodi, Peter W. Schreiber, Kati Seidl, Denise Kühnert, Roger D. Kouyos, Annelies S. Zinkernagel

## Abstract

Healthcare-associated outbreaks of vancomycin-resistant *Enterococcus faecium* (VREfm) are a worldwide problem with increasing prevalence. The genomic plasticity of this hospital-adapted pathogen contributes to its efficient spread despite infection control measures. Here, we aimed to identify the genomic and phenotypic determinants of healthcare-associated transmission of VREfm. We assessed the VREfm transmission networks at the tertiary-care University Hospital of Zurich (USZ) between October 2014 and February 2018 and investigated microevolutionary dynamics of this pathogen. We performed whole-genome sequencing for the 69 VREfm isolates collected during this timeframe and assessed the population structure and variability of the vancomycin resistance transposon. Phylogenomic analysis allowed us to reconstruct transmission networks and to unveil external or indirect transmission networks, not detectable by traditional surveillance. Notably, it unveiled a persistent clone, sampled 31 times over a 29-month period. Exploring the evolutionary dynamics of this clone and characterizing the phenotypic consequences revealed the spread of a variant with decreased daptomycin susceptibility and the acquired ability to utilize N-acetyl-galactosamine (GalNAc), one of the primary constituents of the human gut mucins. This nutrient utilization advantage was conferred by a novel plasmid, termed pELF_USZ, which exhibited a linear topology. This plasmid, which was harbored by two distinct clones, was transferable by conjugation. Overall, this work provides an example of the potential of the integration of epidemiological, functional genomic and evolutionary perspectives to understand adaptation strategies contributing to the successful spread of VREfm.

**Significance statement:** Sequencing microbial pathogens causing outbreaks has become a common practice to characterize transmission networks. In addition to the signal provided by vertical evolution, bacterial genomes harbor mobile genetic elements, shared horizontally between clones. While macroevolutionary studies have revealed an important role of plasmids and genes encoding carbohydrate utilization systems in the adaptation of *Enterococcus faecium* to the hospital environment, mechanisms of dissemination and the specific function of many of these genetic determinants remain to be elucidated. Here, we characterize a plasmid providing a nutrient utilization advantage and show evidence for its clonal and horizontal spread at a local scale. Further studies integrating epidemiological, functional genomics and evolutionary perspectives will be critical to identify changes shaping the success of this pathogen.

## Background

*Enterococcus faecium* is a gut commensal that has globally emerged as a leading cause of healthcare associated infections in the past 20 years ^1^. *E. faecium* has substantially diversified into ecotypes adapted to environmental niches, one of which is the hospital ^2,3^. Worldwide, strains causing hospital-acquired infections, such as endocarditis, urinary infections or surgical site infections, most often belong to a single spreading lineage, originally designated as clonal complex 17 (CC17) ^3^ and more recently clade A1 ^2^. Strains belonging to this lineage have accumulated multiple resistance genes, enabling them to survive beta-lactam, aminoglycoside and fluoroquinolone antibiotics ^1,4^. Since the glycopeptide vancomycin has been the treatment of choice for infections caused by these multidrug resistant strains, prevalence of vancomycin resistance, conferred by the mobile inducible *van* genes operons, is increasing worldwide, making vancomycin-resistant *E. feacium* (VREfm) a top priority pathogen for the development of new treatment options by the World Health Organization ^5^. In Switzerland, VREfm prevalence was below 5% until 2010, when the first VREfm hospital outbreaks were reported ^6,7^. A marked increase in cases followed ^8–11^ and successive waves with specific clones propagated regionally, as in other countries ^12,13^. To curb the spread of these epidemic waves of VREfm clones, it is crucial to characterize transmission networks and understand their structure. While this has been traditionally achieved by contact tracing and molecular typing methods, more recent genomic approaches have contributed to a better understanding of the mechanisms underlying VREfm dissemination.

The spread of vancomycin resistance is driven by both *de novo* generation and clonal spread of resistant strains ^14,15^. Hospitals host polyclonal VREfm populations, introduced by multiple events ^16^. VREfm asymptomatic colonization, rarely identified in routine settings, is epidemiologically significant, as it represents a reservoir for transmission ^17,18^ and it often precedes infection ^19,20^. During asymptomatic colonization, which can persist for several months and involve multiple strains ^18,19^, *E. faecium* can evolve dynamically, with frequent gain and loss of genes ^21^. This phase is likely critical for adaptation of the pathogen because innovative traits introduced by horizontal gene transfer allow clones to adapt to environmental challenges and capitalize on the available growth resources ^22^.

Adaptation of *E. faecium* has been studied from a macroevolutionary perspective and a central role for carbohydrate utilization genes has been identified in the speciation of the *Enterococcus* genus ^23^ as well as further differentiation of *E. faecium* lineages ^2,24–27^. In particular, components of phosphotransferase systems (PTS) seem to play a pivotal role in adaptation. These systems couple the uptake and phosphorylation of carbohydrates based on extracellular and intracellular cues ^28^. Notably, a putative PTS (PTS^clin^) is highly enriched in clinical isolates and is involved in intestinal colonization ^27^. However, at a microevolutionary scale, the gene flow experienced by individual clones in the hospital environment has been less extensively studied and the functional consequences of the detected changes remain largely unknown ^21^. While a crucial role for *E. faecium* niche adaptation has been attributed to plasmids, most plasmid genes are poorly characterized ^24^.

In this study, we retrospectively characterized VREfm transmission networks at the tertiary-care University Hospital of Zurich (USZ) between 2014 and 2018 using a genomic approach. We explored evolutionary dynamics of the hospital-dwelling VREfm clones. Notably, we identified and characterized a novel linear plasmid.

## Results

### Isolate collection and initial characterization

Between 2011 and 2014, ≤3 cases of infection or colonization by VREfm were detected each year at the USZ. Since 2014, the detected VREfm cases have increased steadily (Fig. 1A). Between October 2014 and February 2018, a total of 69 VREfm isolates were collected from 61 patients. This collection included 45 isolates derived from colonizing strains and 24 from infecting strains. They were sampled from different materials and body sites, such as urine, feces, abdominal cavity, surgical sites, wounds and occasionally bloodstream (Supplemental Table 1). Eight different multi-locus sequence-types (ST) were identified. The most frequent were ST-203 (n=36), ST-117 (n=16) and ST-80 (n=10). Most of the isolates (92%, n=63) were *vanA* positive, while only six isolates were *vanB* positive (8%, n=6). Based on a combination of epidemiological criteria and traditional genotyping (multi-locus sequence typing, MLST, and pulsed-field-gel electrophoresis, PFGE, of *SmaI* digested total DNA), some transmission events were suspected, prompting the screening of exposed patients.

**Figure 1.**
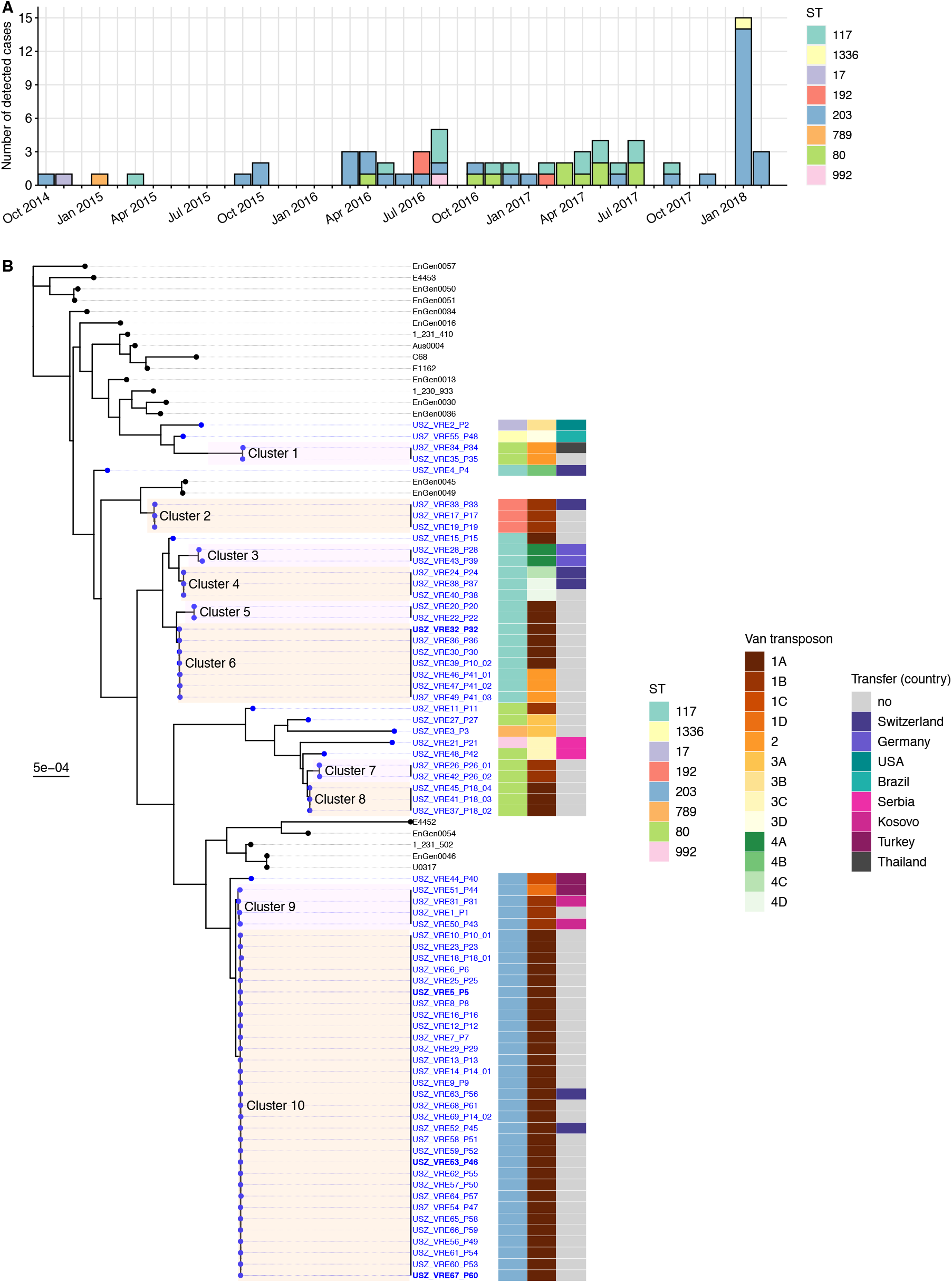
Epidemiology and population structure of VREfm at the University hospital of Zurich (USZ), Switzerland. **A** All VREfm cases detected at the USZ between October 2014 and March 2018. The colors represent the multilocus sequence type (ST) of each isolate. **B** Maximum-likelihood phylogenetic tree based on the core genes alignment (1,546,174 bp), rooted using the clade regrouping EnGen0051, EnGen0050, EnGen0057 and E4453 as an outgroup. The tips of the phylogeny marked with black dots correspond to 21 isolates collected worldwide 2 (Supplemental Table 2) those marked with blue dots correspond to the 69 isolates of this study. The labels of the latter indicate the isolation location: USZ; the unique isolate ID: VRExx and the ID of the patient from which it was isolated: Pxx. In the few cases in which multiple isolates were derived from the same patient, the last number is the index of each isolate within the longitudinal series. The four isolates with bold labels were additionally subjected to long-read sequencing to complete their assembly. Clusters, shaded with alternating colors for a clear visualization, were defined based on distances in this core gene alignment. The first column of the color map shows the ST of each isolate. The second column shows the variant of the vancomycin transposon. Three different types of transposon Tn1546, carrying the *vanA* genes, are labelled 1-3 and the subtypes they comprise further distinguished with letters and each highlighted with a different shade ranging from dark brown to light yellow. The transposon Tn1549 carrying the *vanB* genes is labelled 4 and the four subtypes it comprises further distinguished with letters and highlighted with green shades. The third column of the color map shows which patients were admitted at USZ upon transfer from another hospital and the corresponding country of the hospital.

Notably, in January and February 2018, an outbreak was suspected. Six patients suffered from various infections and eleven patients were found to be colonized with VREfm (Fig. 1A). Traditional genotyping indicated clonality of the 17 isolates and an epidemiological link (e.g., shared hospital ward) was established between all but one of the patients.

### A multiclonal population dominated by one clone

We carried out whole genome sequencing (WGS) for the 69 VREfm isolates. To contextualize their genomes in the global VREfm population, we compared them to the diverse collection of *E. faecium* genomes from Lebreton et al. 2, which documents the bifurcation of the clade grouping human-infecting strains (Clade A1) from the clades grouping most human and animal commensal strains (Clade B and Clade A2, respectively, Supplemental Table 2). The maximum-likelihood phylogenetic tree resulting from the core genes alignment of the 137 genomes showed that our isolates clustered within Clade A1 of human-infecting strains (Supplemental Fig. S1).

Next, we constructed a new maximum-likelihood tree, based on the 1,676 genes shared by our 69 isolates and the 21 isolates belonging to clade A1 from Lebreton et al. ^2^ (Fig. 1B). The resulting tree reflected both the collection’s diversity and the striking domination of a single clone. Ten isolates did not cluster with any other. The other 59 isolates were part of ten clusters, most containing two to seven isolates, except for *cluster 10*, which contained 31 isolates: the 17 isolates of the 2018 outbreak, as well as 14 earlier isolates (Supplemental Fig. S2).

To explore routes of vancomycin resistance spread, we assessed diversity of the vancomycin resistance transposon within the collection. We defined transposon variants based on the insertion sequences incorporated in the transposons, as well as on the sequence similarity to the transposon M97297 Tn1546 for isolates encoding the *vanA* genes or the transposon Aus0004 Tn1549 for isolates encoding the *vanB* genes. This led to the identification of 13 distinct transposon variants: four types, based on the combinations of insertion sequences incorporated, further divided into subtypes, based on variation compared to the reference transposons (Supplemental Fig. S3). Within clusters, the isolates’ respective transposon variants were highly concordant, illustrating the clonal spread of vancomycin resistance (Fig. 1B). Three out of the ten clusters included isolates with different variants of the transposon. Across clusters, some transposon variants were shared, suggesting that, in our setting, vancomycin resistance might have spread horizontally as well.

### Transmission networks revealed by phylogenomic analysis

Next, we compared transmission events indicated by the routine surveillance approach, which combined epidemiological data with traditional genotyping (PFGE and MLST), to the phylogenomic clusters, which we assumed to reflect transmission networks. All the transmission events suspected by routine surveillance were confirmed by the phylogenomic analysis (Supplemental Fig. S4). Two series of isolates sampled longitudinally from the same host were confirmed to be isogenic (*cluster* 7, 8). Some transmission networks partially detected by the routine surveillance were completed by the phylogenomic analysis (*cluster 6, 2, 10*). Finally, five transmission networks undetected by routine surveillance were revealed by the phylogenomic analysis (*cluster 1, 3, 4, 5, 9*) (Supplemental Table 3). To identify potential external transmission networks, we assessed which patients had been admitted to USZ after having been hospitalized in another hospital (Fig. 1B). For all transfer patients, VREfm was first detected at the USZ, sometimes multiple weeks after admission and/or after a first negative screening, except for patient 39 for whom VREfm colonization was known prior to transfer to our hospital (VRE43_P39, *cluster 3*). While in most cases the transmission event could well have happened internally, on three occasions, we observed an association between the country of previous hospitalization and either the isolates’ core genomes (*cluster 3* and *cluster 9*) or their vancomycin resistance transposon (VRE21_P21 and VRE48_P42), suggesting that in these cases, transmission networks may have been external. Comparative genomics between the isolates of *cluster 3* sampled from patients previously hospitalized in Germany (VRE28_P28 and VRE43_P43) and an isolate sampled in 2017 at a tertiary care hospital of the Rhine-Main metropolitan area by Falgenhauer et al. ^29^ (VRE01-01) revealed high similarity (18 and 38 SNPs differences respectively), further supporting the hypothesis of repeated sampling of an external transmission network.

In conclusion, the phylogenomic analysis allowed us to reconstruct indirect transmission events and to unveil potentially external transmission networks not detectable by traditional surveillance.

### Vertical evolution during longitudinal surveillance

To investigate the evolutionary dynamics of *E. faecium* clones, we first considered the clones’ vertical evolution, by comparing isolates within each cluster and removing single-nucleotide polymorphisms (SNPs) resulting from recent recombination events. Before recombination filtering, multiple clusters (*cluster 3, 4, 6, 9*) had a maximum pairwise distance >100 SNPs. After recombination filtering, the highest maximum pairwise distance (63 SNPs) was obtained in *cluster 9*, which included four isolates spanning a two-and-a-half-year period (Supplemental Fig. S2 and Supplemental Table 3).

We subsequently focused on *cluster 10*, representing the dominant clone repeatedly sampled (n=31, between September 2015 and February 2018), further referred to as the *persistent clone*. Transmission events supported by epidemiological links and consistent typing information (MLST and PFGE) had been established in three instances along this transmission chain. Overall, the sampled isolates differed pairwise by up to 12 SNPs in their core genome. The temporal signal of the phylogeny resulting from the recombination free alignment was strong, as indicated by the correlation between root-to-tip divergence and sampling time (Fig. 2A). Therefore, we performed Bayesian phylogenetic analysis to estimate the substitution rate of this clone, obtaining a rate of 3.8 substitutions per genome per year (95% credibility interval 2.3, 5.5).

**Figure 2.**
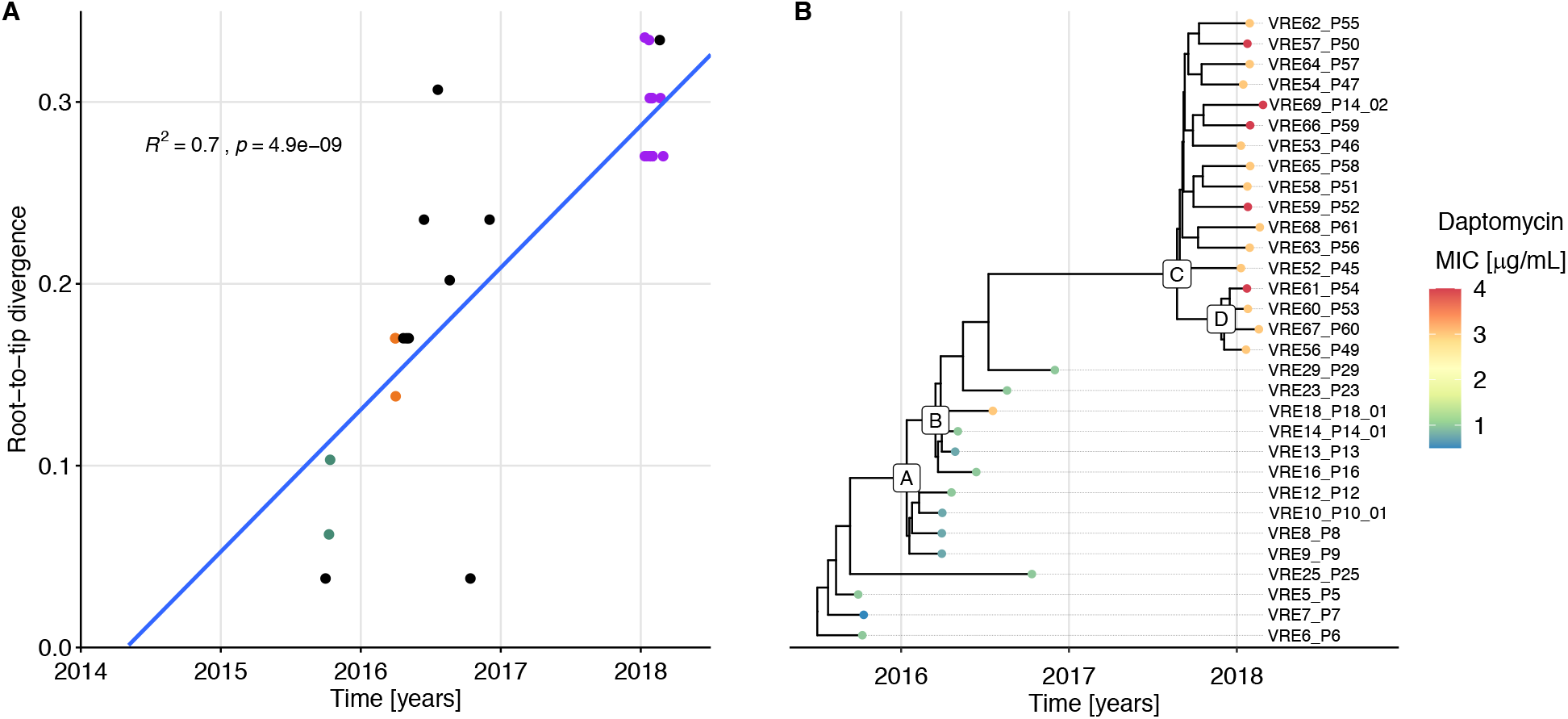
Vertical evolution of the *persistent clone*. **A** Regression of the root-to-tip divergence against sampling date, based on the phylogeny resulting from the recombination-free SNP alignment of the isolates from the *persistent clone*. R^2^ and p-value are based on a Pearson correlation. Divergence is based on the 32 bp SNPs alignment, so 0.32 corresponds to 10 SNPs. Dots corresponding to isolates from patients with an established epidemiological link share the same color, e.g., all but one of the 2018 outbreak’s isolates are in purple. All isolates from patients without established epidemiological link are in black. **B** Time-scaled maximum clade credibility (MCC) tree inferred by Bayesian analysis. The color of the dots marking the tips of the phylogeny reflects the degree of susceptibility to daptomycin (MIC: minimum inhibitory concentration). The letters labelling selected internal nodes are meant as a reference to specific clades, referred to in Table 1 and Supplemental Table 4.

Mapping each isolate to the fully assembled VRE5_P5 genome allowed us to establish a catalogue of all mutations (SNPs, insertions and deletions) differentiating the 31 isolates (Supplemental Table 4). The clade grouping the isolates of the outbreak of 2018 (*outbreak clade*, originating from node C in Fig. 2B) was characterized by six mutations as compared to the *ancestral* isolates (Table 1).

**Table 1.** Mutations characteristic of the *outbreak clade* originating from the node labelled C on Fig. 2B. The mutation with no effect (-) was in an intergenic region on plasmid 1 (pl1). All other mutations were chromosomal and non-synonymous. del: deletion, ins: insertion

| Position in VRE_5_P5 (bp) | Position in Aus0004 (bp) | Type | Length (bp) | Effect | Nucleotide change | Amino acid change | Gene | Protein |
| --- | --- | --- | --- | --- | --- | --- | --- | --- |
| 60,123 | 95,524 | SNP | 1 | Missense variant | 2032G>A | Val678Ile | *fusA* | elongation factor G |
| 389,298 | 416,003 | ins | 12 | Conservative inframe insertion | 536_537ins CGAAGACA GCAA | Asn179_Glu180 ins GluAspSerAsn | *malP* | glycoside hydrolase, family 65 |
| 1,666,026 | 1,271,778 | del | 1 | Frameshift variant | 1187delA | Lys399fs | *mvaE* | hydroxymethylglutaryl-CoA reductase |
| 1,672,764 | 1,278,516 | SNP | 1 | Missense variant | 59C>A | Ala20Asp | *cls* | cardiolipin synthase |
| 2,013,711 | 2,022,517 | SNP | 1 | Missense variant | 1261G>A | Ala421Thr | *brnQ* | branched-chain amino acid transport system II carrier protein |
| pl1: 184,545 | - | del | 63 | - | - | - | - | - |

One of these mutations was in the *cls* gene, which encodes cardiolipin synthase (Cls), an enzyme involved in phospholipid metabolism. Mutations in this gene have been associated with decreased susceptibility to daptomycin, one of the last-resort drugs used to treat VREfm. Daptomycin disrupts the cell membrane of Gram-positive bacteria and resistance is typically mediated by a few point mutations in specific genes ^30,31^. The Cls substitution Ala20Asp characterizing the *outbreak clade* isolates has been previously reported in an *E. faecium* clinical isolate belonging to an isogenic pair differing in daptomycin susceptibility ^32^. In accordance, the daptomycin minimum inhibitory concentration (MIC) was 3-4 μg/mL for the *outbreak clade* isolates and 0.5-1 μg/mL for the *ancestral* isolates, with the exception of VRE18_P18, for which the daptomycin MIC was 3 μg/mL (Fig. 2B, Supplemental Table 1, Supplemental Fig. S5).

An additional *outbreak clade*-specific mutation was a SNP in *fusA*, the gene encoding elongation factor G (EF-G). Mutations in *fusA* are associated with resistance to fusidic acid, which inhibits protein synthesis by preventing the release of the complex formed by EF-G-GDP and the ribosome ^33–35^. However, this specific mutation did not affect the fusidic acid MIC, which was 0.19 μg/mL for both VRE5_P5, *ancestral* isolate, and VRE53_P46, *outbreak clade* isolate.

In conclusion, tracking the vertical evolution of the *persistent clone* based on genomes of isolates sampled from different patients at different timepoints revealed that only few mutations accumulated over time, including a SNP leading to decreased daptomycin susceptibility.

### Horizontal evolution: acquisition of a novel plasmid

Next, we focused on changes in genome content, since the acquisition of accessory genes has the potential to confer new traits to a bacterial strain, which may be advantageous in given environments. We observed both the acquisition and the loss of genes. The most notable difference among the *persistent clone* isolates, which shared 2,802 genes, were 121 genes specific to the *outbreak clade* isolates, excluding VRE67_P60 (Fig. 3A). Most of these accessory genes had their product annotated as *hypothetical protein.* Among the 22 genes with a putative function, there was a cluster of eleven consecutive genes encoding proteins involved in carbohydrate uptake and metabolism and transcriptional regulators (Supplemental Table 5). The complete set of accessory genes was also present in two isolates from another genetic background (VRE32_P32 and VRE46_P41_01, *cluster 6*, ST-117, Supplemental Fig. S6). In the short-reads assemblies, the set of accessory genes was distributed on three to four contiguous sequences (*contigs*) exhibiting a coverage two- to three-fold higher than the assembly median coverage and closely connected based on the assembly graphs (Supplemental Fig. S7). This suggested that they may be part of a single mobile genetic element of extrachromosomal nature, i.e., a plasmid.

**Figure 3.**
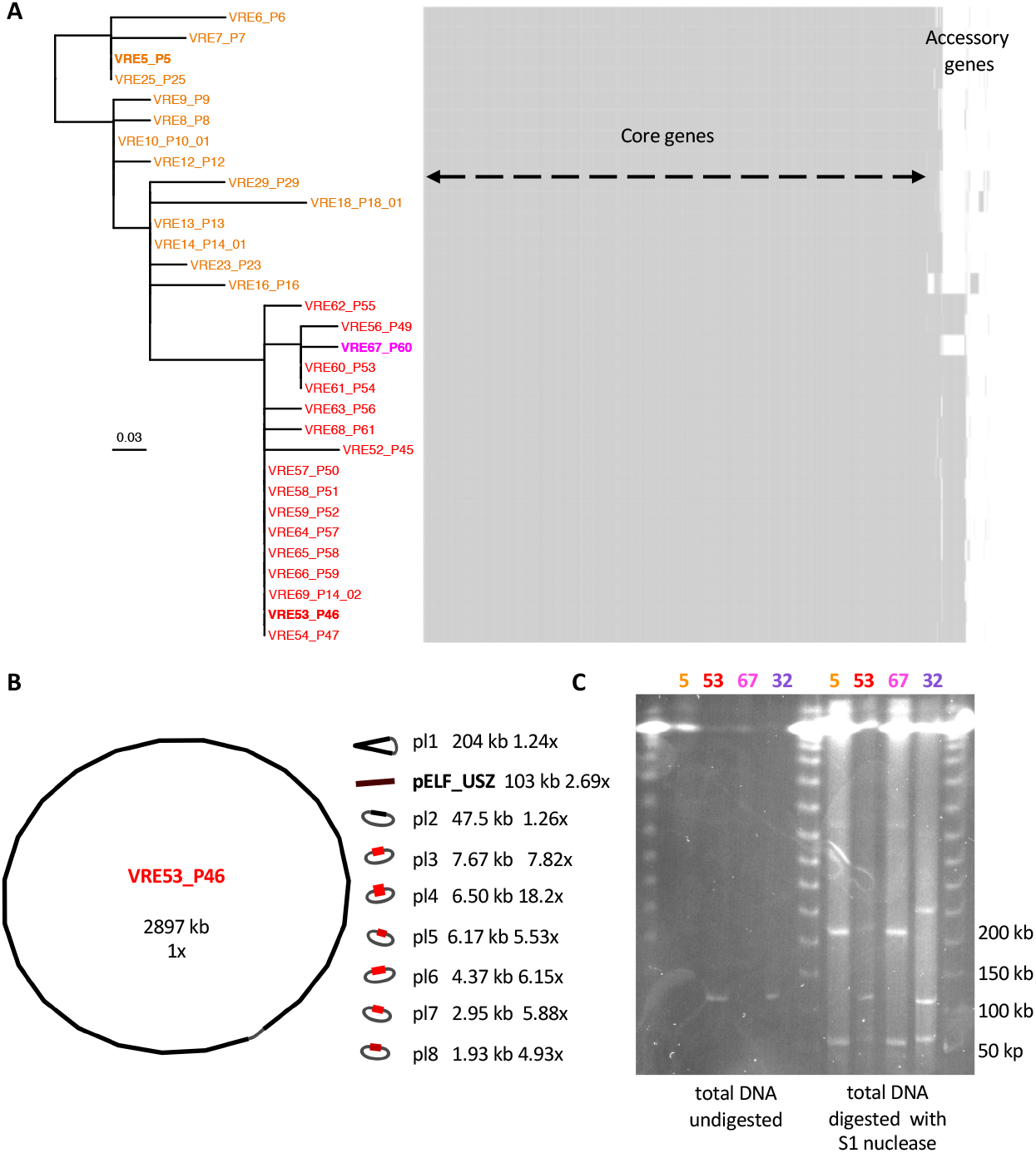
Content and structure of the genome of isolates representing the *persistent clone*. **A** Maximum likelihood phylogenetic tree along with gene content map. The phylogeny is based on the recombination-free SNPs alignment (32 bp). The *ancestral* isolates are labelled in orange, the *outbreak clade* isolates are labelled in red, excluding the only clade-member which did not carry the additional mobile genetic element, labelled in magenta. The three isolates chosen as representatives for further experiments are highlighted in bold. Each isolate has its corresponding row in the gene content map, where each column represents one gene. **B** Visualization of the complete assembly of VRE53_P46 with Bandage 36. The length of each sequence is given in kilobase-pairs (kb). Next, the coverage (depth) with respect to the chromosome coverage (set to 1x). The color, ranging from black, for the chromosome, to light red reflects the coverage and a grey string indicates circularization. **C** PFGE of three representative isolates of the *persistent clone*: VRE5_P5, VRE53_P46 and VRE67_P60 (labeled 5, 53 and 67 respectively), as well as one isolate from *cluster 6*, VRE32_P32 (labeled 32). Four columns on the left: undigested total DNA, four columns on the right: total DNA digested by S1 nuclease, for plasmid linearization.

To confirm this, we subjected selected isolates to long-read sequencing. When combining short and long reads, the mobile genetic element detected in the short-read assemblies of VRE32_P32 and VRE53_P46 was assembled into un-circularized sequences of 102 and 103 kb, respectively (Fig. 3B, Supplemental Fig. S8). Since Unicycler ^37^, the assembler we employed, attempts to circularize each contig, we assumed that these un-circularized sequences were either the result of sequencing/assembly artifacts or the result of a biological linear topology. We considered the first interpretation less plausible, as the phenomenon was consistent in two different strains/assemblies (Supplemental Fig. S8). Accordingly, we carried out molecular analyses to evaluate the possibility of a linear topology.

PFGE of the total DNA digested with S1 nuclease, which linearizes supercoiled plasmids by unspecific cleavage, revealed bands at the expected size based on the assembled plasmid sequences. One of these was at around 100 kb, corresponding to the size of the un-circularized plasmid sequence (Fig. 3C, Supplemental Fig. S8). In parallel, the control consisting of undigested total DNA also showed the same band for the isolates VRE32_P32 and VRE53_P46, supporting an original linear form. Taking up the nomenclature *pELF* for plasmid Element Linear Form introduced for the first reported *E.faecium* linear plasmids ^38,39^, we termed the linear plasmids detected in this study pELF_USZ.

### Sharing of plasmids between clones

Since two isolates from distinct clones each harbored a ~100 kb linear plasmid sequence with equivalent gene content, we sought to evaluate the similarity between these as well as all other assembled plasmid sequences of the two clones. The comparison revealed several matches: the linear plasmids, denoted pELF_USZ in both assemblies; the vancomycin resistance plasmids, denoted pl2 in both assemblies; and two smaller plasmids, denoted pl6, pl8 in VRE53_P46 and pl4, pl7 in VRE32_P32, were identical or identical except for small insertions or inversions (Supplemental Fig. S9A).

VRE53_P46 pELF_USZ was 1.5 kb longer than VRE32_P32 pELF_USZ due to the incorporation of an insertion sequence (IS256) between two coding sequences (Supplemental Fig. S9B). The sequence of the vancomycin resistance plasmid, which included several replicons and multiple resistance determinants in addition to the *vanA* operon (*catA*, *ant6-la* and *ermB*), was 3.8 kb longer in VRE32_P32 than in VRE53_P46. The additional stretch included three coding sequences (CDS) flanked by two directly oriented identical copies of the insertion sequence IS1216, forming a *pseudo compound transposon*, in place of a single copy of the same IS1216 in VRE53_P46 (Supplemental Fig. S9C). The conversion between these two configurations is known to occur with a single conservative transposition event ^40,41^.

The two patients from whom these two isolates had been sampled were never in direct contact. Yet, the fact that the two clones shared several plasmids suggested that horizontal gene transfer between them had occurred. Two other isolates of these clones (VRE10_P10_01, *cluster 10,* VRE39_P10_02, *cluster 6*) had been sampled from the same patient at an interval of 13 months. Although neither of the isolates harbored pELF_USZ, this supported plausibility of the coexistence of these two clones within the same host.

### Genetic structure of the novel linear plasmid pELF_USZ

We aimed to characterize the structure of the novel plasmid pELF_USZ. Hashimoto et al. observed a striking similarity between the backbones of the enterococcal linear plasmids pELF1 and pELF2 (143,416 kb and 108,102 kb, respectively) 38. Comparing pELF_USZ with these plasmids also uncovered a clear homology (Fig. 4). While no replication initiation protein had been detected on the three plasmids when querying them against the PlasmidFinder database 42, the RAST annotation pipeline 43 identified on each of them two CDSs encoding putative replication initiation proteins, RepB, which belonged to the Rep_3 family. None of the CDS displayed similarity with known relaxases, the other hallmark traditionally used for plasmid classification 44. Nevertheless, each plasmid encoded three proteins with a putative role in DNA partitioning and transfer: the sporulation initiation inhibitor protein Soj (ParAB), the translocase FtsK (SpoIII) and the toxin RelG (RelE/StBE) (Fig. 4, Supplemental Table 5).

**Figure 4.**
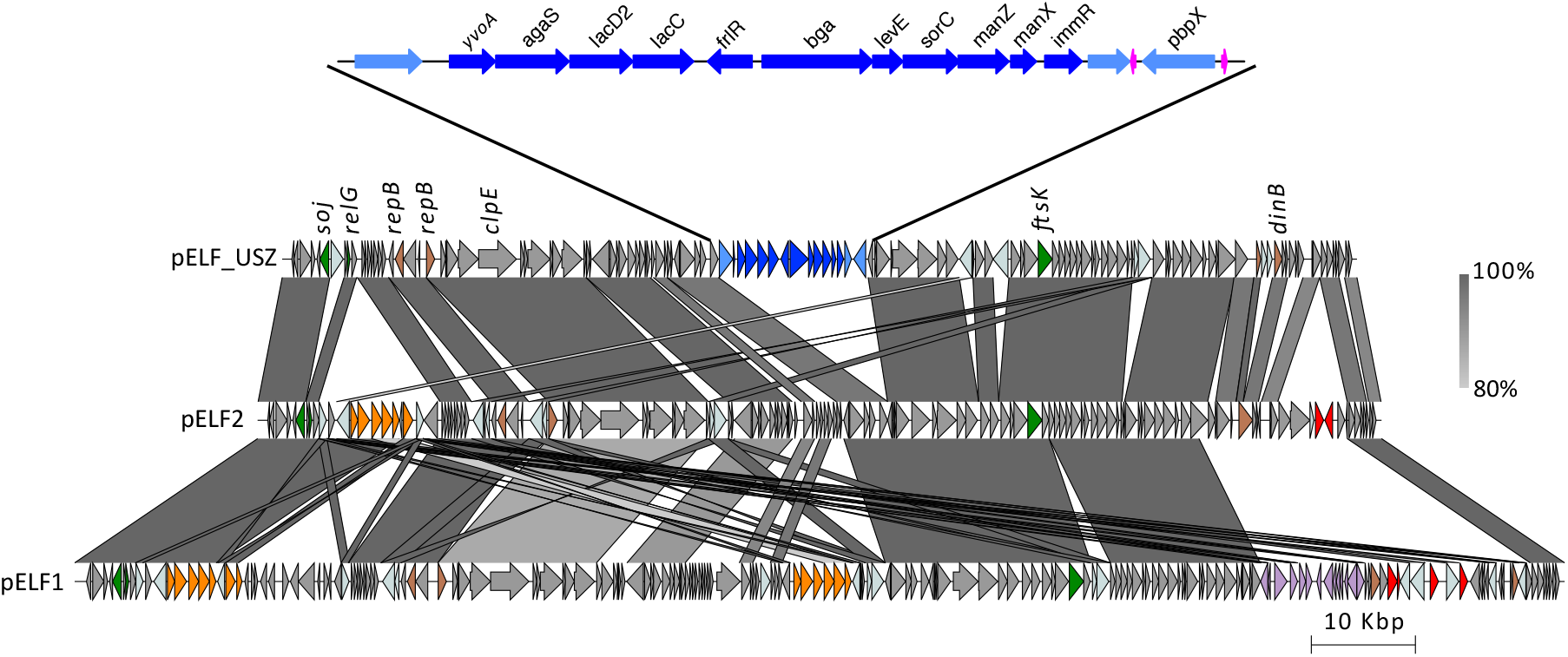
Genetic structure of pELF_USZ. Adapted and expanded from Fig. 3 of Hashimoto et al. 38. Visualization of the multiple alignment of the three plasmid sequences, produced with Easyfig 45. Regions shared between sequences are connected with vertical blocks. A minimum blast length of 500 bp and minimum identity of 80% was considered. Color gradient reflects percent identity. The coding sequences are shown with arrows and their color reflects the genes’ function: the *van* genes are in orange, the other resistance genes in red, the genes with a putative role in DNA replication are in brown, those with a putative role in DNA partitioning and transfer are in green and the insertion sequences in light grey/blue. Furthermore, an intact phage identified on pELF1 is highlighted in purple and the cargo genes of pELF_USZ are highlighted in blue. This 14 kb stretch is amplified for a clearer visualization. The eleven consecutive genes encoding a carbohydrate utilization system are in dark blue and two tRNA coding genes are in magenta.

The left extremity, reported to form a hairpin loop in pELF1 and pELF2, behaved similarly in pELF_USZ. Indeed, when assembling long reads-only, the plasmid sequence was 143 kb, that is 40 kb longer than that obtained upon hybrid assembly. Inspection of these extra 40 kb revealed that they corresponded to the inverse complement of the first 40 kb of pELF_USZ. Furthermore, the motif 5’-TATA-3’ was at the center of the loop resulting from the folding of these inverted repeats, as identified in pELF1 and pELF2 (Supplemental Fig. S10). When querying pELF_USZ against the nucleotide collection database (nr/nt), several *E. faecium* plasmids were identified as hits, and further inspection revealed that they corresponded to un-circularized sequences within their respective complete assemblies (Supplemental Table 6). The left-end inverted repeat was included in the sequence of many of them (Supplemental Fig. S11).

Within this conserved backbone, the plasmids facultatively harbored a set of cargo genes. While pELF1 and pELF2 carried resistance determinants, including *van* genes operons, pELF_USZ carried a stretch of 14 kb consisting of two consecutive operons encoding a carbohydrate degradation pathway (*agaS*, *lacD2*, *lacC*, *bga*) and a mannose-family PTS (*levE*, *sorC*, *manZ*, *manX*) interleaved with transcriptional regulators of the GntR family (*yvoA*, *flrR*) and the Xre family (*immR*) (Fig. 4, Supplemental Table 5). These operons were framed by a transposase on one side, and tRNA coding genes and *pbpX* on the other side. Since tRNA genes are normally chromosomally encoded, and this latter part was identified only in chromosomal sequences of *Enterococcus avium* assemblies upon a nucleotide collection database (nr/nt) query, this genetic determinant may have originated from the chromosome of that closely related species.

In contrast to the backbone, the set of cargo genes of pELF_USZ was not common in other *E. faecium* assemblies. A partial version, including the carbohydrate operons and transcriptional regulators (99.28% identity), but lacking the described framing CDSs, was part of one *E. faecium* plasmid assembly, pSRR24 ^46^ (Supplemental Fig. S11).

In conclusion, the homology in sequence and topology of the backbone of pELF_USZ and other *E. faecium* plasmid sequences with the backbones of pELF1 and pELF2 suggested the existence of a conserved family of linear plasmids in *E. faecium*. Moreover, the faculty of this backbone to harbor various sets of cargo genes in diverse clones suggests that it plays a part in the continuous adaptation by horizontal gene transfer of this species.

### Function and transferability of pELF_USZ

To determine the role of the carbohydrate utilization genes of pELF_USZ, we screened three representative isolates of the *persistent clone* for their ability to utilize 190 different carbon sources (Supplemental Fig. S12 and S13). We used the semi-defined medium M1, which minimizes growth of *E. faecium* when no carbon source is added ^47^. We found that 49 carbon sources sustained growth of all three isolates (Supplemental Fig. S14). Furthermore, glycerol sustained the growth of the two *outbreak clade* isolates VRE53_P46 and VRE67_P60 but not of the *ancestral* isolate VRE5_P5 (Supplemental Fig. S12 and S14). We attributed this difference and small differences in growth magnitude to the core genome mutations differentiating the three isolates. However, one condition distinguished the isolate harboring pELF_USZ (VRE53_P46) from those without pELF_USZ (VRE5_P5 and VRE67_P60): while the former could utilize N-acetyl-D-Galactosamine (GalNAc) for growth in M1, the latter two could not (Fig. 5A). Further growth assays including all isolates of the *persistent clone* confirmed that none of the *ancestral* isolates could utilize GalNAc as growth substrate, while all the *outbreak clade* isolates except for VRE67_P67 could, which confirmed that the growth advantage was conferred by the plasmid pELF_USZ (Fig. 5B, Supplemental Fig. S15A).

**Figure 5.**
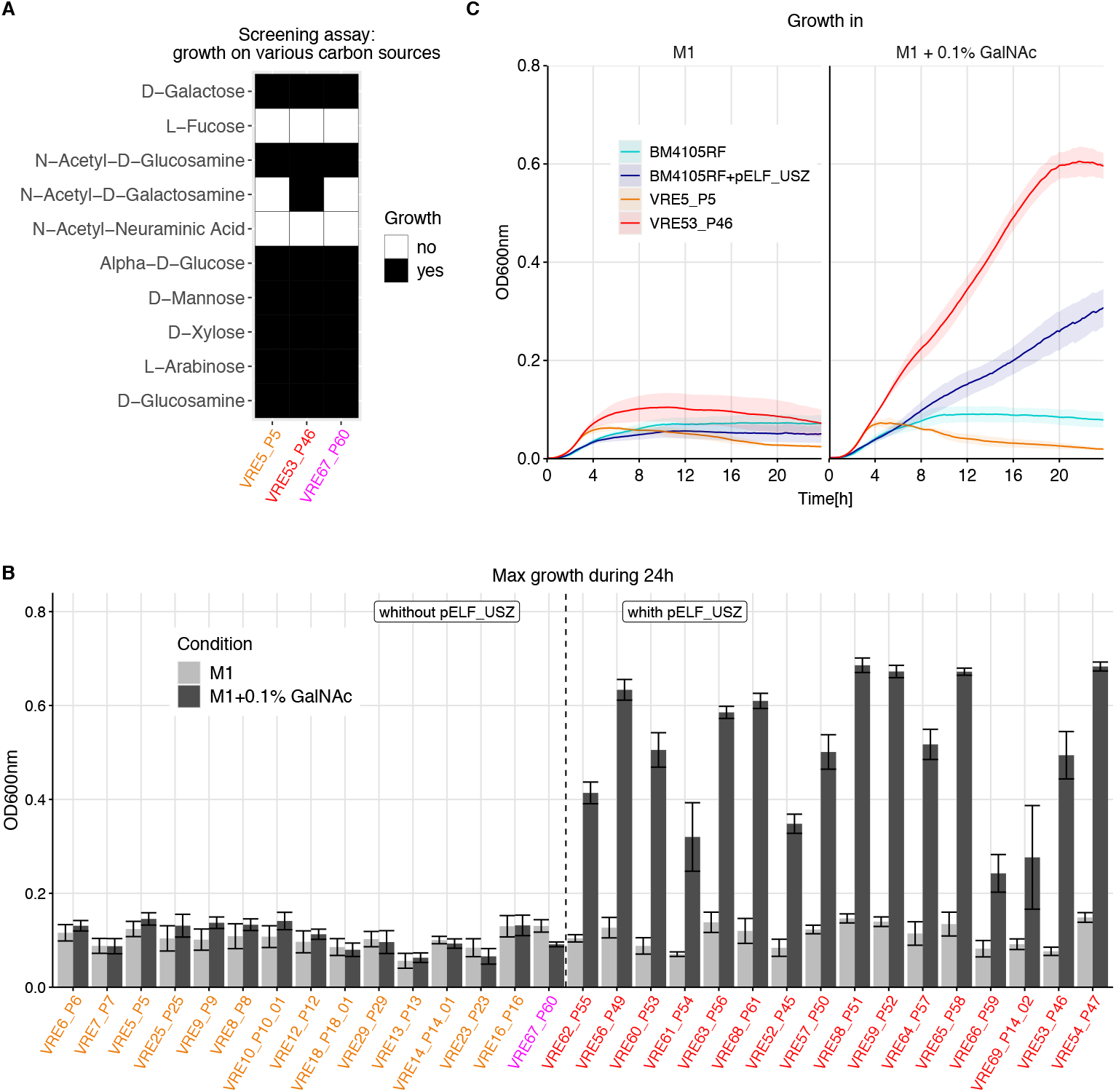
Function and transferability of pELF_USZ. **A** Growth in M1 supplemented with various carbon sources for three representative isolates of the *persistent clone*. This screening experiment was performed with the Phenotype MicroArrays PM1 and PM2A, BioLog Inc. Growth was measured by monitoring the optical density at 600nm (OD_600_) during 48 h. Conditions yielding an area under the curve above an empirically determined threshold were considered as growth permissive. Here ten out of the 190 conditions screened are displayed, selected based on their biologic relevance: the first five are monosaccharide components of mucins, and the following five are monosaccharide components of plant polysaccharides. **B** Maximum OD_600_ reached by the 31 isolates of the *persistent clone* during 24 h of growth in either M1 only or in M1 supplemented with 0.1 % GalNAc (mean and standard error of mean are displayed, N≥3). The color of the labels highlights the groups previously introduced: *ancestral* isolates are in orange and the *outbreak clade* isolates are in red, excluding the only clade member which did not carry pELF_USZ, which is in magenta. The isolates are ordered based on their position on the phylogenetic tree of Fig. 3A, except for VRE67_P60, shifted to be grouped with the isolates which did not carry pELF_USZ. The two groups, comprising isolates with or without pELF_USZ, differed significantly in their growth in M1 supplemented with GalNAc, Welch’s *t* test p= 1.17e-08. **C** Growth curves in M1 and in M1 supplemented with 0.1 % GalNAc for the recipient strain BM4105RF, the transconjugant BM4105RF+pELF_USZ, as well as two representative isolates of the *persistent clone*, VRE5_P5, which did not carry pELF_USZ, and VRE53_P46, which did carry pELF_USZ. The average of three biological replicates is shown, and the shaded area corresponds to the standard error of mean.

We supposed that this nutrient utilization trait would also distinguish the isolates of *cluster 6*, given that they differed regarding presence of pELF_USZ as well (Supplemental Fig. S6). However, we found that all seven isolates could grow in M1 supplemented with GalNAc, regardless of the presence or absence of pELF_USZ (Supplemental Fig. S15B). We therefore hypothesized that their genome included another genetic determinant conferring the same trait, rendering pELF_USZ redundant. We observed differences in the intrinsic ability to metabolize GalNAc for the other isolates of our collection (Supplemental Fig. S16). We could not pin down this trait to specific gene clusters, indicating it is likely conferred by various genetic determinants.

We then sought to verify whether pELF_USZ itself was sufficient for GalNAc utilization and, in parallel, to investigate its transferability. We performed a filter-mating experiment, using VRE53_P46 as donor and BM4105RF as recipient. The latter did not carry any plasmid and could not utilize GalNAc for growth. Upon antibiotic selection and accumulation in M1 supplemented with GalNAc, a descendant that could utilize GalNAc for growth was obtained, suggesting that pELF_USZ had successfully conjugated into BM4105RF (Fig. 5C). WGS confirmed that the presence of pELF_USZ was the only difference between the recipient and transconjugant.

Based on growth rate in rich liquid medium, the *outbreak clade* isolates had a lower fitness than the *ancestral* isolates, with their minimal doubling time (MDT) of 2 h 12 min ± 4 min versus 1 h 51 min ± 2 min for the former (Supplemental Fig. S17). The fact that the only clade member without pELF_USZ (VRE67_P60) had a minimal doubling time of 2 h 12 min suggested that this difference was not associated with the plasmid. Comparing growth dynamics of BM4105RF and BM4105RF+pELF_USZ confirmed that the presence of the plasmid was not associated with a fitness cost in the tested conditions (MDT of 1 h 27 min ± 4 min and 1 h 21 min ± 5 min, respectively, Supplemental Fig. S18). In conclusion, pELF_USZ was sufficient to confer the ability to utilize GalNAc for growth. Moreover, it was transferrable by conjugation and did not impose a measurable fitness cost to the bacterial host cells in a nutrient rich environment.

## Discussion

In this study, we characterized transmission networks of VREfm clones at the Swiss tertiary care hospital USZ, between October 2014 and February 2018. A ST-203 VREfm clone was repeatedly isolated from several patients over a time span of 29 months. During this period, a few mutations accumulated in its core genome, including a SNP leading to decreased daptomycin susceptibility. Furthermore, the clone acquired a linear plasmid, named pELF_USZ, which conferred the ability to use GalNAc for growth.

During the last decade, the superiority of genomics over traditional molecular typing methods for epidemiological surveillance of nosocomial pathogens has been repeatedly demonstrated ^48^. Indeed, MLST has a low discriminatory power, since multilocus sequence types encompass genetically diverse strains, especially for extensively recombining species like *E. faecium*. In this study some ST-80 isolates were more distantly related within their sequence type than across isolates of a different sequence type. PFGE has a higher discriminatory power but more technical limitations ^49^. In routine settings, the two methods are often used complementarily and further combined with epidemiological criteria to infer transmission events. This was the standard approach for VREfm surveillance at USZ between October 2014 and February 2018. The phylogenomic analysis presented here confirmed 100% of the transmission events suspected by this standard surveillance approach. Conversely, only 35% of the links established by the phylogenomic analysis had been suspected by the standard approach. It must be emphasized that the genetic relatedness of the clinical isolates does not allow us to establish the route, localization and directness of transmission events.

We showed that some patients carrying the same clones/resistance determinants, that were not epidemiologically linked at USZ, had been transferred to the USZ from hospitals of the same country. We concluded that these cases likely corresponded to sporadic sampling of external transmission networks. This could be further supported in the case of *cluster 3,* representing a *vanB*/ST-117 clone. This strain type has been reported to predominate over other clones in Germany in the last few years ^29,50–52^. The finding that the isolates from *cluster 3* were very closely related with one of the isolates of these studies highlights the power of genomics for global surveillance of nosocomial pathogens and the advantage of genomic data sharing at an international level.

The maximum of 12 SNPs differentiating the 31 isolates of the *persistent clone* sampled over 29 months represents an impressively low diversity as compared to that characterizing the isolates of e.g., *cluster 6* (<9 months, 21 SNPs) or *cluster 9* (32 months, 63 SNPs) (Supplemental Table 3). The low diversity of the isolates of the *persistent clone* indicated a recent emergence, consistent with the estimation by Bayesian phylogenetic analysis of a most recent common ancestor dating to 3 months (credibility interval 0.8-5.6 months) before the first sampling event (Fig. 2B) and a narrow geographical expansion, with transmission events likely to have occurred locally, at the USZ.

The substitution rate of this clone was estimated at 3.8 substitutions per genome per year, which is slightly lower but in the range of previously observed between-host evolutionary rates for *E. faecium*: five and seven substitutions per genome per year ^15,16^, respectively. Patient 14, from whom the *persistent clone* was sampled in 2016 (VRE14_P14_01) and again in 2018 (VRE69_P14_02), is the only epidemiological link between the 2018 outbreak and earlier sampling of the *persistent clone*. However, there was no evidence for prolonged carriage by this patient, who tested negative for VREfm carriage three times between the two sampling events. The clone might as well have persisted in unsampled individuals. Alternatively, given its limited evolution during the 13-month sampling gap and the well-known ability of *Enterococci* to survive on dry surfaces ^53^, a scenario in which it survived and seeded from the environment is also imaginable.

A limitation of this study was the lack of sampling of potential environmental contamination and only partial sampling of asymptomatic carriage, as screening for carriage was not carried out systematically, but only upon transmission or outbreak investigation. A recent study demonstrated that both the environment as well as asymptomatic carriage are important drivers of transmission ^18^, thus extensive sampling might have elucidated further epidemiological links.

The main strength of this study is the functional genomic analysis of longitudinally sampled isolates, allowing the identification of natural changes adopted by an individual clone persisting in the hospital environment. Indeed, while long-term persistence of VREfm clones within healthcare systems has been previously described ^15,16^, the evolution of individual hospital-dwelling clones and the horizontal gene transfer between them has not been investigated in detail.

This led to a first observation: a variant of the *persistent clone* with decreased daptomycin susceptibility spread among seventeen patients during the outbreak of 2018. In the first place, this variant was likely selected within-host, upon antimicrobial pressure. It could have evolved within one of the patients exposed to daptomycin after sampling in 2016 (e.g., P23), or prior to sampling in 2018 (i.e., P45, P52 and P65, Supplemental Fig. S5). Alternatively, it could have evolved in the off-target carriage population of any unsampled individual, as suggested by Kinnear et al. ^54^. However, while within-host evolution of variants with decreased daptomycin susceptibility is commonly reported, their clonal spread is rarer ^55–57^. In hospitals of the New York metropolitan area, a ST-376 clone harboring two mutations in the liaSR two-component system resulting in a baseline daptomycin MIC of 3-4 μg/mL was observed to spread clonally. This background of decreased daptomycin susceptibility was hypothesized to facilitate the selection of further mutations upon drug exposure, since the clone has been highly associated with daptomycin nonsusceptibility (MIC ≥8 μg/mL) ^32,58,59^. In the case presented here, further mutations leading to daptomycin nonsusceptibility have not been observed: the *persistent clone* has not been further identified by genomic surveillance up to the present.

During the same 13-month sampling gap, the *persistent clone* underwent the second genetic change highlighted in this study: the acquisition of a novel plasmid harboring carbohydrate utilization genes. Such genes have been repeatedly identified as determinants of adaptation to specific niches for *Enterococci*, yet the specific function of most of the concerned operons is unknown. In the few cases in which this was specifically investigated, it could not always be unveiled; e.g., the substrate of the mannose-family PTS enriched in the clade A1, PTS^clin^, remains to be elucidated ^27^.

The substrate of the PTS of pELF_USZ was GalNAc, and the operon accompanying it encoded enzymes necessary for breakdown of this monosaccharide, which is one of the primary components of the intestinal mucins. While *E. faecium* strains cannot degrade mucins nor plant polysaccharides, they are competing with other members of the gut microbiota for monosaccharide components released by the glycosidases secreted by anaerobes ^60^. We thus speculate that an *E. faecium* clone able to utilize GalNAc could colonize the gut more efficiently under some conditions, i.e., specific compositions of gut bacterial community, affected by the patient’s underlying illness or antibiotic treatment.

We hypothesized that the uptake of the plasmid happened within the bacterial population of one host and conferred an advantage to the subpopulation concerned, resulting in fixation and subsequent spread to other hosts. The only *outbreak clade* isolate that did not harbor the plasmid, VRE67_P60, was derived from a wound infection of the only patient for whom no epidemiological link with the other patients of the outbreak could be established. This descendant might have lost pELF_USZ in absence of selection, or conversely, it might have stemmed from an ancestor dating prior to the plasmid acquisition. Alternatively, it could have been part of a mixed population with some bacterial cells harboring the plasmid and others not. Indeed, by sampling a single isolate from each patient, we did not explore within-host diversity.

The backbone of the plasmid exhibited a linear topology, in contrast to the traditional circular topology of bacterial plasmids. Linear forms have been found in both Gram-negative and Gram-positive species ^61,62^, and while the latter mostly belong to the Actinobacteria phylum, two linear plasmids were recently identified in the genome of *E. faecium* clinical isolates in Japan ^38,39^. Both harbour vancomycin resistance genes and have a broad host range for *Enterococci*. The fact that pELF_USZ and other publicly available plasmid sequences that do not circularize had a backbone similar to that of the linear plasmids identified in Japan, suggests the existence of a conserved family of linear plasmids in *Enterococci.*

Further research is needed to assess epidemiological relevance as well as the replication and transfer mechanisms of such linear plasmids. From the bacterial genomics perspective, our finding encourages the consideration of non-circularized sequences as plausible biological occurrences. Up to now, eventual linear plasmids would have been difficult to detect with genomics, as the short-read sequencing technology does not allow to reconstruct complete assemblies. With the advent of long-read sequencing technology, genome structures are being unveiled. Yet, sequences that are un-circularized upon hybrid assembly tend to be considered as sequencing or assembly artifacts and might be disregarded in downstream analyses. Notably, a recently proposed machine-learning classifier predicting *E. faecium* plasmid-derived sequences from short-read assemblies, *mlplasmids* ^63^, was trained with 62 hybrid assemblies, from which the un-circularized sequences were excluded because they were considered as incomplete, e.g., the plasmids 3, 3, 3 and 2 of E6043, E8414, E6988 and E7196 respectively (Supplemental Fig. S11, Supplemental Table 6). As a result, while the algorithm recognized with accuracy most of the contigs of our short-read-assemblies making up plasmids, most of those making up pELF_USZ were not recognized as plasmid-derived (Supplemental Fig. S18, Supplemental Table 7). In the light of our findings, we propose including non-circularized sequences in future training datasets, to avoid perpetuating a detection bias.

So far, genomic surveillance has focused on the signal provided by vertical evolution of pathogens to track spread of clones. Recent studies started using the high resolution offered by genomic surveillance to scrutinize the dynamics of dissemination of plasmids carrying resistance determinants within healthcare systems ^64–66^. In this study, by combining epidemiological, functional genomics and evolutionary perspectives, we observed clonal and horizontal spread of a plasmid providing an innovative nutrient utilization trait, which could be adaptive under specific conditions. We envision that further studies at the microevolutionary scale will allow us to understand adaptation strategies that contribute to the success of the hospital-adapted pathogen VREfm.

## Data availability

Illumina paired-end reads of the 69 VREfm isolates; Nanopore reads and hybrid assemblies of VRE5_P5, VRE53_P46, VRE67_P60 and VRE32_P32; as well as Illumina paired-end reads of BM4105RF and BM4105RF + pELF_USZ are available through the European Nucleotide Archive project PRJEB44616.

## Authors’ contributions

K.S. and A.S.Z. initiated the project. A.S.Z. acquired funding. B.C. and K.S. collected and typed the isolates. L.M. and P.WS collected epidemiological data. M.B. performed sequencing, bioinformatic analyses and experiments. V.D.H. and T.A.S. performed preliminary experiments and contributed to the design and interpretation of experiments. D.K. and R.K. contributed to the design and interpretation of bioinformatic analyses. V.D.H., R.K. and A.S.Z. supervised the research. M.B. wrote the original manuscript draft, which was reviewed and edited by all authors.

## Acknowledgements

This work was funded by the Clinical Research Priority Program of the University of Zurich *Precision Medicine for Bacterial Infections* to A.S.Z. as well as the Swiss National Science Foundation grant 31003A_176252 to A.S.Z. The Hartmann Müller Foundation travel grant to M.B. supported presentation of part of this work at the Meeting on Microbial Epidemiological Markers XII, Dubrovnik, Croatia in September 2019. We thank Ana R. Freitas (UCIBIO, University of Porto) for kindly providing the *E. faecium* recipient strain BM4105RF.

## Materials and Methods

### Isolates collection and epidemiological characterization

From October 2014 to February 2018, 69 VREfm isolates were sampled from 61 patients during routine diagnostic and surveillance screenings at the USZ, a tertiary care facility in Switzerland. Surveillance screenings were carried out for exposed patients. Patients were considered exposed if they shared the same room as an index patient for more than 24 h on a normal ward or were in a neighboring bed of an index patient for more than 24 h on the intensive care unit. In addition, upon identification of multiple cases in the same ward, all patients hospitalized in that ward were considered exposed.

All isolates were typed with MLST according to the PubMLST typing scheme and PFGE of *SmaI* digests according to standard protocols. Transmission events were suspected by routine surveillance if isolates recovered from patients who had been exposed to each other as defined above shared the same MLST and PFGE pattern. Moreover, isolates sampled longitudinally from an individual were also considered to be linked if they showed the same MLST and PFGE pattern and no more than one intermediary negative screening. To identify potential external transmission networks, all cases of admissions to USZ upon transfer from another hospital and the corresponding country of the hospital were retrieved. To identify the isolates derived from strains having experienced daptomycin selection pressure within host, the eventual occurrence and time relative to sampling of daptomycin treatment was assessed for each patient. The necessity of a formal ethical evaluation was waived by the Zurich Cantonal Ethics Commission (req-2021-00037).

### DNA isolation, whole-genome sequencing and assembly

The DNA of the 69 clinical isolates was extracted using the DNeasy Blood & Tissue Kit (Qiagen) from single colonies inoculated in brain heart infusion broth (BHI, Bacto) and grown overnight (ON) at 37 °C, shaking at 220 r.p.m. The DNA samples were pooled into two libraries prepared with the NEBNext Ultra kit (New England BioLabs). They were sequenced on a MiSeq sequencer (Illumina) with 600-cycles paired-end runs. The quality of the paired-end reads was assessed using FastQC 0.11.15 (https://www.bioinformatics.babraham.ac.uk/projects/fastqc/) and low quality reads were filtered with Trimmomatic 0.36 67. *De novo* assemblies were constructed with SPAdes 3.10 68, with the option --careful and default k-mer sizes. Contigs with less than 2-fold average k-mer coverage and shorter than 200bp were discarded, and final assemblies were evaluated with Quast 3.1 ^69^. Detailed information is given in the Supplemental Table 8.

Four isolates were selected for long-read sequencing with the Oxford Nanopore Technology (ONT). The Gentra Puregene Kit (Qiagen) was used to extract high-molecular-weight DNA and the four DNA samples were pooled onto one MinION flow-cell and sequenced on the GridION X5 system. Hybrid and long-read-only assemblies were generated with Unicycler 0.4.8 ^37^ with default parameters.

### Phylogenomic analyses

All assemblies including the published assemblies of 73 reference genomes 2 (Supplemental Table 2) were annotated with Prokka 1.13 70. The Prokka output files were used as input to Roary 3.12 71, to construct two pangenomes: one including the 69 genomes of this study and all 73 reference genomes; and another, including the 69 genomes of this study and the 21 reference genomes belonging to clade A1. To generate core gene alignments, the command roary -e --mafft was used. These core genes alignments, of 1,131,323 bp and 1,546,174 bp respectively, were used to build maximum likelihood phylogenetic trees with FastTree 2.1.10 72, using the Jukes-Cantor substitution model. The second alignment, resulting in the phylogenetic tree of Fig.1B, was used to define clusters. A cutoff of 102 SNPs was chosen, so that, upon within-cluster whole-genome comparison followed by recombination filtering, <100 SNPs differentiated pairs of isolates of a same cluster.

The within-cluster comparisons were performed by mapping the short-reads of all isolates of a cluster to the de novo assembly of one of the isolates with Snippy 3.2 (https://github.com/tseemann/snippy). Then, the command snippy-core was used to produce a whole genome alignment, which included both variant and invariant sites. This alignment was used as input to Gubbins 2.3.1 ^73^ to identify and filter SNPs having arisen from recent recombination events.

Variant positions were extracted with SNP-sites ^74^ (with -c as input parameter) from: first, the core gene alignment used to define clusters; second, the full alignment prior to recombination filtering and third, the recombination-filtered SNPs alignment. Corresponding pairwise distance matrices were constructed with snp-dists (https://github.com/tseemann/snp-dists) (Supplemental Table 3). Maximum likelihood phylogenetic trees were built based on the recombination-free SNPs alignments with FastTree 2.1.10 ^72^ and temporal signal was assessed with TempEst 1.5.1 ^75^. The phylogeny of the *persistent clone*, which was the only one to exhibit a strong temporal signal, was rooted so as to maximize the correlation function with TempEst.

To establish a catalogue of all mutations differentiating the 31 isolates of the *persistent clone* and identify their outcome upon translation, Snippy 3.2 was used: the reads of each isolate were mapped to the fully assembled genome of VRE5_P5, as well as to the annotated reference genome Aus0004 ^76^ (Genbank accession: GCA_000250945.1) and variant-calling was performed with default parameters.

### Characterization of the vancomycin resistance transposon

The SRST2 0.2.0 tool 80 was used to screen for resistance genes in each genome, by matching the raw reads to the clustered database ARGannot_r3. Genomes including *vanA* genes were subsequently mapped to the 1,0851 bp transposon Tn1546 of BM4147 (Genbank accession: M97297.1) 81,82 and those including the *vanB* genes to the 33,810 bp transposon Tn1549 of Aus0004 (nucleotide positions 2,835,430 - 2,869,240 of the Genbank accession: CP003351) 14 using Snippy 3.2, as well as bwa-mem 83 and the SAMtools suite 84. Moreover, to assess the incorporation of insertion sequences and determine their position, the contigs of the short-read assemblies that mapped to these references were queried against the Insertion Sequence database (https://github.com/thanhleviet/ISfinder-sequences, 2020-Oct version, compiled from ISFinder 85). The alignments (coverage and SNPs) were inspected with IGV 86 and visually reproduced using the packages GenomicRanges and genoPlotR 87,88.

### Bayesian phylogenetic inference

To estimate the height of the phylogeny of the *persistent clone* (Fig. 2B) and the clock rate, Bayesian Markov Chain Monte Carlo (MCMC) phylogenetic inference implemented in BEAST 2.6.3 77 was performed. The 32 bp recombination-free SNPs alignment corrected for compositional bias was used as input. The analysis employed a Hasegawa–Kishino–Yano (HKY) substitution model, a strict clock model and a birth-death skyline serial model (BDSKY) for the tree prior ^78^. The output and chain convergence were examined with Tracer 1.6.0 ^80^ and a maximum clade credibility (MCC) tree was built with the TreeAnnotator utility of BEAST.

When assuming a 1% sampling proportion, a clock rate of 1.17·10^-6^ substitutions/site/year with a 95% highest posterior density interval of 7.07·10^-7^ to 1.68·10^-6^ substitutions/site/year and a tree height of 2.66 years with a 95% highest posterior density interval of 2.47 to 2.87 years were obtained. These posterior densities were robust when changing the sampling proportion prior to center around 10%. The whole-genome consisted of 3,281,836 sites (when summing up the length of the assembled chromosome and plasmid sequences of VRE5_P5), hence the clock rate corresponded to 3.8 substitutions/genome/year.

### Daptomycin MIC testing

Daptomycin MICs were determined using test strips (Liofilchem), which contained Ca2+ in addition to the daptomycin concentration gradient. They were placed on Mueller-Hinton II agar plates that had been inoculated with a cotton swab dipped into a 0.5 McFarland suspension. The inhibition zone was inspected after 24 h incubation at 37 °C.

### Characterization of the linear plasmid

The topology of pELF_USZ was investigated with molecular analysis and whole-genome sequences. For four representative isolates, PFGE of total DNA either undigested or digested 15 min with S1 nuclease (Promega) was carried out as described by Freitas et al. 89. Contigs’ coverage and dead ends of both the short-read and hybrid assembly graphs were inspected using Bandage 0.8.1 36 (Supplemental Fig. S7 and S8). The hybrid assembly of pELF_USZ was compared with the corresponding segment in the long-read-only assembly, and their alignment visualized using Easyfig 2.2.5 ^45^. The DNA folding of the 100 bp stretch at the center of the inverse repeats of the left extremity of pELF_USZ was inspected with mfold 3.6 90 (Supplemental Fig. S10). pELF_USZ and the other linear plasmids were screened for prophage sequences, replicons and relaxases with the web server tools PHASTER 91, PlasmidFinder 2.1 42 and MOBscan 92. They were annotated with Prokka 70 and RAST 43. CDSs with a putative product other than *hypothetical protein*, *mobile element protein*, *phage-related protein, phage protein*, *transposase*, *XX family protein* were queried against the non-redundant protein sequences (nr) database and the eggNog 5.00 database 93 to determine putative functions (Supplemental Table 5).

### Growth curves

For the growth screening assay, the Phenotype MicroArrays PM1 and PM2A (BioLog Inc) were used. Since no growth was obtained in the defined medium proposed by the manufacturer (inoculating fluid IF0a GN/GP + additive solution), the medium M1 (10 g tryptone and 0.5 g yeast extract into 1 L phosphate buffer saline, PBS), designed for minimal growth of *E. faecium* by Zhang et al. 47, was employed. Bacteria were inoculated directly from Columbia Sheep Blood agar plates (CSB, BioMérieux) into PBS at an optical density (OD) of 0.1. A mix containing 880 μl of this suspension, 11 mL of M1 and 120 μl Dye Mix H (BioLog Inc) was prepared, and 100 μl were transferred in each of the 96 wells of the PM microplates. The absorbance at 600 nm (OD600) was monitored every 10 min for 48 h with the plate-reader instrument Tecan infinite 200, while incubating at 37 °C under constant shaking (orbital, amplitude 3 mm and 5 s before measuring: linear, amplitude 1.5 mm).

Growth curves were calibrated by subtracting the baseline, i.e., the first OD_600_ value measured for each well. For each isolate and condition, the area under the average curve from three replicates was calculated (Supplemental Fig. S12 and S13). The area under the curve (AUC) for the blank, i.e., M1 only, was 133.8 ± 6.5. Conditions in which the AUC was ≥230 were considered as growth permissive. The cases for which this thresholding approach yielded a different outcome among the three isogenic isolates were closely inspected. In these cases, a few isolates/conditions for which the average growth curve had an AUC>230 but the average growth curve subtracted by the standard error of mean had an AUC<230 were attributed to noise and deemed as false positives (Supplemental Fig. S14).

To validate the role of pELF_USZ in GalNAc utilization, the 31 isolates of the *persistent clone*, BM4105RF and BM4105RF + pELF_USZ were cultivated in M1 and M1 supplemented with 0.1% GalNAc (Sigma Aldrich). Moreover, to assess their minimal doubling time in nutrient-rich liquid medium, they were grown in BHI broth. For both experiments, bacteria were inoculated in the respective media to achieve a final OD of 0.01. Technical triplicates of 200 μl from this suspension were transferred in 96 flat-wells microplates for cell culture (TPP). The OD_600_ was monitored every 10 min for 24 h, with conditions, instrument and settings described above. Doubling times were calculated for each 1 h interval, using 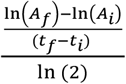, where t and A represent the timepoint and absorbance, and the indexes *i* and *f* stand for initial and final.

### Filter-mating experiment

The isolate VRE53_P46 was used as donor, and BM4105RF, a plasmid-free rifampicin and fusidic acid resistant strain 94 as recipient. The mating protocol proposed by Werner et al. 95 was followed. The two strains were grown in BHI broth ON at 37 °C, shaking at 220 r.p.m. They were then diluted 1:20 into fresh BHI medium and allowed to regrow to exponential phase, at 37 °C, shaking at 220 r.p.m, till reaching ~10^8^ CFU/mL. The two cultures were then pooled 1:1 (2 x 1mL) and centrifuged at 14,000 g for 2 min. The pellet was washed twice and finally resuspended in 1mL PBS. Of this, 100 μl were plated onto a 0.2 μm polycarbonate membrane filter (Isopore) placed on BHI agar, air-dried and incubated ON at 37 °C. The mix of donor, recipient and presumably transconjugants was then harvested from the membrane, placed into 20mL PBS and vortexed. A mix of recipient and presumably transconjugants was selected using the rifampicin and fusidic acid resistance phenotype, by plating on BHI agar containing 30 μg/mL rifampicin and 20 μg/mL fusidic acid. Cultivation of the presumed mixed population in M1 supplemented with 0.1% GalNAc resulted in significant growth, indicating that a subpopulation of transconjugants was indeed part of the inoculum, since the recipient could not grow on GalNAc. A single colony was subsequently isolated from the transconjugants-enriched population. The presence of pELF_USZ in its genome was confirmed by polymerase chain reaction, amplifying a fragment of the *agaS* gene with the primers agaS_f 5’-CCATCATTTATCAATCACCTGTGCA-3’ and agaS_r 5’-GTGTCTAAAGCCCATCGATGAATCA-3’. Moreover, the recipient and the transconjugant were subjected to whole-genome sequencing and comparative genomics, as described above.

## References

1. Arias, C. A. & Murray, B. E. The rise of the *Enterococcus*: beyond vancomycin resistance. Nat. Rev. Microbiol. 10, 266–278 (2012).

2. Lebreton, F. et al. Emergence of Epidemic Multidrug-Resistant Enterococcus faecium from Animal and Commensal Strains. mBio 4, e00534–13 (2013).

3. Top, J., Willems, R. & Bonten, M. Emergence of CC17 Enterococcus faecium: from commensal to hospital-adapted pathogen. FEMS Immunol. Med. Microbiol. 52, 297–308 (2008).

4. Gao, W., Howden, B. P. & Stinear, T. P. Evolution of virulence in Enterococcus faecium, a hospital-adapted opportunistic pathogen. Curr. Opin. Microbiol. 41, 76–82 (2018).

5. Tacconelli, E. et al. Discovery, research, and development of new antibiotics: the WHO priority list of antibiotic-resistant bacteria and tuberculosis. Lancet Infect. Dis. 18, 318–327 (2018).

6. Senn, L. et al. Unprecedented nosocoimal spread os vancomycin-resistant Enterococcus faecium in a tertiary-care hospital in Switzerland. BMC Proc. 5, P22 (2011).

7. Thierfelder, C. et al. Vancomycin-resistant Enterococcus. Swiss Med. Wkly. 142, (2012).

8. Abdelbary, M. H. H. et al. Whole-genome sequencing revealed independent emergence of vancomycin-resistant Enterococcus faecium causing sequential outbreaks over 3 years in a tertiary care hospital. Eur. J. Clin. Microbiol. Infect. Dis. Off. Publ. Eur. Soc. Clin. Microbiol. 38, 1163–1170 (2019).

9. Buetti, N. et al. Emergence of vancomycin-resistant enterococci in Switzerland: a nation-wide survey. Antimicrob. Resist. Infect. Control 8, 16 (2019).

10. Metsini, A. et al. Point prevalence of healthcare-associated infections and antibiotic use in three large Swiss acute-care hospitals. Swiss Med. Wkly. 148, (2018).

11. Wassilew, N. et al. Outbreak of vancomycin-resistant Enterococcus faecium clone ST796, Switzerland, December 2017 to April 2018. Eurosurveillance 23, 1800351 (2018).

12. Buultjens, A. H. et al. Evolutionary origins of the emergent ST796 clone of vancomycin resistant Enterococcus faecium. PeerJ 5, (2017).

13. Freitas, A. R. et al. Microevolutionary Events Involving Narrow Host Plasmids Influences Local Fixation of Vancomycin-Resistance in Enterococcus Populations. PLOS ONE 8, e60589 (2013).

14. van Hal, S. J. et al. Evolutionary dynamics of Enterococcus faecium reveals complex genomic relationships between isolates with independent emergence of vancomycin resistance. Microb. Genomics 2, (2016).

15. Howden, B. P. et al. Genomic Insights to Control the Emergence of Vancomycin-Resistant Enterococci. mBio 4, e00412–13 (2013).

16. Raven, K. E. et al. Complex Routes of Nosocomial Vancomycin-Resistant Enterococcus faecium Transmission Revealed by Genome Sequencing. Clin. Infect. Dis. Off. Publ. Infect. Dis. Soc. Am. 64, 886–893 (2017).

17. Brodrick, H. J. et al. Whole-genome sequencing reveals transmission of vancomycin-resistant Enterococcus faecium in a healthcare network. Genome Med. 8, 4 (2016).

18. Gouliouris, T. et al. Quantifying acquisition and transmission of Enterococcus faecium using genomic surveillance. Nat. Microbiol. 1–9 (2020) doi:10.1038/s41564-020-00806-7.

19. Moradigaravand, D. et al. Within-host evolution of Enterococcus faecium during longitudinal carriage and transition to bloodstream infection in immunocompromised patients. Genome Med. 9, 119 (2017).

20. Ubeda, C. et al. Vancomycin-resistant *Enterococcus* domination of intestinal microbiota is enabled by antibiotic treatment in mice and precedes bloodstream invasion in humans. J. Clin. Invest. 120, 4332–4341 (2010).

21. Bayjanov, J. R. et al. Enterococcus faecium genome dynamics during long-term asymptomatic patient gut colonization. Microb. Genomics (2019) doi:10.1099/mgen.0.000277.

22. Hall, J. P. J., Brockhurst, M. A. & Harrison, E. Sampling the mobile gene pool: innovation via horizontal gene transfer in bacteria. Philos. Trans. R. Soc. B Biol. Sci. 372, (2017).

23. Lebreton, F. et al. Tracing the Enterococci from Paleozoic Origins to the Hospital. Cell 169, 849–861.e13 (2017).

24. Arredondo-Alonso, S. et al. Plasmids Shaped the Recent Emergence of the Major Nosocomial Pathogen Enterococcus faecium. mBio 11, (2020).

25. Heikens, E. et al. Contribution of the Enterococcal Surface Protein Esp to pathogenesis of Enterococcus faecium endocarditis. Microbes Infect. Inst. Pasteur 13, 1185–1190 (2011).

26. Kim, E. B. & Marco, M. L. Nonclinical and Clinical Enterococcus faecium Strains, but Not Enterococcus faecalis Strains, Have Distinct Structural and Functional Genomic Features. Appl. Environ. Microbiol. 80, 154–165 (2014).

27. Zhang, X. et al. Identification of a Genetic Determinant in Clinical Enterococcus faecium Strains That Contributes to Intestinal Colonization During Antibiotic Treatment. J. Infect. Dis. 207, 1780–1786 (2013).

28. Deutscher, J., Francke, C. & Postma, P. W. How Phosphotransferase System-Related Protein Phosphorylation Regulates Carbohydrate Metabolism in Bacteria. Microbiol. Mol. Biol. Rev. 70, 939–1031 (2006).

29. Falgenhauer, L. et al. Near-ubiquitous presence of a vancomycin-resistant Enterococcus faecium ST117/CT71/vanB -clone in the Rhine-Main metropolitan area of Germany. Antimicrob. Resist. Infect. Control 8, 128 (2019).

30. Arias, C. A. et al. Genetic Basis for In Vivo Daptomycin Resistance in Enterococci. N. Engl. J. Med. 365, 892–900 (2011).

31. Tran, T. T., Munita, J. M. & Arias, C. A. Mechanisms of Drug Resistance: Daptomycin Resistance. Ann. N. Y. Acad. Sci. 1354, 32–53 (2015).

32. Wang, G. et al. Evolution and mutations predisposing to daptomycin resistance in vancomycin-resistant Enterococcus faecium ST736 strains. PLoS ONE 13, (2018).

33. Bender, J. K. et al. Update on prevalence and mechanisms of resistance to linezolid, tigecycline and daptomycin in enterococci in Europe: Towards a common nomenclature. Drug Resist. Updat. Rev. Comment. Antimicrob. Anticancer Chemother. 40, 25–39 (2018).

34. Bodley, J. W., Zieve, F. J., Lin, L. & Zieve, S. T. Formation of the ribosome-G factor-GDP complex in the presence of fusidic acid. Biochem. Biophys. Res. Commun. 37, 437–443 (1969).

35. Nagaev, I., Björkman, J., Andersson, D. I. & Hughes, D. Biological cost and compensatory evolution in fusidic acid-resistant Staphylococcus aureus. Mol. Microbiol. 40, 433–439 (2001).

36. Wick, R. R., Schultz, M. B., Zobel, J. & Holt, K. E. Bandage: interactive visualization of de novo genome assemblies. Bioinformatics 31, 3350–3352 (2015).

37. Wick, R. R., Judd, L. M., Gorrie, C. L. & Holt, K. E. Unicycler: Resolving bacterial genome assemblies from short and long sequencing reads. PLoS Comput. Biol. 13, (2017).

38. Hashimoto, Y. et al. First Report of the Local Spread of Vancomycin-Resistant Enterococci Ascribed to the Interspecies Transmission of a vanA Gene Cluster-Carrying Linear Plasmid. mSphere 5, (2020).

39. Hashimoto, Y. et al. Novel Multidrug-Resistant Enterococcal Mobile Linear Plasmid pELF1 Encoding vanA and vanM Gene Clusters From a Japanese Vancomycin-Resistant Enterococci Isolate. Front. Microbiol. 10, (2019).

40. Harmer, C. J. & Hall, R. M. IS26 Family Members IS257 and IS1216 Also Form Cointegrates by Copy-In and Targeted Conservative Routes. mSphere 5, (2020).

41. Harmer, C. J., Moran, R. A. & Hall, R. M. Movement of IS26-Associated Antibiotic Resistance Genes Occurs via a Translocatable Unit That Includes a Single IS26 and Preferentially Inserts Adjacent to Another IS26. mBio 5, (2014).

42. Carattoli, A. et al. In Silico Detection and Typing of Plasmids using PlasmidFinder and Plasmid Multilocus Sequence Typing. Antimicrob. Agents Chemother. 58, 3895–3903 (2014).

43. Aziz, R. K. et al. The RAST Server: Rapid Annotations using Subsystems Technology. BMC Genomics 9, 75 (2008).

44. Clewell, D. B. et al. Extrachromosomal and Mobile Elements in Enterococci: Transmission, Maintenance, and Epidemiology. in Enterococci: From Commensals to Leading Causes of Drug Resistant Infection (eds. Gilmore, M. S., Clewell, D. B., Ike, Y. & Shankar, N.) (Massachusetts Eye and Ear Infirmary, 2014).

45. Sullivan, M. J., Petty, N. K. & Beatson, S. A. Easyfig: a genome comparison visualizer. Bioinforma. Oxf. Engl. 27, 1009–1010 (2011).

46. Sun, L. et al. Tandem amplification of the vanM gene cluster drives vancomycin resistance in vancomycin-variable enterococci. J. Antimicrob. Chemother. 75, 283–291 (2020).

47. Zhang, X., Vrijenhoek, J. E. P., Bonten, M. J. M., Willems, R. J. L. & Schaik, W. van. A genetic element present on megaplasmids allows Enterococcus faecium to use raffinose as carbon source. Environ. Microbiol. 13, 518–528 (2011).

48. Van Goethem, N. et al. Status and potential of bacterial genomics for public health practice: a scoping review. Implement. Sci. 14, 79 (2019).

49. Pinholt, M. et al. Multiple hospital outbreaks of vanA Enterococcus faecium in Denmark, 2012–13, investigated by WGS, MLST and PFGE. J. Antimicrob. Chemother. 70, 2474–2482 (2015).

50. Liese, J. et al. Expansion of Vancomycin-Resistant Enterococcus faecium in an Academic Tertiary Hospital in Southwest Germany: a Large-Scale Whole-Genome-Based Outbreak Investigation. Antimicrob. Agents Chemother. 63, (2019).

51. Weber, A., Maechler, F., Schwab, F., Gastmeier, P. & Kola, A. Increase of vancomycin-resistant Enterococcus faecium strain type ST117 CT71 at Charité - Universitätsmedizin Berlin, 2008 to 2018. Antimicrob. Resist. Infect. Control 9, 109 (2020).

52. Xanthopoulou, K. et al. Vancomycin-resistant Enterococcus faecium colonizing patients on hospital admission in Germany: prevalence and molecular epidemiology. J. Antimicrob. Chemother. 75, 2743–2751 (2020).

53. Neely, A. N. & Maley, M. P. Survival of Enterococci and Staphylococci on Hospital Fabrics and Plastic. J. Clin. Microbiol. 38, 724–726 (2000).

54. Kinnear, C. L. et al. Daptomycin treatment impacts resistance in off-target populations of vancomycin-resistant Enterococcus faecium. PLOS Biol. 18, e3000987 (2020).

55. Egli, A. et al. Association of daptomycin use with resistance development in Enterococcus faecium bacteraemia—a 7-year individual and population-based analysis. Clin. Microbiol. Infect. 23, 118.e1–118.e7 (2017).

56. Lellek, H. et al. Emergence of daptomycin non-susceptibility in colonizing vancomycin-resistant Enterococcus faecium isolates during daptomycin therapy. Int. J. Med. Microbiol. 305, 902–909 (2015).

57. Storm, J. C., Diekema, D. J., Kroeger, J. S., Johnson, S. J. & Johannsson, B. Daptomycin exposure precedes infection and/or colonization with daptomycin non-susceptible enterococcus. Antimicrob. Resist. Infect. Control 1, 19 (2012).

58. Chacko, K. I. et al. Genetic Basis of Emerging Vancomycin, Linezolid, and Daptomycin Heteroresistance in a Case of Persistent Enterococcus faecium Bacteremia. Antimicrob. Agents Chemother. 62, (2018).

59. Wang, G. et al. Identification of a Novel Clone, ST736, among Enterococcus faecium Clinical Isolates and Its Association with Daptomycin Nonsusceptibility. Antimicrob. Agents Chemother. 58, 4848–4854 (2014).

60. Pultz, N. J., Hoskins, L. C. & Donskey, C. J. Vancomycin-resistant Enterococci may obtain nutritional support by scavenging carbohydrate fragments generated during mucin degradation by the anaerobic microbiota of the colon. Microb. Drug Resist. Larchmt. N 12, 63–67 (2006).

61. Hinnebusch, J. & Tilly, K. Linear plasmids and chromosomes in bacteria. Mol. Microbiol. 10, 917–922 (1993).

62. Meinhardt, F., Schaffrath, R. & Larsen, M. Microbial linear plasmids. Appl. Microbiol. Biotechnol. 47, 329–336 (1997).

63. Arredondo-Alonso, S. et al. mlplasmids: a user-friendly tool to predict plasmid- and chromosome-derived sequences for single species. Microb. Genomics 4, e000224 (2018).

64. Arredondo-Alonso, S., Top, J., Corander, J., Willems, R. J. L. & Schürch, A. C. Mode and dynamics of vanA-type vancomycin resistance dissemination in Dutch hospitals. Genome Med. 13, 9 (2021).

65. Evans, D. R. et al. Systematic detection of horizontal gene transfer across genera among multidrug-resistant bacteria in a single hospital. eLife 9, e53886 (2020).

66. León-Sampedro, R. et al. Pervasive transmission of a carbapenem resistance plasmid in the gut microbiota of hospitalized patients. Nat. Microbiol. 1–11 (2021) doi:10.1038/s41564-021-00879-y.

67. Bolger, A. M., Lohse, M. & Usadel, B. Trimmomatic: a flexible trimmer for Illumina sequence data. Bioinformatics 30, 2114–2120 (2014).

68. Bankevich, A. et al. SPAdes: A New Genome Assembly Algorithm and Its Applications to Single-Cell Sequencing. J. Comput. Biol. 19, 455–477 (2012).

69. Gurevich, A., Saveliev, V., Vyahhi, N. & Tesler, G. QUAST: quality assessment tool for genome assemblies. Bioinformatics 29, 1072–1075 (2013).

70. Seemann, T. Prokka: rapid prokaryotic genome annotation. Bioinformatics 30, 2068–2069 (2014).

71. Page, A. J. et al. Roary: rapid large-scale prokaryote pan genome analysis. Bioinformatics 31, 3691–3693 (2015).

72. Price, M. N., Dehal, P. S. & Arkin, A. P. FastTree 2 – Approximately Maximum-Likelihood Trees for Large Alignments. PLoS ONE 5, (2010).

73. Croucher, N. J. et al. Rapid phylogenetic analysis of large samples of recombinant bacterial whole genome sequences using Gubbins. Nucleic Acids Res. 43, e15 (2015).

74. Page, A. J. et al. SNP-sites: rapid efficient extraction of SNPs from multi-FASTA alignments. Microb. Genomics 2, e000056 (2016).

75. Rambaut, A., Lam, T. T., Max Carvalho, L. & Pybus, O. G. Exploring the temporal structure of heterochronous sequences using TempEst (formerly Path-O-Gen). Virus Evol. 2, (2016).

76. Lam, M. M. C. et al. Comparative Analysis of the First Complete Enterococcus faecium Genome. J. Bacteriol. 194, 2334–2341 (2012).

77. Bouckaert, R. et al. BEAST 2: A Software Platform for Bayesian Evolutionary Analysis. PLOS Comput. Biol. 10, e1003537 (2014).

78. Stadler, T., Kühnert, D., Bonhoeffer, S. & Drummond, A. J. Birth-death skyline plot reveals temporal changes of epidemic spread in HIV and hepatitis C virus (HCV). Proc. Natl. Acad. Sci. U. S. A. 110, 228–233 (2013).

79. Rambaut, A., Drummond, A. J., Xie, D., Baele, G. & Suchard, M. A. Posterior Summarization in Bayesian Phylogenetics Using Tracer 1.7. Syst. Biol. 67, 901–904 (2018).

80. Inouye, M. et al. SRST2: Rapid genomic surveillance for public health and hospital microbiology labs. Genome Med. 6, 90 (2014).

81. Arthur, M., Molinas, C., Depardieu, F. & Courvalin, P. Characterization of Tn1546, a Tn3-related transposon conferring glycopeptide resistance by synthesis of depsipeptide peptidoglycan precursors in Enterococcus faecium BM4147. J. Bacteriol. 175, 117–127 (1993).

82. Wardal, E. et al. Diversity of plasmids and Tn1546-type transposons among VanA Enterococcus faecium in Poland. Eur. J. Clin. Microbiol. Infect. Dis. 36, 313–328 (2017).

83. Li, H. Aligning sequence reads, clone sequences and assembly contigs with BWA-MEM. ArXiv13033997 Q-Bio (2013).

84. Li, H. et al. The Sequence Alignment/Map format and SAMtools. Bioinforma. Oxf. Engl. 25, 2078–2079 (2009).

85. Siguier, P., Perochon, J., Lestrade, L., Mahillon, J. & Chandler, M. ISfinder: the reference centre for bacterial insertion sequences. Nucleic Acids Res. 34, D32–36 (2006).

86. Robinson, J. T. et al. Integrative genomics viewer. Nat. Biotechnol. 29, 24–26 (2011).

87. Lawrence, M. et al. Software for Computing and Annotating Genomic Ranges. PLOS Comput. Biol. 9, e1003118 (2013).

88. Guy, L., Kultima, J. R. & Andersson, S. G. E. genoPlotR: comparative gene and genome visualization in R. Bioinforma. Oxf. Engl. 26, 2334–2335 (2010).

89. Freitas, A. R. et al. Global Spread of the hylEfm Colonization-Virulence Gene in Megaplasmids of the Enterococcus faecium CC17 Polyclonal Subcluster. Antimicrob. Agents Chemother. 54, 2660–2665 (2010).

90. Zuker, M. Mfold web server for nucleic acid folding and hybridization prediction. Nucleic Acids Res. 31, 3406–3415 (2003).

91. Arndt, D. et al. PHASTER: a better, faster version of the PHAST phage search tool. Nucleic Acids Res. 44, W16–W21 (2016).

92. Garcillán-Barcia, M. P., Redondo-Salvo, S., Vielva, L. & de la Cruz, F. MOBscan: Automated Annotation of MOB Relaxases. Methods Mol. Biol. Clifton NJ 2075, 295–308 (2020).

93. Huerta-Cepas, J. et al. eggNOG 5.0: a hierarchical, functionally and phylogenetically annotated orthology resource based on 5090 organisms and 2502 viruses. Nucleic Acids Res. 47, D309–D314 (2019).

94. Poyart, C. & Trieu-Cuot, P. Heterogeneric conjugal transfer of the pheromone-responsive plasmid pIP964 (IncHlyI) of Enterococcus faecalis in the apparent absence of pheromone induction. FEMS Microbiol. Lett. 122, 173–179 (1994).

95. Werner, G. et al. Host range of enterococcal vanA plasmids among Gram-positive intestinal bacteria. J. Antimicrob. Chemother. 66, 273–282 (2011).

