## Supplemental Information for "Genomic surveillance of vancomycin-resistant *Enterococcus faecium* reveals spread of a linear plasmid conferring a nutrient utilization advantage"

### Supplemental Figures

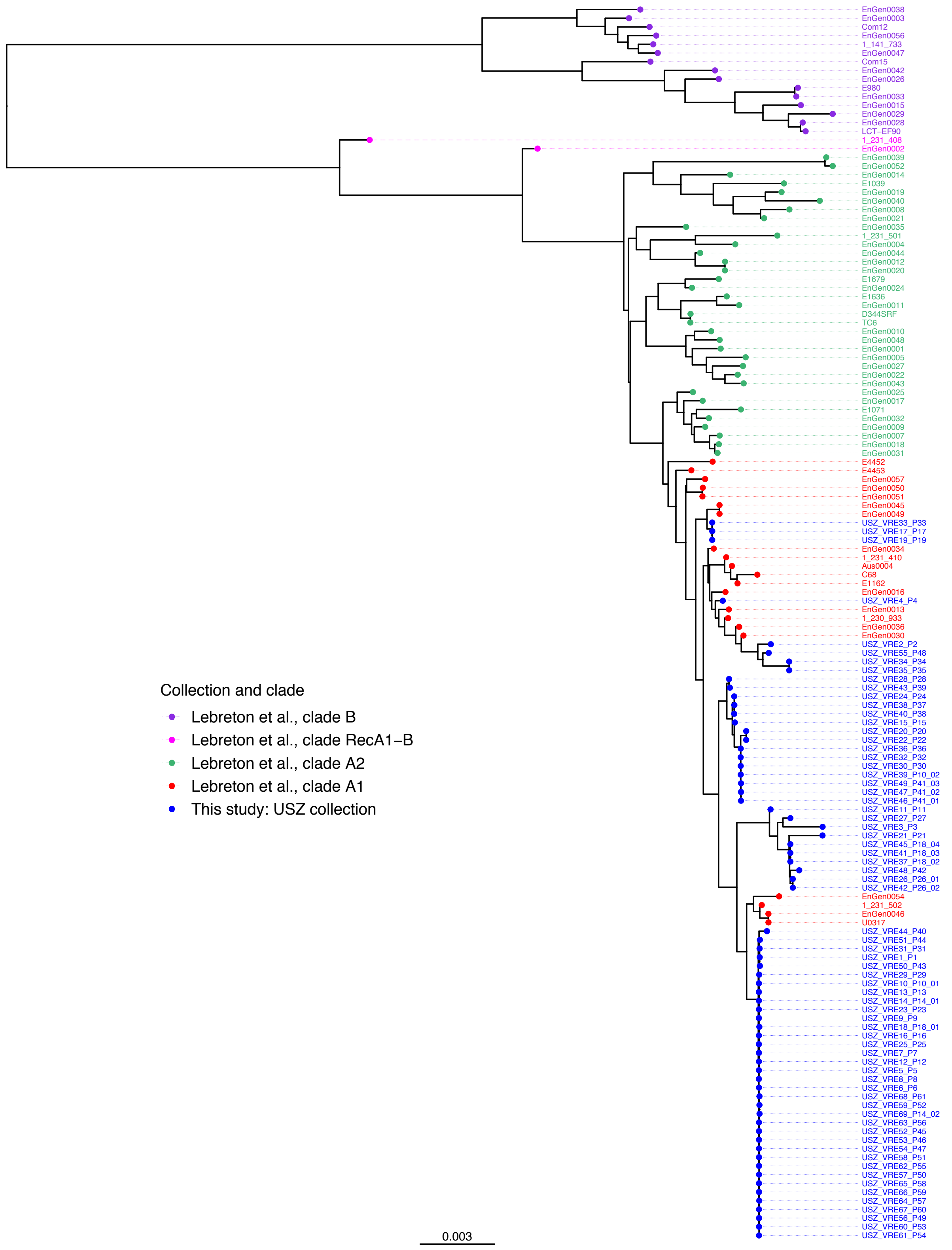

**Figure S1. Maximum-likelihood phylogeny with all reference isolates**. This tree was built using the 1,131,323 bp alignment of 1,224 core genes of the full collection (73 isolates from Lebreton et al. [1] and the 69 isolates of this study) and was midpoint-rooted. Colors reflect study and clade affiliation.

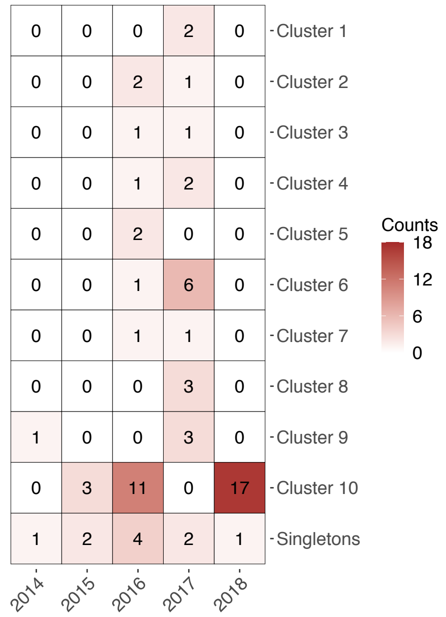

**Figure S2. Number of isolates of each clone sampled per year**. This map merges the information enclosed in Fig. 1A and 1B: sampling timing (year of isolation) and genetic relatedness (i.e., phylogenomic clusters). Values indicate the number of isolates of a given cluster (row) which were isolated in a given year (column). Color intensity reflects the magnitude of the corresponding value.

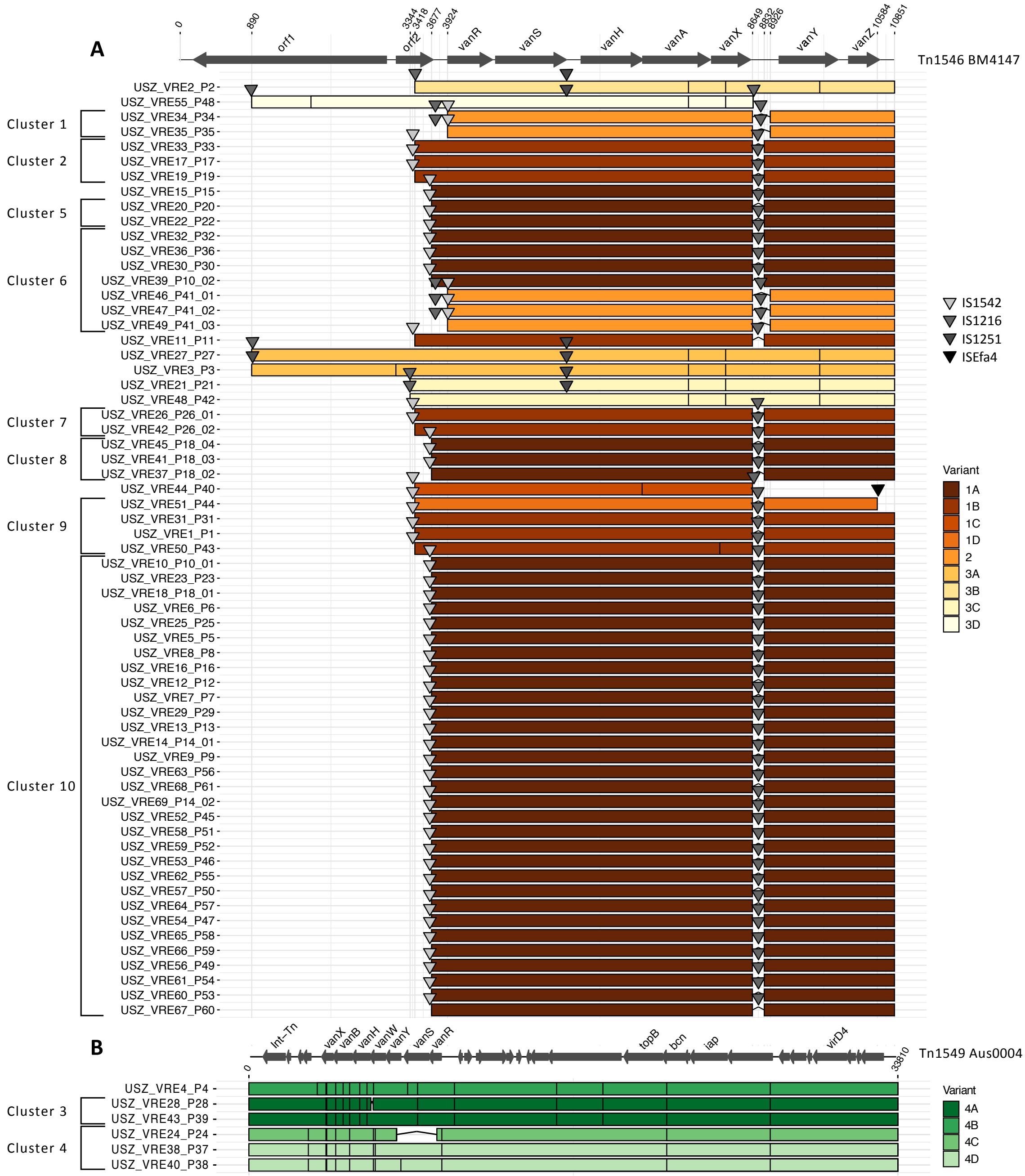

**Figure S3. Vancomycin transposon variability**. Alignment to Tn1546 of B4147 [2,3] for isolates with *vanA* genes (**A**) and to Tn1549 of Aus0004 [4] for isolates with *vanB* genes (**B**). Within each panel, isolates are ordered based on their position in the phylogenetic tree of Fig. 1B. The x-axis shows the position on the reference in bp. Note that the scale is different for Tn1546 and Tn1549. Based on genotype (*vanA* or *vanB*) and insertion sequences incorporated, four transposon types (labeled 1-4) were defined, and based on similarity with the references, visualized here for each isolate, these were subdivided into subtypes (labelled A-D). The horizontal segments indicate covered areas, vertical black lines represent SNPs, and the triangles represent insertion sequences. Distinct variants of the vancomycin transposon were each attributed a color (brown shades for *vanA*, type 1-3 and green shades for *vanB*, type 4), taken up in Fig. 1B. All *vanA* transposon types exhibited full or partial loss of *orf1* and *orf2*, due to the incorporation of insertion sequences. In contrast, all *vanB* transposon types covered the 33,810 bp Tn1549 reference, except for the loss of the *vanR* and *vanS* genes by one isolate (USZ_VRE24_P24).

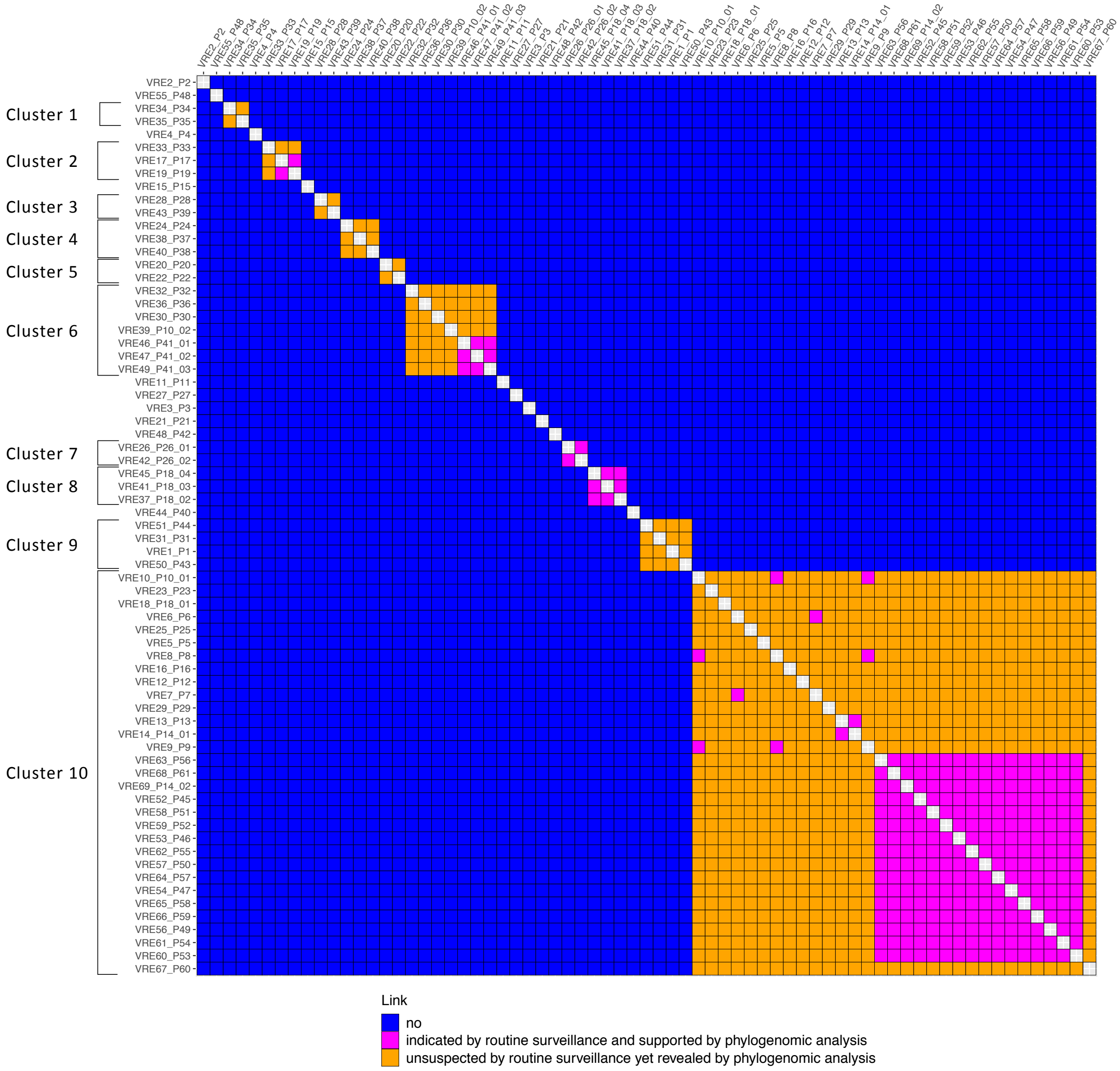

**Figure S4. Pairwise matrix of transmission networks**. The links indicated by routine surveillance (based on epidemiological criteria and traditional typing), which were all supported by the phylogenomic analysis, are highlighted in magenta. The links between isolates revealed by the phylogenomic analysis are highlighted in orange. The isolates are ordered based on their position in the phylogenetic tree of Fig. 1B.

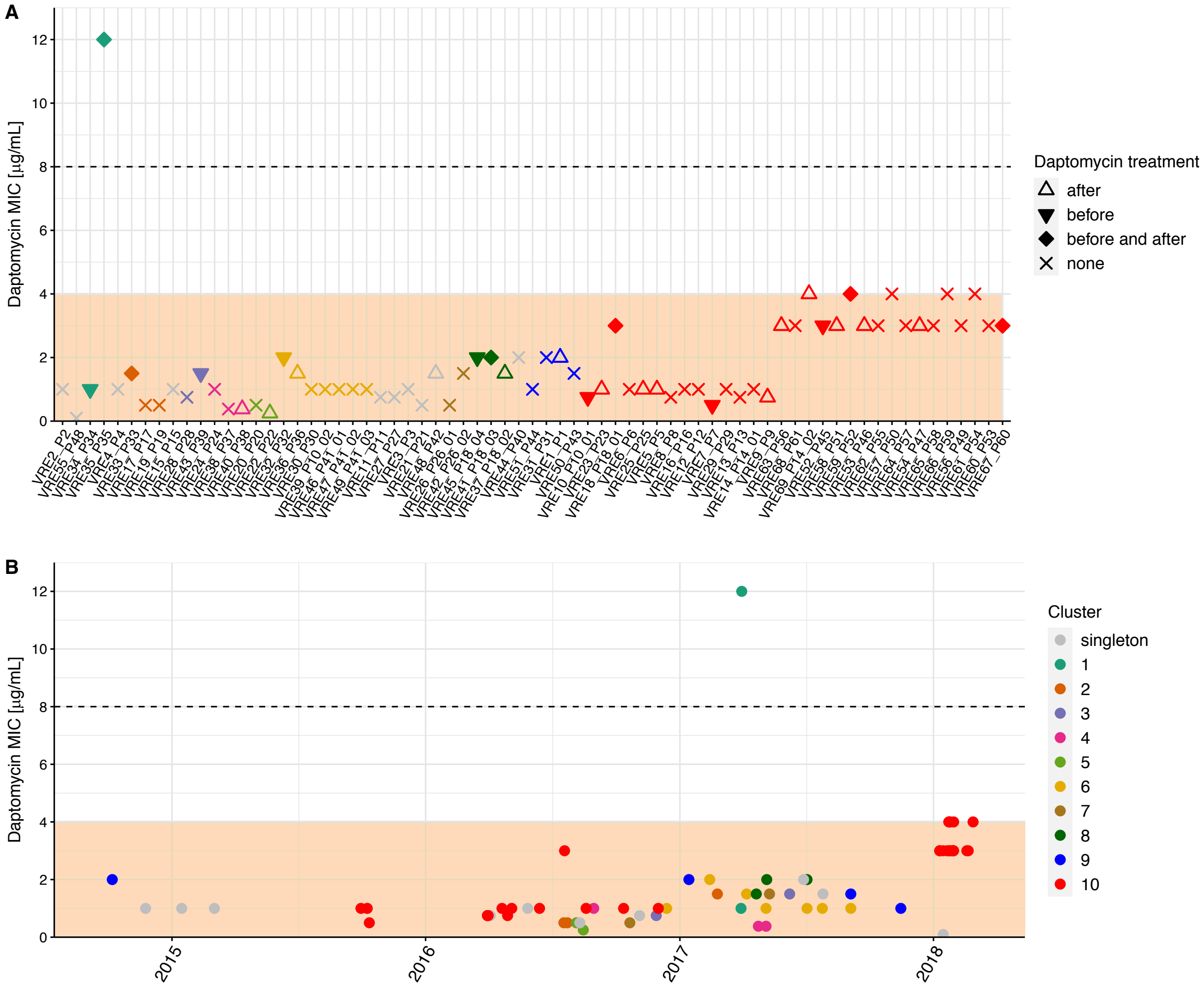

**Figure S5. Daptomycin MIC**. *E. faecium* strains with daptomycin MIC falling in the range ≤ 4 µg/mL (color highlighted area) are considered susceptible-dose-dependent while those with a MIC ≥ the breakpoint of 8 µg/mL (shown with a dashed line) are considered non-susceptible, according to the CLSI guidelines of 2019 [5]. The same data are presented in **A** and **B**. In both panels, color of data points reflects the clusters. **A** highlights changes in the MIC within clusters. To this scope, isolates are presented in the order in which they appear in the phylogenetic tree of Fig. 1B. Shape of the data points illustrates whether the patient from whom the isolate was sampled had been treated with daptomycin after, before, before and after or not at all. The two cases including "before" are highlighted with a closed shape, indicating that the isolate was derived from a strain which had experienced antibiotic pressure within host. **B** shows the global temporal evolution.

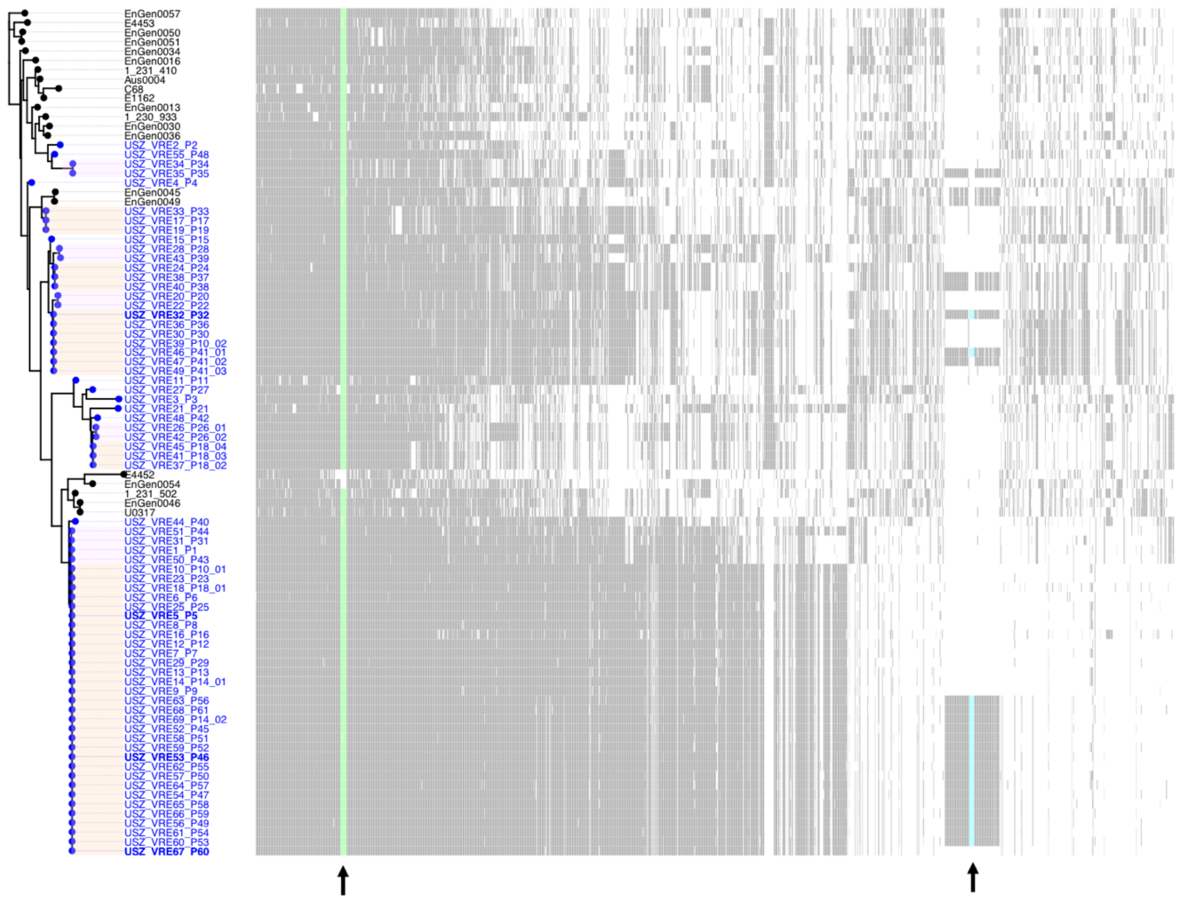

**Figure S6. Map of accessory genes**. The maximum-likelihood phylogenetic tree of Fig. 1B is shown on the left and a map of accessory genes, i.e., genes present in >14% and <99% genomes of the collection, is shown on the right. The pattern of presence/absence is shown with vertical segments (black/white respectively) for each row, which corresponds to one isolate (tree tip). Arrows at the bottom of this map point towards specific gene clusters. These represent two carbohydrate utilization operons: the one including PTS$^{clin}$, the mannose family PTS previously shown to be enriched in clinical isolates [6] (highlighted in green) and the cargo genes of pELF_USZ, which included a mannose family PTS as well (highlighted in cyan).

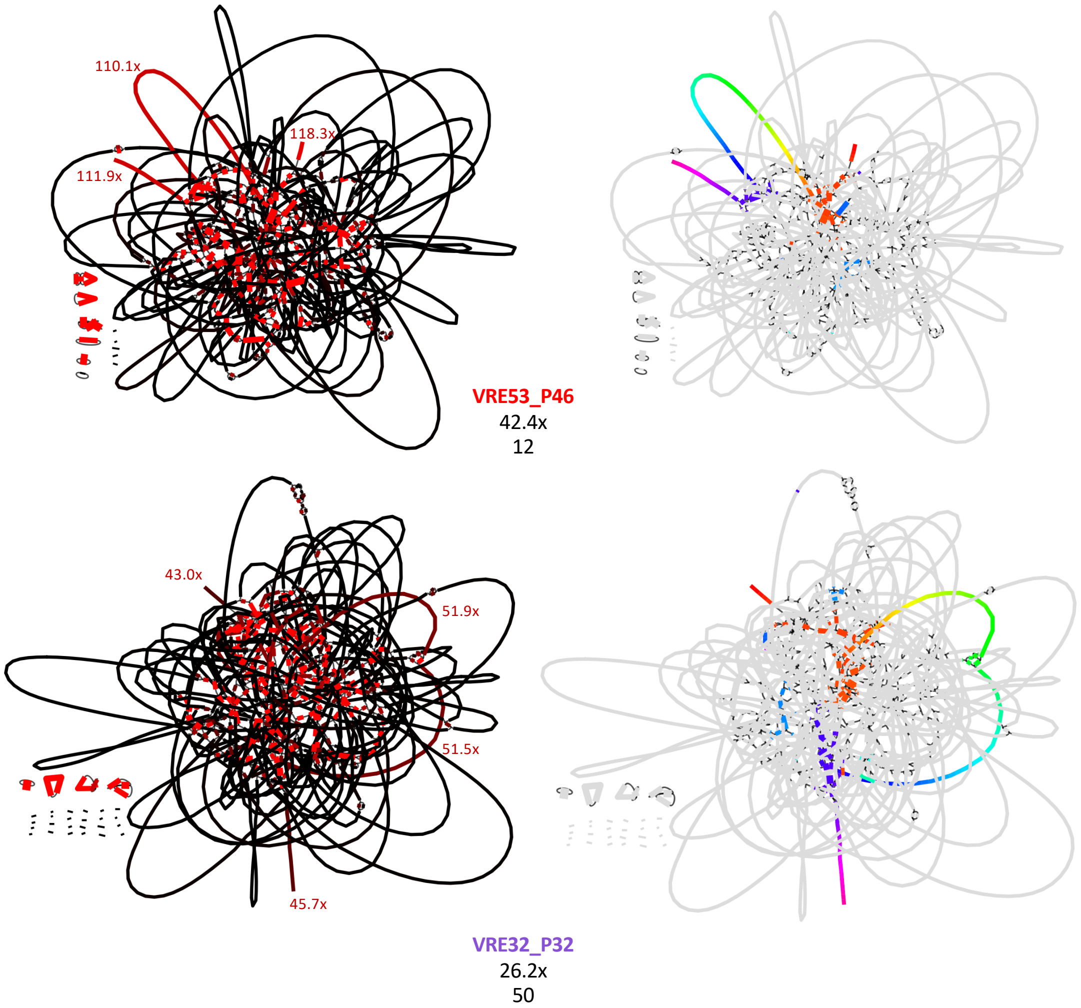

**Figure S7. Short-read assembly graphs and pELF_USZ**. Visualization with Bandage [7] of the assembly graphs of VRE53_P46 and VRE32_P32 obtained with SPAdes [8]. These graphs consist of the contigs and their connections to other contigs, ambiguous in repetitive regions too long to be spanned by short-read sequences. For each isolate, the same assembly graph is shown twice: on the left, color reflects the coverage (depth), ranging from black for the median coverage, to light red for highest coverage; on the right, the rainbow-colored contigs are BLAST hits of pELF_USZ, to visualize its localization within the graph.

The median coverage as well as the total number of dead ends of each assembly is displayed below the label. The coverage of the main three, respectively four, contigs corresponding to pELF_USZ is annotated on the left graph. Note that this coverage corresponds to two to three-fold the median coverage. Moreover, the contigs corresponding to the extremities of pELF_USZ are the only contigs longer than 600 bp that exhibit dead ends.

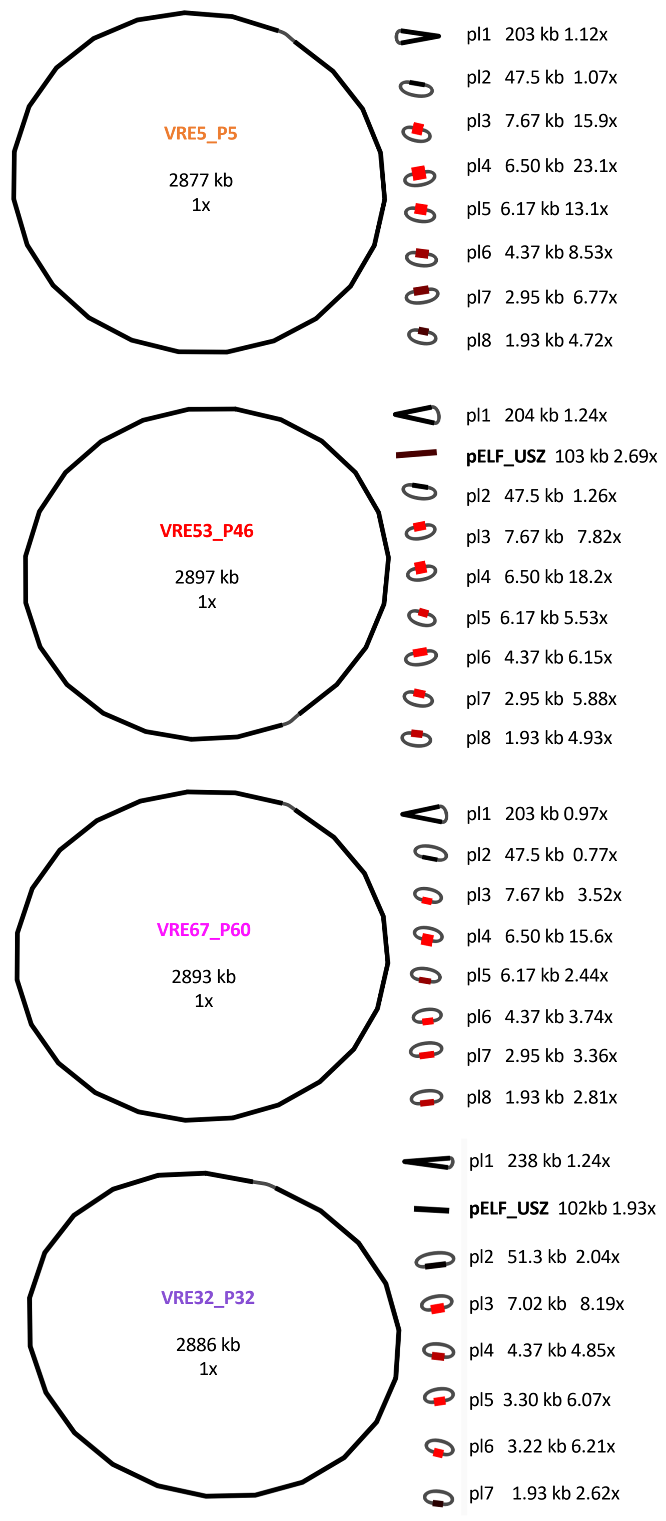

**Figure S8. Complete assemblies of four selected isolates**. Visualization with Bandage [7] of the hybrid assemblies obtained by scaffolding short- with long-reads using Unicycler [9]. The length of each sequence is given in kilobase-pairs (kb) and is followed by the coverage (depth) with respect to the chromosome coverage (set to 1x). Color, ranging from black for the chromosome to light red reflects the coverage and grey strings indicate circularization.

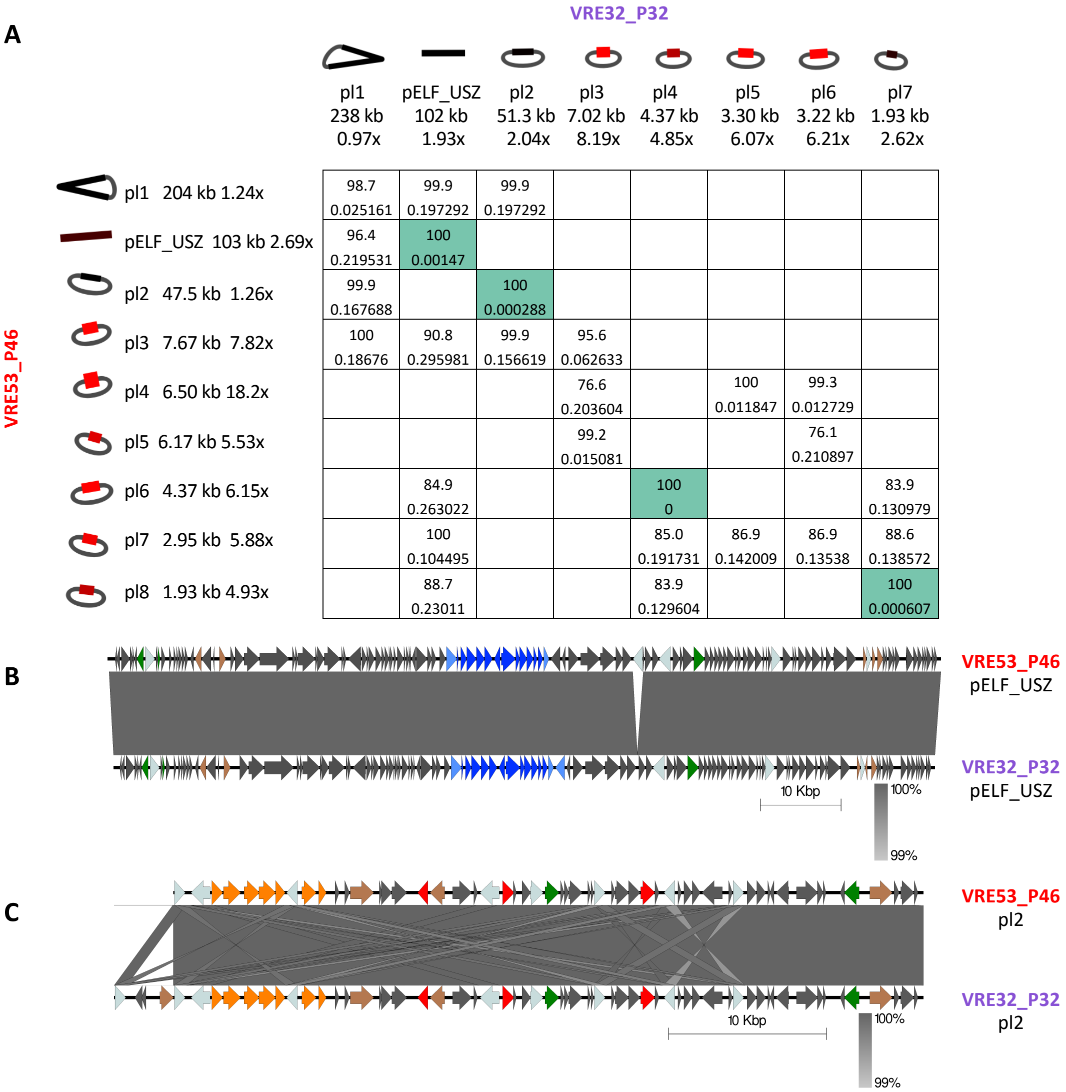

| VRE53_P46 \ VRE32_P32 | pl1 | pELF_USZ | pl2 | pl3 | pl4 | pl5 | pl6 | pl7 |
|---|---|---|---|---|---|---|---|---|
| pl1 | 98.7 0.025161 | 99.9 0.197292 | 99.9 0.197292 | | | | | |
| pELF_USZ | 96.4 0.219531 | 100 0.00147 | | | | | | |
| pl2 | 99.9 0.167688 | | 100 0.000288 | | | | | |
| pl3 | 100 0.18676 | 90.8 0.295981 | 99.9 0.156619 | 95.6 0.062633 | | | | |
| pl4 | | | | 76.6 0.203604 | | 100 0.011847 | 99.3 0.012729 | |
| pl5 | | | | 99.2 0.015081 | | | 76.1 0.210897 | |
| pl6 | | 84.9 0.263022 | | | 100 0 | | | 83.9 0.130979 |
| pl7 | | 100 0.104495 | | | 85.0 0.191731 | 86.9 0.142009 | 86.9 0.13538 | 88.6 0.138572 |
| pl8 | | 88.7 0.23011 | | | 83.9 0.129604 | | | 100 0.000607 |

**Figure S9. Comparison of the plasmid sequences of two isolates from distinct clones**. **A** Comparison matrix including two similarity indexes: first, the percentage Average Nucleotide Identity (ANI), which is based on alignment and thus only considers the core [10]; below, the mash distance, which relies on k-mers comparisons and thus also reflects differences in sequence length [11]. Comparisons identifying matching plasmid sequences, defined as ANI>99% and mash distance<0.01, are highlighted in green. Comparisons for which the ANI was of 0% are left blank. **B, C** Visualization of the alignments of two pairs of matching plasmid sequences: pELF_USZ (**B**) and pl2, the plasmid conferring vancomycin-resistance (**C**) produced with Easyfig [12]. A minimum blast length of 500 bp and minimum identity of 99% were considered to plot the alignment blocks. The coding sequences are shown with arrows and their color reflects their function: the *van* genes are in orange, the other resistance genes in red, the proteins with a putative role in DNA replication are in brown, those with a putative role in DNA partitioning and transfer are in green and the insertion sequences in light grey/blue. The cargo genes of pELF_USZ are highlighted in dark blue.

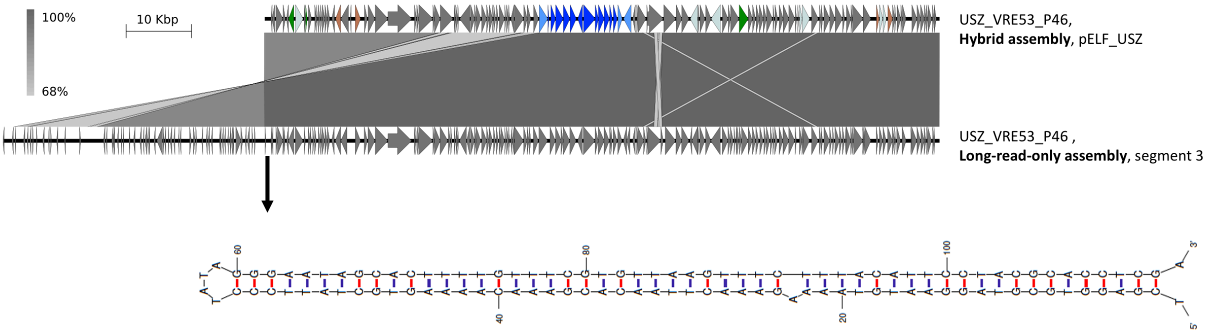

**Figure S10. Left-end hairpin structure of pELF_USZ**. Visualization of the alignment of the plasmid sequence resulting from a hybrid assembly with the corresponding segment in the long-read-only assembly, produced with Easyfig [12]. Coding sequences are shown with arrows. On pELF_USZ, their color reflects gene function, as described in the legends of Fig. 4 and Fig. S9. Color was omitted for the long-read only assembly segment because, due to the low-accuracy of nanopore reads, coding sequences were very disrupted. A minimum blast length of 500 bp was considered to plot the alignment blocks. The arrow originating from the center of the inverse repeats indicates the DNA folding of the 113 bp spanning this area, obtained with mfold [13].

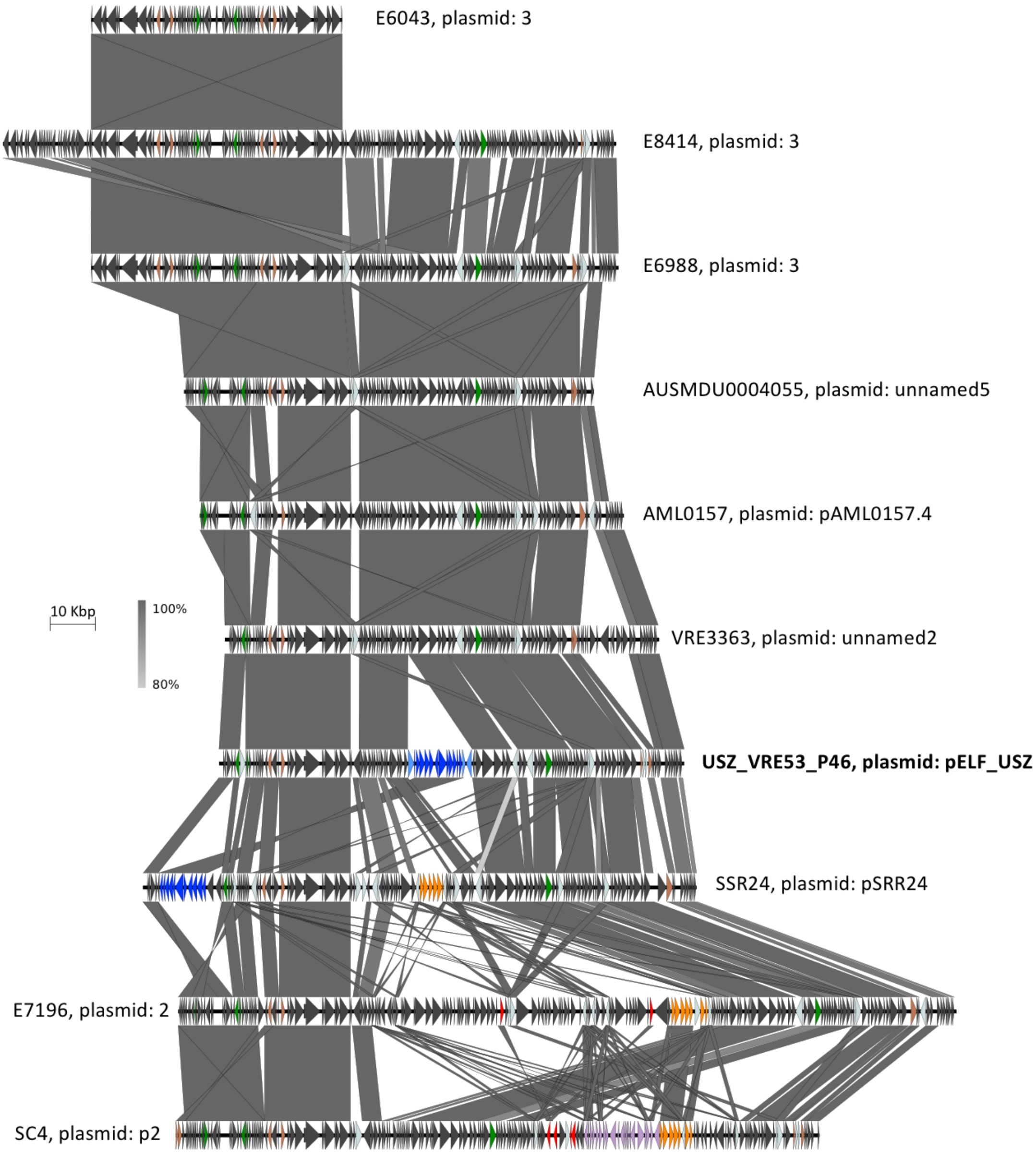

**Figure S11. Homology of pELF_USZ with other plasmid sequences**. Multiple sequence alignment visualized with Easyfig [12]. Eight of these sequences were the best hits when querying pELF_USZ against the nucleotide collection database (nr/nt), based on the maximum score, as per 04.12.2020 (Supplemental Table 6). In addition, pSRR24 in spite of not ranking in the top-8 hits, was, as per 04.12.2020, the only *E. faecium* assembly including the cargo genes of pELF_USZ, highlighted in dark blue. As in Fig. 4, S9 and S10, the *van* genes are in orange, the other resistance genes in red, the proteins with a putative role in DNA replication are in brown, those with a putative role in DNA partitioning and transfer are in green, the insertion sequences in light grey/blue and a complete prophage sequence is in purple.

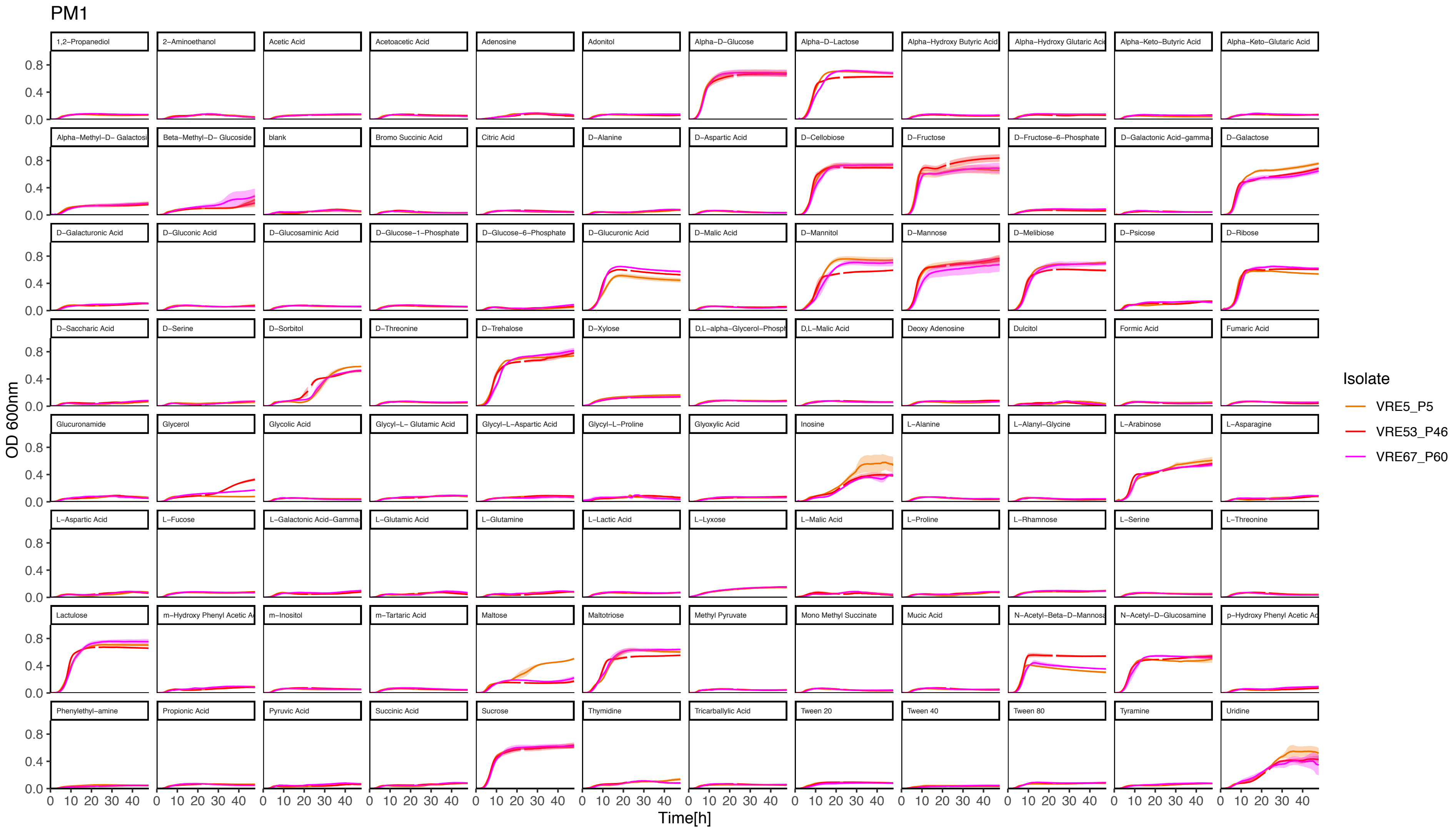

**Figure S12. Growth in M1 supplemented with 95 single carbon sources (PM1 BioLog Inc)**. The mean of three biological replicates for each representative isolate is displayed, with the standard error of mean shaded. Conditions are ordered alphabetically, blank well (only M1) is also shown (second row, third column).

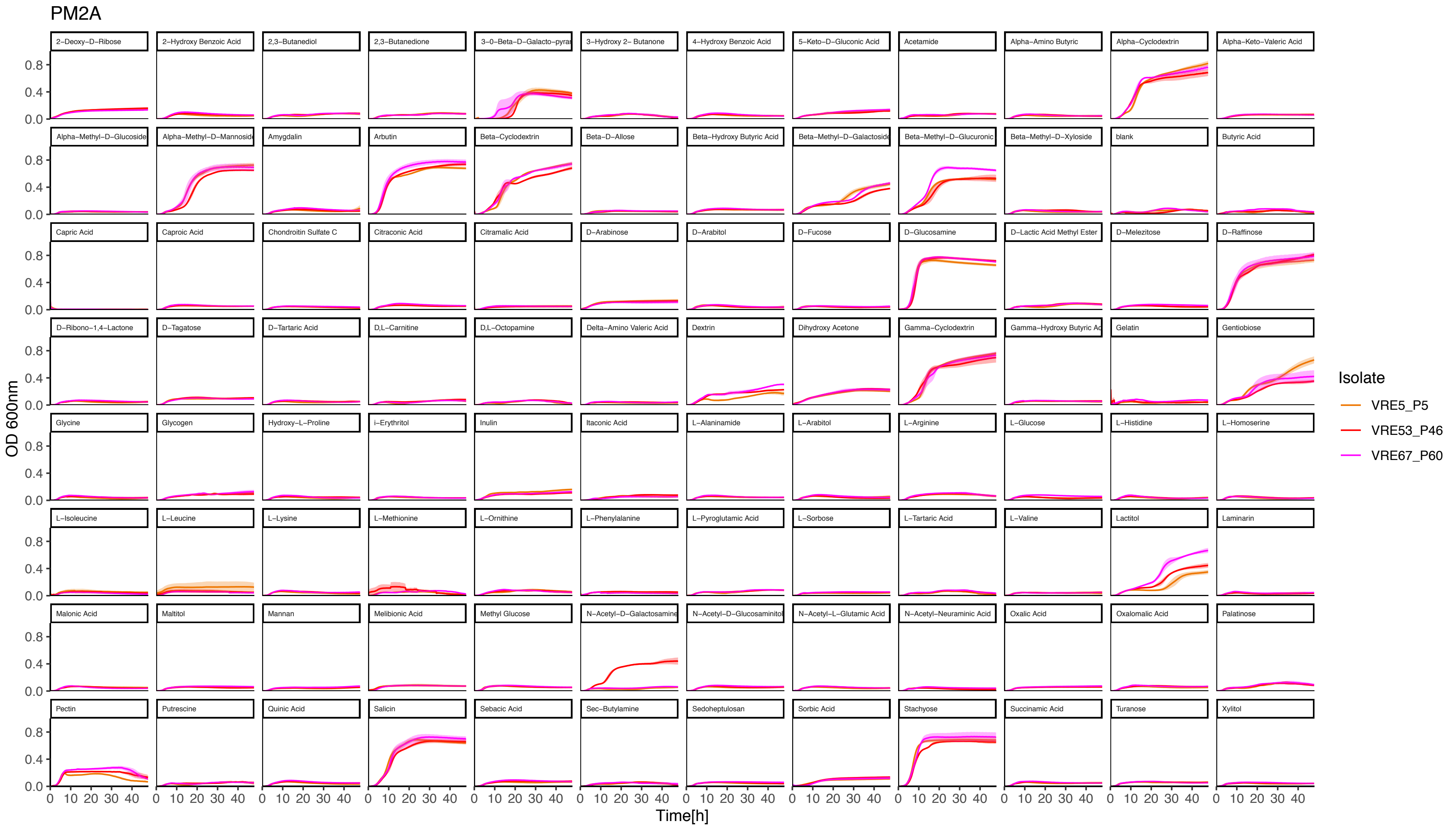

**Figure S13. Growth in M1 supplemented with 95 single carbon sources (PM2A BioLog Inc)**. The mean of three biological replicates for each representative isolate is displayed, with the standard error of mean shaded. Conditions are ordered alphabetically, blank well (only M1) is also shown (second row, eleventh column).

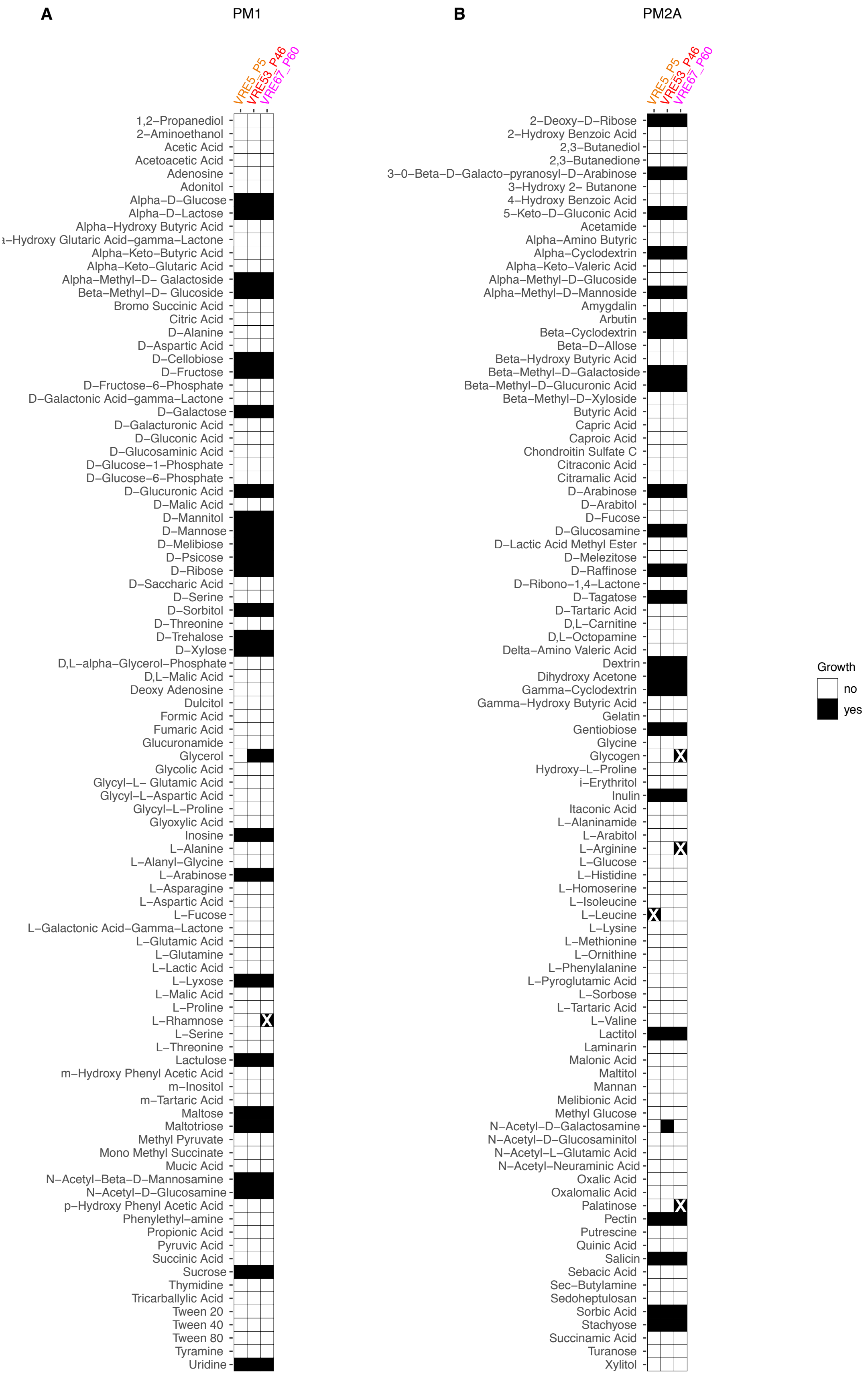

**Figure S14 Utilization of 190 single carbon sources for representative isolates of the *persistent clone*.** Conditions in which the area under the curve (AUC) was ≥ 230 were considered to be growth permissive (AUC of the blank was 133.8 ± 6.5). White crosses over a black square indicate false positives attributable to noise.

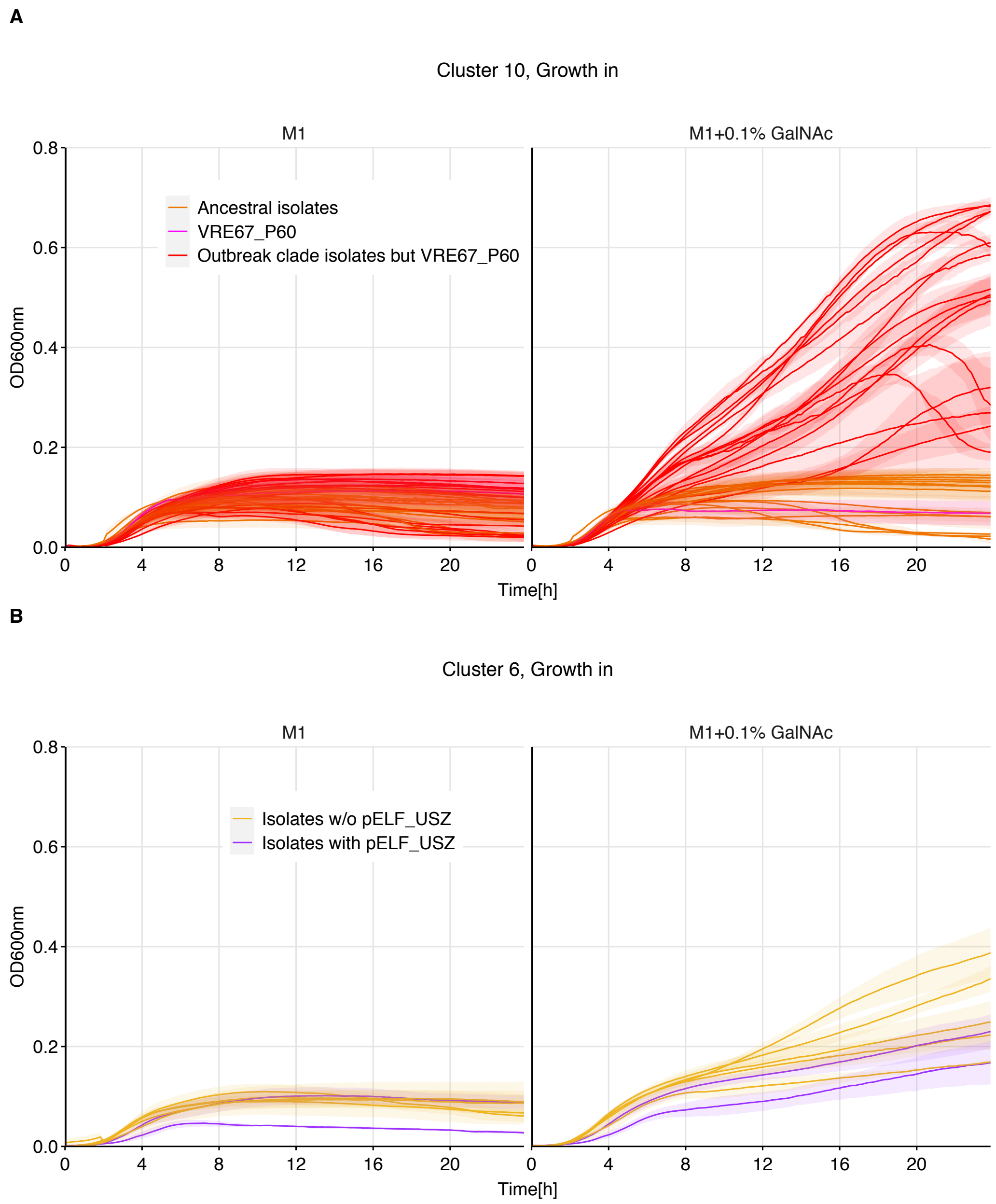

**Figure S15. Growth in M1 medium and M1 medium supplemented with 0.1% GalNAc** for the isolates of *Cluster 10* (**A**) and *Cluster 6* (**B**). In **A**, colors are the same as those introduce for Fig. 3 and 5: the *ancestral* isolates are in orange and the *outbreak clade* isolates are in red, excluding the only clade member which did not carry pELF_USZ, which is in magenta. In **B**, the isolates which carried pELF_USZ are in purple (VRE32_P32 and VRE46_P41_01), while the others are in yellow (VRE30_P30, VRE36_P36, VRE39_P10_02, VRE47_P41_02, VRE49_P41_03). The maximal OD reached during these 24h of growth are displayed in Fig. 5B and Supplemental Fig. S16.

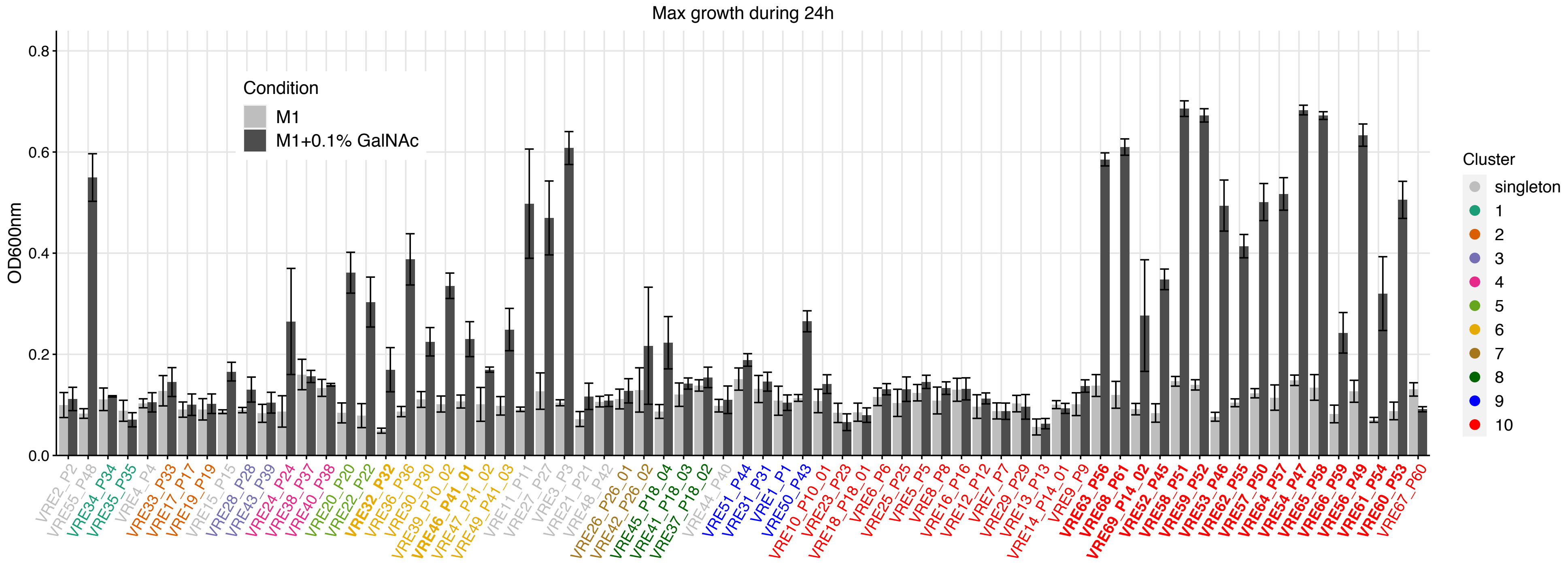

**Figure S16. GalNAc utilization by all isolates of the collection**. Maximum $OD_{600}$ reached during 24h of growth in either M1 medium only or in M1 medium supplemented with 0.1% GalNAc (N=3, mean and standard error of mean are shown). The labels of isolates harboring pELF_USZ are in bold. Isolates are ordered based on their position in the phylogenetic tree of Fig. 1B. Color of the labels reflect cluster affiliation.

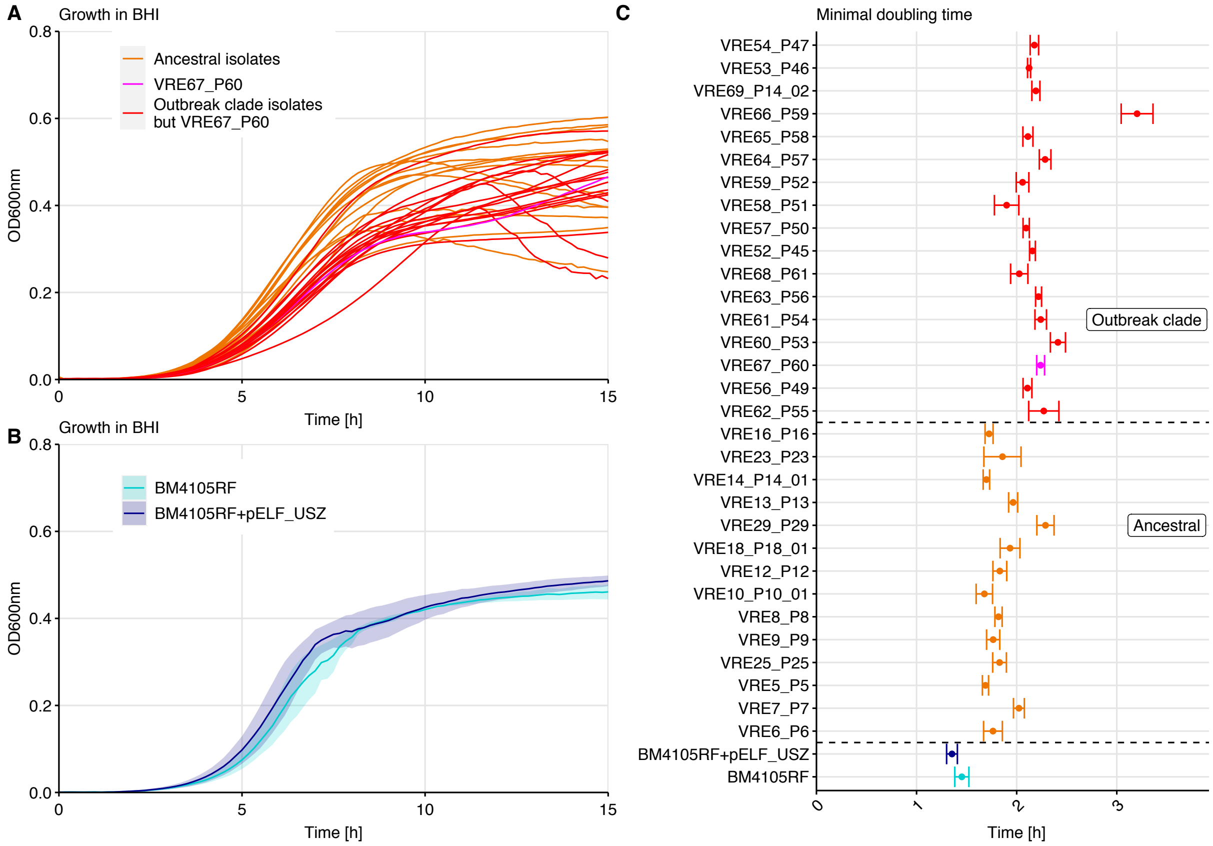

**Figure S17. Growth in liquid nutrient rich medium**. Growth curves based on optical density measurements ($OD_{600}$) every 10min during 24h for the isolates of the persistent clone (**A**); and the recipient and transconjugant (**B**). The average of three biological replicates is shown. Standard error of mean is shown with a shaded area in **B** and omitted in **A** for better visibility. **C** Minimal doubling time (MDT) of the 33 isolates was calculated based on the optical density measurements of each of the 3 biological replicates, using 1h intervals. Mean and standard error of mean are shown here. The isolates are ordered based on their position on the phylogenetic tree of Fig. 3A. The two groups, comprising *ancestral* isolates or *outbreak clade* isolates, differed significantly in their MDT, Welch's *t* test p= 7.19 e-05, with a mean of 1 h and 51 min and 2 h 13 min, respectively. The recipient (BM4105RF) and transconjugant (BM4105RF+pELF_USZ) did not differ significantly in their MDT.

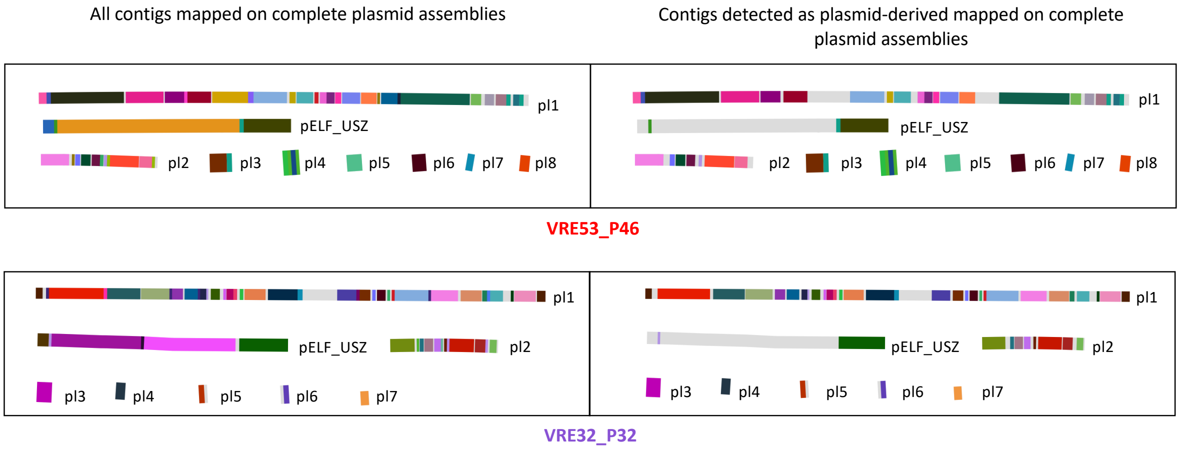

**Figure S18. Screening of short-read-assemblies-contigs with mlplasmid** [14]. Visualization with Bandage [7] of the contigs from the short-read assemblies mapped on the complete plasmid sequences. Different colors represent different contigs and the light grey represents unmatched regions. BLAST parameters: minimum query coverage of 40%, minimum identity of 99% and minimal length of 1,000 bp. On the left, all contigs are displayed, while on the right, only the contigs detected as plasmid-derived by mlplasmid are displayed. The proportion of contigs and percent of length accurately detected as plasmid-derived for each plasmid sequence are reported in the Supplemental Table S8.

#### Supplemental Tables

| Isolate | Sampling date | Sample site | Case | ST | Van genotype | Dap MIC | Dap treatment |
|---|---|---|---|---|---|---|---|
| VRE1_P1 | Oct-2014 | Tracheal swab | colonization | 203 | vanA | 2 | after |
| VRE2_P2 | Nov-2014 | Urin | colonization | 17 | vanA | 1 | - |
| VRE3_P3 | Jan-2015 | Urin | colonization | 789 | vanA | 1 | - |
| VRE4_P4 | Mar-2015 | Tracheal swab | colonization | 117 | vanB | 1 | - |
| VRE5_P5 | Sep-2015 | Superficial wound | infection | 203 | vanA | 1 | after |
| VRE6_P6 | Oct-2015 | Catheter | colonization | 203 | vanA | 1 | - |
| VRE7_P7 | Oct-2015 | Urin | colonization | 203 | vanA | 0.5 | before |
| VRE8_P8 | Mar-2016 | Perianal swab | colonization | 203 | vanA | 0.75 | - |
| VRE9_P9 | Mar-2016 | Inguinal swab | colonization | 203 | vanA | 0.75 | after |
| VRE10_P10_01 | Mar-2016 | Superficial wound | colonization | 203 | vanA | 0.75 | before |
| VRE11_P11 | Apr-2016 | Urin | infection | 80 | vanA | 0.75 | - |
| VRE12_P12 | Apr-2016 | Urin | colonization | 203 | vanA | 1 | - |
| VRE13_P13 | Apr-2016 | Urin | colonization | 203 | vanA | 0.75 | - |
| VRE14_P14_01 | May-2016 | Inguinal swab | colonization | 203 | vanA | 1 | - |
| VRE15_P15 | May-2016 | Urin | colonization | 117 | vanA | 1 | - |
| VRE16_P16 | Jun-2016 | Inguinal swab | colonization | 203 | vanA | 1 | - |
| VRE17_P17 | Jul-2016 | Urin | colonization | 192 | vanA | 0.5 | - |
| VRE18_P18_01 | Jul-2016 | Percutaneous biliary drainage | infection | 203 | vanA | 3 | before and after |
| VRE19_P19 | Jul-2016 | Rectal swab | colonization | 192 | vanA | 0.5 | - |
| VRE20_P20 | Aug-2016 | Inguinal/rectal swab | infection | 117 | vanA | 0.5 | - |
| VRE21_P21 | Aug-2016 | Inguinal/rectal swab | colonization | 992 | vanA | 0.5 | - |
| VRE22_P22 | Aug-2016 | Abdomen | infection | 117 | vanA | 0.25 | after |
| VRE23_P23 | Aug-2016 | Urin | infection | 203 | vanA | 1 | after |
| VRE24_P24 | Aug-2016 | Rectal swab | colonization | 117 | vanB | 1 | - |
| VRE25_P25 | Oct-2016 | Rectal swab | infection | 203 | vanA | 1 | after |
| VRE26_P26_01 | Oct-2016 | Inguinal swab | colonization | 80 | vanA | 0.5 | - |
| VRE27_P27 | Nov-2016 | Urin | colonization | 80 | vanA | 0.75 | - |
| VRE28_P28 | Nov-2016 | Inguinal swab | colonization | 117 | vanB | 0.75 | - |
| VRE29_P29 | Dec-2016 | Abdomen | colonization | 203 | vanA | 1 | - |
| VRE30_P30 | Dec-2016 | Urin | colonization | 117 | vanA | 1 | - |
| VRE31_P31 | Jan-2017 | Rectal swab | colonization | 203 | vanA | 2 | - |
| VRE32_P32 | Feb-2017 | Inguinal/rectal swab | colonization | 117 | vanA | 2 | before |
| VRE33_P33 | Feb-2017 | Abdomen | infection | 192 | vanA | 1.5 | before and after |
| VRE34_P34 | Mar-2017 | Rectal swab | infection | 80 | vanA | 1 | before |
| VRE35_P35 | Mar-2017 | Rectal swab | infection | 80 | vanA | 12 | before and after |
| VRE36_P36 | Apr-2017 | Deep wound | infection | 117 | vanA | 1.5 | after |
| VRE37_P18_02 | Apr-2017 | Catheter | infection | 80 | vanA | 1.5 | after |
| VRE38_P37 | Apr-2017 | Abdomen | infection | 117 | vanB | 0.38 | - |
| VRE39_P10_02 | May-2017 | Inguinal swab | colonization | 117 | vanA | 1 | - |
| VRE40_P38 | May-2017 | Abdomen | infection | 117 | vanB | 0.38 | after |
| VRE41_P18_03 | May-2017 | Blood culture | infection | 80 | vanA | 2 | before and after |

| Isolate | Sampling date | Sample site | Case | ST | Van genotype | Dap MIC | Dap treatment |
|---|---|---|---|---|---|---|---|
| VRE42_P26_02 | May-2017 | Rectal swab | colonization | 80 | vanA | 1.5 | - |
| VRE43_P39 | Jun-2017 | Rectal swab | colonization | 117 | vanB | 1.5 | before |
| VRE44_P40 | Jun-2017 | Urin | colonization | 203 | vanA | 2 | - |
| VRE45_P18_04 | Jul-2017 | Blood culture | infection | 80 | vanA | 2 | before |
| VRE46_P41_01 | Jul-2017 | Rectal swab | colonization | 117 | vanA | 1 | - |
| VRE47_P41_02 | Jul-2017 | Inguinal swab | colonization | 117 | vanA | 1 | - |
| VRE48_P42 | Jul-2017 | Inguinal swab | infection | 80 | vanA | 1.5 | after |
| VRE49_P41_03 | Sep-2017 | Inguinal swab | colonization | 117 | vanA | 1 | - |
| VRE50_P43 | Sep-2017 | Inguinal swab | colonization | 203 | vanA | 1.5 | - |
| VRE51_P44 | Nov-2017 | Inguinal swab | colonization | 203 | vanA | 1 | - |
| VRE52_P45 | Jan-2018 | Inguinal swab | infection | 203 | vanA | 3 | before |
| VRE53_P46 | Jan-2018 | Superficial wound | colonization | 203 | vanA | 3 | after |
| VRE54_P47 | Jan-2018 | Perianal swab | infection | 203 | vanA | 3 | after |
| VRE55_P48 | Jan-2018 | Inguinal swab | colonization | 1336 | vanA | 0.094 | - |
| VRE56_P49 | Jan-2018 | Urin | colonization | 203 | vanA | 3 | - |
| VRE57_P50 | Jan-2018 | Urin | colonization | 203 | vanA | 4 | - |
| VRE58_P51 | Jan-2018 | Urin | infection | 203 | vanA | 3 | after |
| VRE59_P52 | Jan-2018 | Rectal swab | colonization | 203 | vanA | 4 | before and after |
| VRE60_P53 | Jan-2018 | Rectal swab | colonization | 203 | vanA | 3 | - |
| VRE61_P54 | Jan-2018 | Pooled swab | colonization | 203 | vanA | 4 | - |
| VRE62_P55 | Jan-2018 | Rectal swab | colonization | 203 | vanA | 3 | - |
| VRE63_P56 | Jan-2018 | Blood culture | infection | 203 | vanA | 3 | after |
| VRE64_P57 | Jan-2018 | Inguinal/rectal swab | colonization | 203 | vanA | 3 | - |
| VRE65_P58 | Jan-2018 | Rectal swab | colonization | 203 | vanA | 3 | - |
| VRE66_P59 | Jan-2018 | Urin | colonization | 203 | vanA | 4 | - |
| VRE67_P60 | Feb-2018 | Superficial wound | infection | 203 | vanA | 3 | before and after |
| VRE68_P61 | Feb-2018 | Urin | colonization | 203 | vanA | 3 | - |
| VRE69_P14_02 | Feb-2018 | Urin | infection | 203 | vanA | 4 | after |

**Table S1. Metadata of the isolates**. For each isolate, the date of sampling, sampling site and whether the strain caused and infection or colonized the patient are given in the columns 2-4. The sequence type (ST) and the vancomycin (Van) genotype determined with routine molecular genotyping and confirmed *in silico* using mlst (https://github.com/tseemann/mlst) and SRST2 [15] respectively, are given in column 5-6. Finally, columns 7-8 report daptomycin (Dap) MIC as well as timing with respect to sampling of an eventual treatment with daptomycin.

| Clade | Isolate | Year of isolation | Country | ST | GenBank accession |
|---|---|---|---|---|---|
| B | EnGen0003 | 2001 | IRL | 163 | GCF_000321865.1 |
| B | Com12 | 2006 | USA | 107 | GCF_000157635.1 |
| B | EnGen0056 | 2000 | NLD | 327 | GCF_000322265.1 |
| B | EnGen0047 | 2004 | NLD | 328 | GCF_000322305.1 |
| B | 1_141_733 | 2005 | USA | 327 | GCF_000157575.1 |
| B | EnGen0038 | 2006 | NLD | 331 | GCF_000322225.1 |
| B | Com15 | 2007 | USA | 583 | GCF_000157655.1 |
| B | EnGen0042 | 2001 | ESP | 289 | GCF_000322085.1 |
| B | E980 | 1998 | NLD | 94 | GCF_000172615.1 |
| B | EnGen0033 | 2000 | DEU | 299 | GCF_000322125.1 |
| B | EnGen0015 | 1998 | NLD | 61 | GCF_000321625.1 |
| B | EnGen0028 | 1956 | NOR | 75 | GCF_000321885.1 |
| B | LCT-EF90 | nd | CHI | 76 | GCF_000258325.1 |
| B | EnGen0029 | 1964 | NOR | 77 | GCF_000321905.1 |
| B | EnGen0026 | 2000 | DEU | 296 | GCF_000322145.1 |
| Rec | 1_231_408 | 2005 | USA | 582 | GCF_000157615.1 |
| Rec | EnGen0002 | 2001 | USA | 117 | GCF_000321665.1 |
| A1 | EnGen0013 | 1997 | ISR | 80 | GCF_000321545.1 |
| A1 | 1_230_933 | 2005 | USA | 18 | GCF_000157435.1 |
| A1 | EnGen0034 | 2001 | USA | 117 | GCF_000322205.1 |
| A1 | EnGen0046 | 2006 | NLD | 78 | GCF_000322405.1 |
| A1 | U0317 | 2005 | NLD | 78 | GCF_000172915.1 |
| A1 | E4452 | 2008 | NLD | 266 | GCF_000239115.1 |
| A1 | EnGen0054 | 1999 | ITA | 78 | GCF_000322345.1 |
| A1 | 1_231_502 | 2005 | USA | 203 | GCF_000157535.1 |
| A1 | EnGen0049 | 2010 | PRT | 78 | GCF_000322465.1 |
| A1 | EnGen0045 | 2010 | LVA | 78 | GCF_000322445.1 |
| A1 | EnGen0016 | 2000 | GBR | 64 | GCF_000321725.1 |
| A1 | EnGen0036 | 2002 | TZA | 18 | GCF_000322065.1 |
| A1 | EnGen0030 | 2002 | NLD | 325 | GCF_000322245.1 |
| A1 | E4453 | 2008 | NLD | 192 | GCF_000239095.1 |
| A1 | EnGen0051 | 2002 | DEU | 78 | GCF_000322365.1 |
| A1 | EnGen0050 | 2005 | HUN | 78 | GCF_000322385.1 |
| A1 | 1_231_410 | 2005 | USA | 17 | GCF_000157595.1 |
| A1 | Aus0004 | 1998 | AUS | 17 | GCF_000250945.1 |
| A1 | C68 | 1996 | USA | 16 | GCF_000160315.2 |
| A1 | E1162 | 1997 | FRA | 17 | GCF_000172675.1 |
| A1 | EnGen0057 | 2008 | DNK | 78 | GCF_000322425.1 |
| A2 | EnGen0018 | 2001 | ZAF | 159 | GCF_000321825.1 |
| A2 | EnGen0031 | 1960 | NLD | 22 | GCF_000321965.1 |
| A2 | EnGen0007 | 2001 | DEU | 160 | GCF_000321845.1 |
| A2 | EnGen0017 | 1998 | NLD | 92 | GCF_000321645.1 |
| A2 | EnGen0025 | 1965 | NLD | 92 | GCF_000321985.1 |
| A2 | EnGen0009 | 1994 | BEL | 21 | GCF_000321765.1 |
| A2 | E1071 | 2000 | NLD | 32 | GCF_000172655.1 |
| A2 | EnGen0032 | 1959 | NLD | 104 | GCF_000321945.1 |
| A2 | D344SRF | nd | USA | 25 | GCF_000176295.1 |
| A2 | TC6 | nd | USA | 25 | GCF_000161855.1 |
| A2 | EnGen0011 | 1998 | FRA | 26 | GCF_000321685.1 |
| A2 | E1636 | 1961 | NLD | 106 | GCF_000172835.1 |
| A2 | EnGen0010 | 1996 | NLD | 26 | GCF_000321505.1 |
| A2 | EnGen0048 | 2004 | SWE | 310 | GCF_000322325.1 |
| A2 | EnGen0005 | 1992 | GBR | 9 | GCF_000321465.1 |
| A2 | EnGen0022 | 1996 | NLD | 9 | GCF_000321525.1 |
| A2 | EnGen0043 | 2004 | NLD | 12 | GCF_000322185.1 |

| Clade | Isolate | Year of Isolation | Country | ST | GenBank accession |
|---|---|---|---|---|---|
| A2 | EnGen0027 | 1957 | NLD | 67 | GCF_000321925.1 |
| A2 | EnGen0001 | 1995 | BEL | 158 | GCF_000321805.1 |
| A2 | E1679 | 1998 | BRA | 114 | GCF_000172875.1 |
| A2 | EnGen0024 | 2001 | NLD | 210 | GCF_000322105.1 |
| A2 | EnGen0020 | 1995 | BEL | 27 | GCF_000321785.1 |
| A2 | EnGen0012 | 1995 | NLD | 27 | GCF_000321485.1 |
| A2 | EnGen0044 | 2001 | DNK | 27 | GCF_000322165.1 |
| A2 | 1_231_501 | 2005 | USA | 52 | GCF_000157555.1 |
| A2 | EnGen0004 | 1998 | ESP | 127 | GCF_000321705.1 |
| A2 | EnGen0052 | 2002 | NLD | 332 | GCF_000322285.1 |
| A2 | EnGen0039 | 1981 | NLD | 69 | GCF_000322025.1 |
| A2 | EnGen0019 | 1995 | DEU | 151 | GCF_000321585.1 |
| A2 | EnGen0040 | 1982 | NLD | 66 | GCF_000322045.1 |
| A2 | EnGen0008 | 1995 | ESP | 5 | GCF_000321605.1 |
| A2 | EnGen0021 | 2002 | NLD | 5 | GCF_000321745.1 |
| A2 | E1039 | 1998 | NLD | 42 | GCF_000174935.1 |
| A2 | EnGen0014 | 1995 | BEL | 150 | GCF_000321565.1 |
| A2 | EnGen0035 | 1979 | NLD | 66 | GCF_000322005.1 |

**Table S2. Reference genomes**. Table adapted from Table S1 of Lebreton et al. [1]. Rec=recombined A1/B, ST=sequence type

| | | | | Max pairwise distance (# SNPs) | | | | |
|---|---|---|---|---|---|---|---|---|
| | **Links** | **# isolates** | **Timespan (years)** | **Core** | **Full** | **Full rec filtered** | **Length full (Mb)** | **From phylogenomic analysis** |
| *Cluster 1* | between hosts | 2 | 0.003 | 9 | 6 | 6 | 2.943 | revealed |
| *Cluster 2* | between hosts | 3 | 0.60 | 31 | 81 | 42 | 3.081 | completed |
| *Cluster 3* | between hosts | 2 | 0.53 | 102 | 112 | 36 | 3.158 | revealed |
| *Cluster 4* | between hosts | 3 | 0.68 | 5 | 234 | 8 | 3.072 | revealed |
| *Cluster 5* | between hosts | 2 | 0.03 | 1 | 1 | 1 | 3.023 | revealed |
| *Cluster 6* | between and within-host | 7 | 0.73 | 19 | 496 | 21 | 3.087 | completed |
| *Cluster 7* | within-host | 2 | 0.55 | 6 | 4 | 4 | 2.968 | confirmed |
| *Cluster 8* | within-host | 3 | 0.2 | 1 | 60 | 8 | 2.833 | confirmed |
| *Cluster 9* | between hosts | 4 | 2.7 | 85 | 157 | 63 | 2.975 | revealed |
| *Cluster 10* | between hosts | 31 | 2.4 | 29 | 20 | 12 | 3.008 | completed |

**Table S3. Phylogenomic clusters details**. The first column indicates whether linked isolates were retrieved from several hosts (between hosts) or from one host (within-host). *Cluster 6* included 5 hosts, one of which had 3 isolates sampled longitudinally (between and within-host). Following, the number of isolates and timespan from sampling of the first to sampling of the last isolate are given.

Then, the maximum pairwise distance among isolates is shown, based on:

1. the core gene alignment used to define clusters (core)
2. the full alignment prior to recombination filtering (full)
3. the recombination-filtered alignment (full rec filtered)

See Material and Methods, *Phylogenomic analyses* for further details.

The length of the full alignment in mega base pairs (Mb) is given in the next column. Finally, the agreement of the links indicated by routine surveillance (based on epidemiological criteria and traditional typing) with those inferred based on these phylogenomic clusters is shown. *Confirmed* indicates that each link within the cluster was already suspected by routine surveillance, *completed* indicates that some links among isolates part of the cluster had been suspected by routine surveillance, but more isolates, unsuspected to be linked by routine surveillance, were connected to these. Finally *revealed* means that none of the isolates part of the cluster had been suspected to be linked by routine surveillance.

| ID | Isolate(s) | Position VRE5_P5 | Type | Length | Position Aus00004 | effect | nt change | aa change | Annotation in Aus0004 |
|---|---|---|---|---|---|---|---|---|---|
| 1 | VRE6_P6, VRE7_P7, VRE25_P25 | 298,217 | ins | 1 | intergenic | | | | |
| 2 | VRE6_P6 | 474,437 | snp | 1 | 499,501 | missense variant | 383T>C | Leu128Ser | MFS transporter |
| 3 | VRE6_P6 | 1,598,977 | del | 1 | 1,207,774 | frameshift variant | 339delT | Tyr116fs | tyrosine recombinase XerC |
| 4 | VRE6_P6 | 1,747,597 | snp | 1 | 22,950 | missense variant | 596G>A | Gly199Glu | DegV family protein |
| 5 | VRE6_P6 | 2,695,108 | del | 1 | 2,729,645 | frameshift variant | 27delT | Phe9fs | hypothetical protein |
| 6 | VRE6_P6 | 2,811,274 | del | 1 | 2,890,897 | frameshift variant | 622delA | Asn210fs | gfo/Idh/MocA family oxidoreductase |
| 7 | VRE6_P6 | 17,707 | snp | 1 | 1,350,210 | missense variant | 109G>A | Glu37Lys | 4-oxalocrotonate tautomerase |
| 8 | VRE6_P6 | 877,841 | del | 17 | intergenic | | | | |
| 9 | VRE6_P6 | pl2, 17,627 | snp | 1 | intergenic | | | | |
| 10 | VRE7_P7 | 414,037 | snp | 1 | 439,098 | missense variant | 161G>T | Gly54Val | PadR family transcriptional regulator |
| 11 | VRE7_P7 | 1,819,105 | snp | 1 | 1,417,538 | missense variant | 215T>A | Ile72Asn | transferase |
| 12 | VRE7_P7 | 2,770,582 | snp | 1 | 2,815,761 | synonymous variant | 3820T>C | Leu1274Leu | Enterococcal surface protein |
| 13 | VRE7_P7 | 2,770,600 | snp | 1 | 2,815,779 | missense variant | 3802A>G | Ile1268Val | Enterococcal surface protein |
| 14 | *Clade A* | pl1, 183,829 | complex | 3 | intergenic | | | | |
| 15 | *Clade A* | pl1, 183,840 | snp | 1 | intergenic | | | | |
| 16 | VRE8_P8 | 1,197,350 | snp | 1 | intergenic | | | | |
| 17 | VRE9_P9 | 1,958,423 | snp | 1 | intergenic | | | | |
| 18 | VRE12_P12 | 1,183,639 | snp | 1 | Putative symporter YjmB, not in the reference | | | | |
| 19 | VRE12_P12 | 2,002,191 | del | 1 | Hypothetical protein, not in the reference | | | | |
| 20 | *Clade B* | 2,697,718 | snp | 1 | intergenic | | | | |
| 21 | VRE18_P18 | 413,952 | snp | 1 | 439,013 | missense variant | 76T>G | Tyr26Asp | PadR family transcriptional regulator |
| 22 | VRE18_P18 | 546,408 | snp | 1 | intergenic | | | | |
| 23 | VRE18_P18 | 758,671 | snp | 1 | 756,134 | missense variant | 344C>T | Ala115Val | ABC transporter permease |
| 24 | VRE18_P18 | 1,996,708 | snp | 1 | Putative sugar transferase EpsL, not in the reference | | | | |
| 25 | VRE23_P23 | 1,200,138 | snp | 1 | 1,537,857 | missense variant | 1754A>G | Tyr585Cys | transcription antiterminator BglG |
| 26 | VRE29_P29 | 1,772,103 | snp | 1 | 1,374,716 | missense variant | 185C>T | Ser62Phe | glycosyltransferase family 4 protein |
| 27 | VRE29_P29 | 2,219,622 | snp | 1 | intergenic | | | | |
| 28 | *Clade C* | 60,123 | snp | 1 | 95,524 | missense variant | 2032G>A | Val678Ile | elongation factor G |
| 29 | *Clade C* | 389,298 | ins | 12 | 416,003 | conservative inframe insertion | 536_537ins CGAAGACAGCAA | Asn179_Glu180ins GluAspSerAsn | glycoside hydrolase, family 65 |

| ID | Isolate(s) | Position VRE5_P5 | Type | Length (bp) | Position Aus0004 | effect | nt change | aa change | Annotation in Aus0004 |
|---|---|---|---|---|---|---|---|---|---|
| 30 | *Clade C* | 1,666,026 | del | 1 | 1,271,778 | frameshift variant | 1187delA | Lys399fs | hydroxymethylglutaryl-CoA reductase, degradative |
| 31 | *Clade C* | 1,672,764 | snp | 1 | 1,278,516 | missense variant | 59C>A | Ala20Asp | cardiolipin synthase |
| 32 | *Clade C* | 2,013,711 | snp | 1 | 2,022,517 | missense variant | 1261G>A | Ala421Thr | branched-chain amino acid transport system II carrier protein |
| 33 | *Clade C* | pl1, 184,545 | del | 63 | intergenic, not in the reference | | | | |
| 34 | VRE52_P45 | 851,711 | snp | 1 | Chromosome partition protein Smc, not in the reference | | | | |
| 35 | VRE52_P45 | 882,347 | del | 1 | 1,855,498 | frameshift variant | 4delA | Met5fs | PRD domain-containing protein |
| 36 | VRE52_P45 | 2,831,679 | snp | 1 | 2,909,882 | missense variant | 683C>T | Ala228Val | ABC transporter permease |
| 37 | VRE56_P49 | 1,967,662 | snp | 1 | hypothetical protein, not in the reference | | | | |
| 38 | *Clade D* | 1,040,660 | snp | 1 | 1,707,458 | missense variant | 479G>A | Arg160His | YvcK family protein |
| 39 | VRE62_P55 | pl6, 1,812 | snp | 1 | intergenic | | | | |
| 40 | VRE63_P56 | pl1, 99,634 | snp | 1 | HTH-type transcriptional regulator DegA, not in the reference | | | | |
| 41 | VRE63_P56 | 505,856 | del | 1 | 529,666 | frameshift variant | 49delA | Ile17fs | carbohydrate ABC transporter permease |
| 42 | VRE67_P60 | 567,231 | del | 1 | intergenic | | | | |
| 43 | VRE67_P60 | 1,945,812 | snp | 1 | 1,933,533 | missense variant | 1121C>T | Thr374Ile | transglutaminase |
| 44 | VRE68_P61 | 976,781 | del | 1 | intergenic | | | | |
| 45 | VRE68_P61 | 927,293 | snp | 1 | 1,814,997 | stop gained | 2035G>T | Glu679* | LTA synthase family protein |

**Table S4. Mutations differentiating the isolates of the *persistent clone***. Mutations distinguishing each isolate of the *persistent clone* from the earliest isolate of the cluster, VRE5_P5, are reported in this table. In the rows highlighted with grey shading, mutations are shared by all isolates part of a clade. Clades are defined in Fig. 2B, where selected internal nodes are labelled with the corresponding letters. Position in VRE5_P5 refers by default to chromosomal location in bp, except where pl1, pl6, e.g., plasmid 1, plasmid 6, is specified. Columns 4-5 report the mutation type (single nucleotide polymorphism, snp; insertion, ins or deletion, del) and its length. To establish the outcome of each mutation upon translation, this variant-calling was performed against the annotated reference genome Aus0004 [16] as well. In six instances, the mutation locus did not belong to the reference genome (mutations 18, 24, 33, 34, 37, 40). In these cases, the concerned gene based on the Prokka annotation of VRE5_P5 is reported. For all other mutations, the effect on translation is specified in more details. Fs=frameshift, *=stop codon.

| Position on pELF_USZ | UniProtKB | COG/HAMAP | Gene | Product | Product (RAST) | WP_ accession |
|---|---|---|---|---|---|---|
| 3,644..4,462 | P37522 | COG1192 | soj | Sporulation initiation inhibitor protein Soj | | WP_016922501.1 |
| 6,257..6,523 | O33348 | COG2026 | relG | Toxin RelG | RelE/StbE replicon stabilization toxin | WP_010729835.1 |
| 10,867..11,634 | | | | | RepB | WP_002320780.1 |
| 13,873..14,676 | | | | | replication protein | WP_010729683.1 |
| 16,935..18,884 | O31673 | COG0542 | clpE | ATP-dependent Clp protease ATP-binding subunit ClpE | | WP_002287598.1 |
| 43,817..44,548 | O34817 | COG2188 | yvoA | HTH-type transcriptional repressor YvoA | Predicted transcriptional regulator of N-Acetylglucosamine utilization, GntR family | WP_016181189.1 |
| 44,561..45,733 | Q8XAC2 | COG2222 | agaS | D-galactosamine-6-phosphate deaminase AgaS | Galactosamine-6-phosphate isomerase AgaS | WP_123061293.1 |
| 45,756..46,763 | P63705 | COG3684 | lacD2 | Tagatose 1,6-diphosphate aldolase 2 | Tagatose 1,6-bisphosphate aldolase | WP_123061292.1 |
| 46,775..47,737 | Q833W9 | COG1105 | lacC | Tagatose-6-phosphate kinase | Tagatose-6-phosphate kinase | WP_123061291.1 |
| 47,968..48,678 | P45544 | COG2188 | frlR | putative fructoselysine utilization operon transcriptional repressor | Transcriptional regulator, GntR family | WP_016181193.1 |
| 48,849..50,627 | P48982 | | bga | Beta-galactosidase | beta-galactosidase | WP_123061290.1 |
| 50,630..51,109 | P26380 | COG3444 | levE | PTS system fructose-specific EIIB component | PTS system, galactosamine-specific IIB component | WP_016181195.1 |
| 51,122..52,018 | Q9RGG3 | COG3715 | sorC | PTS system sorbose-specific EIIC component | PTS system, galactosamine-specific IIC component | WP_048719458.1 |
| 52,005..52,814 | P69805 | COG3716 | manZ | PTS system mannose-specific EIID component | PTS system, galactosamine-specific IID component | WP_016181197.1 |
| 52,852..53,256 | P69797 | COG2893 | manX | PTS system mannose-specific EIIAB component | PTS system, galactosamine-specific IIA component | |
| 53,400..53,993 | P96631 | COG1396 | immR | HTH-type transcriptional regulator ImmR | | WP_016181199 |
| 54,972..56,120 | O31773 | COG1680 | pbpX | Putative penicillin-binding protein PbpX | Beta-lactamase class C and other penicillin binding proteins | WP_016181201.1 |
| 63,033..63,890 | | | | | Sortase A, LPXTG specific | WP_002350583 |
| 63,880..64,989 | | | | | Sortase A, LPXTG specific | WP_049055866 |
| 72,710..74,086 | P64167 | COG1674 | ftsK | DNA translocase FtsK | DNA translocase FtsK | WP_002353144.1 |
| 93,709..94,110 | | MF_01113 | dinB_1 | DNA polymerase IV | | |
| 95,486..96,172 | | MF_01113 | dinB_2 | DNA polymerase IV | DNA-directed RNA polymerase beta subunit | WP_060805128 |

**Table S5. Coding sequences of pELF_USZ with a putative product**, based on annotation by the pipelines Prokka [17] and RAST [18]. Text color reflects the colors used in Fig. 4: in brown for the genes with a putative role in DNA replication, green for those with a putative role in DNA partitioning and transfer and blue for the cargo genes of pELF_USZ. Products corresponding to *hypothetical protein*, *Mobile element protein*, *Phage-related protein*, *Phage-related protein*, *Phage protein*, *Transposase*, *XX family protein* are not reported in this table.

| Assembly | Name | GenBank acession | Size (kb) | Topology | Reads | Assembler | Publication | Max Score |
|---|---|---|---|---|---|---|---|---|
| AA708 | pELF1 | LC495616 | 143.3 | linear | Illumina/Nanopore | Unicycler | Hashimoto et al. 2019 [19] | 16,090 |
| KUHS13 | pELF2 | AP022343.1 | 108.1 | linear | Illumina/Nanopore | Unicycler | Hashimoto et al. 2020 [20] | 28,108 |
| E6043 | 3 | LR134107.1 | 55.8 | linear | Illumina/Nanopore | Unicycler | Arredondo et al. 2018 [14] | 42,957 |
| E8414 | 3 | LR135490.1 | 136.3 | linear | Illumina/Nanopore | Unicycler | Arredondo et al. 2018 [14] | 42,392 |
| E6988 | 3 | LR135245.1 | 117.2 | linear | Illumina/Nanopore | Unicycler | Arredondo et al. 2018 [14] | 42,968 |
| AUSMDU0004055 | unnamed5 | CP027511.1 | 91.1 | circular [a] | Pacbio (Illumina) | Canu, manual curation | Lee et al. 2018 [21] | 42,954 |
| AML0157 | pAML0157.4 | CP060865.1 | 94.3 | circular [b] | Illumina/Nanopore | Unicycler, manual curation | Xanthopoulou et al. 2020 [22] | 52,756 |
| VRE3363 | unnamed2 | CP064345.1 | 96.5 | linear | Illumina/Nanopore | Unicycler | Wyres et al. 2020 [23] | 42,961 |
| SSR24 | pSRR24 | CP038997.1 | 123 | linear | Illumina/Nanopore | Unicycler | Sun et al. 2020 [24] | 28,986 |
| E7196 | 2 | LR135271.1 | 173 | linear | Illumina/Nanopore | Unicycler | Arredondo et al. 2018 [14] | 43,914 |
| SC4 | p2 | CP025427.1 | 143 | linear | Pacbio | SMRT Analysis pipeline | Li et al. 2019 [25] | 42,398 |

**Table S6. Plasmid sequences compared with pELF_USZ**. The first two correspond to the linear plasmids recently identified in Japan and displayed in Fig. 4. The nine following correspond to the best hits when querying pELF_USZ against the nucleotide collection database (nr/nt), based on the maximum score (shown here in the last column) and are displayed in this order in Supplemental Fig. S11. The plasmid identifiers (assembly, name and accession) are followed by morphology descriptors: size and topology. The column "reads" indicates the sequencing platform used in each study. All plasmids sequences were reported to be linear but two, which were both subjected to manual curation (plasmid unnamed5 from AUSMDU0004055 and pAML0157.4 from AML0157).

[a] Hybrid assembly (combining Illumina reads with Pacbio reads) using Unicycler yielded a linear sequence for this plasmid.

[b] Hybrid assembly (combining Illumina reads with Nanopore reads, obtained from the authors upon request) using Unicycler yielded a linear sequence for this plasmid, as reported by the authors prior to manual curation.

| Assembly | Plasmid | # contigs | Cumulative length | % contigs | % length |
|---|---|---|---|---|---|
| **VRE53_P46** | pl1 | 28 | 193,926 | 79 | 85 |
| | pELF_USZ | 5 | 103,871 | 60 | 22 |
| | pl2 | 11 | 44,370 | 64 | 88 |
| | pl3 | 2 | 7,673 | 100 | 100 |
| | pl4 | 4 | 6,882 | 100 | 100 |
| | pl5 | 2 | 6,301 | 100 | 100 |
| | pl6 | 1 | 4,333 | 100 | 100 |
| | pl7 | 1 | 2,895 | 100 | 100 |
| | pl8 | 1 | 1,887 | 100 | 100 |
| **VRE32_P32** | pl1 | 39 | 207,421 | 82 | 95 |
| | pELF_USZ | 6 | 100,561 | 33 | 21 |
| | pl2 | 12 | 472,38 | 82 | 95 |
| | pl3 | 2 | 7,146 | 67 | 90 |
| | pl4 | 1 | 4,333 | 100 | 100 |
| | pl5 | 1 | 2,103 | 100 | 100 |
| | pl6 | 1 | 2,020 | 100 | 100 |
| | pl7 | 1 | 1,887 | 100 | 100 |

**Table S7. Accuracy of plasmid-derived contigs detection by mlplasmid** [14] for each plasmid sequence of the assemblies of VRE53_P46 and VRE32_P32 (Supplemental Fig. S18). Rows corresponding to pELF_USZ are highlighted with grey shading. # contigs: the total number of contigs of the short-read assembly mapped to a given plasmid sequence; Cumulative length: the cumulative length of all contigs mapped to a given complete plasmid sequence; % contigs: the percentage of the mapped contigs which were detected as plasmid-derived by mlplasmid; % length: percent of length that the contigs detected as plasmid-derived by mlplasmid make up within the cumulative length of all contigs mapped to the corresponding plasmid sequence.

| Isolate | Paired-end reads length (bp) | # reads | # reads after trimming | Median Depth | # contigs | Largest contig | Total Length (bp) | N50 |
|---|---|---|---|---|---|---|---|---|
| USZ_VRE1_P1 | 300 | 188,998 | 172,678 | 17.1x | 195 | 137,926 | 2,965,645 | 40,229 |
| USZ_VRE2_P2 | 300 | 371,558 | 367,812 | 28.6x | 191 | 132,441 | 2,884,091 | 48,994 |
| USZ_VRE3_P3 | 300 | 623,794 | 618,080 | 32.4x | 196 | 135,688 | 2,943,353 | 41,947 |
| USZ_VRE4_P4 | 300 | 724,612 | 714,587 | 68.8x | 211 | 141,578 | 3,098,446 | 41,485 |
| USZ_VRE5_P5 | 300 | 207,580 | 202,455 | 17.5x | 198 | 134,936 | 3,000,886 | 43,273 |
| USZ_VRE6_P6 | 300 | 237,597 | 233,072 | 22.0x | 207 | 134,936 | 2,994,548 | 43,145 |
| USZ_VRE7_P7 | 300 | 243,344 | 233,364 | 20.6x | 199 | 134,936 | 3,001,360 | 45,192 |
| USZ_VRE8_P8 | 300 | 208,831 | 205,047 | 18.5x | 203 | 134,936 | 3,000,457 | 43,145 |
| USZ_VRE9_P9 | 300 | 283,208 | 276,162 | 25.7x | 193 | 134,936 | 2,995,215 | 45,192 |
| USZ_VRE10_P10_01 | 300 | 222,498 | 217,906 | 20.7x | 200 | 134,936 | 3,002,390 | 43,145 |
| USZ_VRE11_P11 | 300 | 531,019 | 521,240 | 51.1x | 198 | 141,021 | 3,007,572 | 49,165 |
| USZ_VRE12_P12 | 300 | 235,012 | 230,371 | 21.9x | 198 | 134,936 | 2,998,503 | 43,145 |
| USZ_VRE13_P13 | 300 | 225,993 | 222,269 | 20.5x | 205 | 134,936 | 2,999,393 | 43,145 |
| USZ_VRE14_P14_01 | 300 | 349,611 | 342,193 | 31.2x | 195 | 134,936 | 3,002,886 | 45,192 |
| USZ_VRE15_P15 | 300 | 731,260 | 722,776 | 64.5x | 243 | 130,398 | 3,099,110 | 34,375 |
| USZ_VRE16_P16 | 300 | 251,640 | 247,273 | 22.7x | 217 | 134,936 | 3,009,228 | 43,145 |
| USZ_VRE17_P17 | 300 | 293,366 | 288,959 | 25.5x | 251 | 130,398 | 3,071,562 | 40,537 |
| USZ_VRE18_P18_01 | 300 | 233,183 | 213,326 | 18.0x | 218 | 130,871 | 3,012,123 | 37,511 |
| USZ_VRE19_P19 | 300 | 280,107 | 262,158 | 23.1x | 242 | 130,398 | 3,063,518 | 40,538 |
| USZ_VRE20_P20 | 300 | 259,608 | 240,690 | 20.3x | 216 | 133,536 | 3,010,393 | 36,875 |
| USZ_VRE21_P21 | 300 | 805,861 | 792,714 | 75.0x | 215 | 160,737 | 3,014,822 | 41,616 |
| USZ_VRE22_P22 | 300 | 274,678 | 253,933 | 22.4x | 221 | 133,536 | 3,014,493 | 36,875 |
| USZ_VRE23_P23 | 300 | 302,401 | 274,975 | 19.5x | 196 | 134,936 | 3,005,046 | 40,229 |
| USZ_VRE24_P24 | 300 | 260,092 | 244,653 | 22.4x | 197 | 145,865 | 3,063,088 | 44,027 |
| USZ_VRE25_P25 | 300 | 302,662 | 298,420 | 27.4x | 201 | 134,936 | 2,992,835 | 40,302 |
| USZ_VRE26_P26_01 | 300 | 220,262 | 206,808 | 17.6x | 221 | 155,322 | 2,956,670 | 46,448 |
| USZ_VRE27_P27 | 300 | 222,582 | 208,064 | 18.0x | 213 | 145,772 | 2,930,951 | 44,182 |
| USZ_VRE28_P28 | 300 | 792,676 | 783,992 | 71.1x | 179 | 145,847 | 3,152,729 | 50,258 |
| USZ_VRE29_P29 | 300 | 247,517 | 234,640 | 20.2x | 213 | 130,871 | 3,002,442 | 37,256 |
| USZ_VRE30_P30 | 300 | 249,102 | 236,305 | 20.1x | 264 | 116,456 | 3,069,422 | 36,363 |
| USZ_VRE31_P31 | 300 | 357,738 | 333,250 | 29.7x | 189 | 134,814 | 2,982,065 | 43,145 |
| USZ_VRE32_P32 | 300 | 311,934 | 303,667 | 26.4x | 232 | 116,456 | 3,115,078 | 36,590 |
| USZ_VRE33_P33 | 300 | 396,589 | 363,613 | 30.5x | 202 | 135,874 | 3,041,716 | 50,258 |
| USZ_VRE34_P34 | 300 | 678,797 | 669,725 | 66.6x | 207 | 107,869 | 2,934,248 | 45,716 |
| USZ_VRE35_P35 | 300 | 479,838 | 474,306 | 39.2x | 231 | 107,869 | 3,087,860 | 47,366 |
| USZ_VRE36_P36 | 300 | 254,337 | 232,623 | 19.0x | 283 | 132,304 | 3,127,038 | 33,028 |
| USZ_VRE37_P18_02 | 300 | 208,070 | 195,449 | 17.7x | 201 | 160,730 | 2,822,971 | 46,448 |
| USZ_VRE38_P37 | 300 | 293,987 | 286,709 | 24.4x | 223 | 145,865 | 3,206,188 | 37,094 |
| USZ_VRE39_P10_02 | 300 | 277,536 | 258,391 | 22.0x | 261 | 132,304 | 3,069,159 | 36,363 |
| USZ_VRE40_P38 | 300 | 277,828 | 256,697 | 20.5x | 227 | 145,865 | 3,207,347 | 36,826 |
| USZ_VRE41_P18_03 | 300 | 361,381 | 338,476 | 34.6x | 190 | 160,730 | 2,798,818 | 46,448 |
| USZ_VRE42_P26_02 | 300 | 311,309 | 289,484 | 27.0x | 199 | 155,322 | 2,944,124 | 53,738 |
| USZ_VRE43_P39 | 300 | 293,001 | 272,994 | 23.0x | 192 | 130,396 | 3,047,635 | 49,140 |
| USZ_VRE44_P40 | 300 | 1,095,012 | 1,082,772 | 100.3x | 218 | 134,936 | 2,937,479 | 39,950 |
| USZ_VRE45_P18_04 | 300 | 331,298 | 307,334 | 27.9x | 185 | 160,730 | 2,793,779 | 46,448 |
| USZ_VRE46_P41_01 | 150 | 712,103 | 703,529 | 19.1x | 305 | 116,976 | 3,154,988 | 33,020 |
| USZ_VRE47_P41_02 | 150 | 892,869 | 887,090 | 26.9x | 239 | 116,976 | 3,110,646 | 33,476 |
| USZ_VRE48_P42 | 300 | 465,594 | 460,360 | 44.1x | 246 | 160,730 | 2,902,516 | 46,310 |
| USZ_VRE49_P41_03 | 150 | 660,657 | 652,402 | 19.8x | 288 | 116,976 | 3,055,697 | 34,294 |
| USZ_VRE50_P43 | 150 | 491,971 | 487,442 | 17.1x | 201 | 100,767 | 2,951,627 | 43,173 |
| USZ_VRE51_P44 | 150 | 582,731 | 578,803 | 20.1x | 195 | 136,045 | 2,962,147 | 39,166 |
| USZ_VRE52_P45 | 300 | 387,897 | 382,951 | 31.4x | 226 | 107,639 | 3,107,702 | 39,274 |

| Isolate | Paired-end reads length (bp) | # reads | # reads after trimming | Median Depth | # contigs | Largest contig | Total Length (bp) | N50 (bp) |
|---|---|---|---|---|---|---|---|---|
| USZ_VRE53_P46 | 300 | 603,441 | 596,793 | 43.3x | 227 | 107,639 | 3,106,978 | 37,388 |
| USZ_VRE54_P47 | 300 | 613,221 | 605,544 | 47.0x | 230 | 107,639 | 3,106,076 | 39,274 |
| USZ_VRE55_P48 | 300 | 408,438 | 401,631 | 39.1x | 224 | 121,414 | 2,989,870 | 44,541 |
| USZ_VRE56_P49 | 300 | 283,151 | 279,483 | 20.6x | 233 | 107,639 | 3,106,040 | 37,511 |
| USZ_VRE57_P50 | 300 | 510,174 | 504,150 | 31.9x | 245 | 100,943 | 3,103,210 | 37,388 |
| USZ_VRE58_P51 | 300 | 381,811 | 374,629 | 26.2x | 227 | 107,639 | 3,106,280 | 39,307 |
| USZ_VRE59_P52 | 300 | 1,079,474 | 1,065,244 | 84.6x | 230 | 107,639 | 3,104,905 | 37,511 |
| USZ_VRE60_P53 | 300 | 644,097 | 636,847 | 48.9x | 224 | 107,639 | 3,108,538 | 37,388 |
| USZ_VRE61_P54 | 300 | 397,480 | 391,251 | 30.3x | 223 | 107,639 | 3,106,863 | 37,511 |
| USZ_VRE62_P55 | 300 | 300,288 | 296,966 | 20.8x | 228 | 107,639 | 3,109,175 | 37,388 |
| USZ_VRE63_P56 | 300 | 523,296 | 515,322 | 41.5x | 230 | 107,638 | 3,107,149 | 37,511 |
| USZ_VRE64_P57 | 300 | 589,128 | 579,000 | 48.2x | 240 | 107,639 | 3,105,773 | 37,388 |
| USZ_VRE65_P58 | 300 | 341,239 | 332,849 | 20.0x | 227 | 107,639 | 3,104,665 | 37,511 |
| USZ_VRE66_P59 | 300 | 940,465 | 929,235 | 67.9x | 239 | 107,639 | 3,107,995 | 37,511 |
| USZ_VRE67_P60 | 300 | 399,252 | 391,734 | 36.1x | 215 | 107,638 | 3,004,756 | 37,511 |
| USZ_VRE68_P61 | 300 | 529,504 | 520,533 | 43.2x | 222 | 107,639 | 3,106,257 | 37,511 |
| USZ_VRE69_P14_02 | 300 | 217,928 | 212,534 | 18.0x | 230 | 107,639 | 3,106,027 | 37,388 |

**Table S8. Quality metrics of the sequences.** Columns 2-4 report raw sequences quality metrics and columns 5-9 report assembly quality metrics. N50 is the length of the shortest contig within the set of largest contigs making up 50% of the total genome length.

#### References

1. Lebreton, F. *et al.* Emergence of Epidemic Multidrug-Resistant Enterococcus faecium from Animal and Commensal Strains. *mBio* **4**, e00534-13 (2013).

2. Arthur, M., Molinas, C., Depardieu, F. & Courvalin, P. Characterization of Tn1546, a Tn3-related transposon conferring glycopeptide resistance by synthesis of depsipeptide peptidoglycan precursors in Enterococcus faecium BM4147. *J. Bacteriol.* **175**, 117–127 (1993).

3. Wardal, E. *et al.* Diversity of plasmids and Tn1546-type transposons among VanA Enterococcus faecium in Poland. *Eur. J. Clin. Microbiol. Infect. Dis.* **36**, 313–328 (2017).

4. van Hal, S. J. *et al.* Evolutionary dynamics of Enterococcus faecium reveals complex genomic relationships between isolates with independent emergence of vancomycin resistance. *Microb. Genomics* **2**, (2016).

5. Humphries, R. M. The New, New Daptomycin Breakpoint for Enterococcus spp. *J. Clin. Microbiol.* **57**, (2019).

6. Zhang, X. *et al.* Identification of a Genetic Determinant in Clinical Enterococcus faecium Strains That Contributes to Intestinal Colonization During Antibiotic Treatment. *J. Infect. Dis.* **207**, 1780–1786 (2013).

7. Wick, R. R., Schultz, M. B., Zobel, J. & Holt, K. E. Bandage: interactive visualization of de novo genome assemblies. *Bioinformatics* **31**, 3350–3352 (2015).

8. Bankevich, A. *et al.* SPAdes: A New Genome Assembly Algorithm and Its Applications to Single-Cell Sequencing. *J. Comput. Biol.* **19**, 455–477 (2012).

9. Wick, R. R., Judd, L. M., Gorrie, C. L. & Holt, K. E. Unicycler: Resolving bacterial genome assemblies from short and long sequencing reads. *PLoS Comput. Biol.* **13**, (2017).

10. Konstantinidis, K. T. & Tiedje, J. M. Genomic insights that advance the species definition for prokaryotes. *Proc. Natl. Acad. Sci. U. S. A.* **102**, 2567–2572 (2005).

11. Ondov, B. D. *et al.* Mash: fast genome and metagenome distance estimation using MinHash. *Genome Biol.* **17**, 132 (2016).

12. Sullivan, M. J., Petty, N. K. & Beatson, S. A. Easyfig: a genome comparison visualizer. *Bioinforma. Oxf. Engl.* **27**, 1009–1010 (2011).

13. Zuker, M. Mfold web server for nucleic acid folding and hybridization prediction. *Nucleic Acids Res.* **31**, 3406–3415 (2003).

14. Arredondo-Alonso, S. *et al.* mlplasmids: a user-friendly tool to predict plasmid- and chromosome-derived sequences for single species. *Microb. Genomics* **4**, e000224 (2018).

15. Inouye, M. *et al.* SRST2: Rapid genomic surveillance for public health and hospital microbiology labs. *Genome Med.* **6**, 90 (2014).

16. Lam, M. M. C. *et al.* Comparative Analysis of the First Complete Enterococcus faecium Genome. *J. Bacteriol.* **194**, 2334–2341 (2012).

17. Seemann, T. Prokka: rapid prokaryotic genome annotation. *Bioinformatics* **30**, 2068–2069 (2014).

18. Aziz, R. K. *et al.* The RAST Server: Rapid Annotations using Subsystems Technology. *BMC Genomics* **9**, 75 (2008).

19. Hashimoto, Y. *et al.* Novel Multidrug-Resistant Enterococcal Mobile Linear Plasmid pELF1 Encoding vanA and vanM Gene Clusters From a Japanese Vancomycin-Resistant Enterococci Isolate. *Front. Microbiol.* **10**, (2019).

20. Hashimoto, Y. *et al.* First Report of the Local Spread of Vancomycin-Resistant Enterococci Ascribed to the Interspecies Transmission of a vanA Gene Cluster-Carrying Linear Plasmid. *mSphere* **5**, (2020).

21. Lee, R. S. *et al.* The changing landscape of vancomycin-resistant Enterococcus faecium in Australia: a population-level genomic study. *J. Antimicrob. Chemother.* **73**, 3268–3278 (2018).

22. Xanthopoulou, K. *et al.* Characterization of a vancomycin-resistant Enterococcus faecium isolate and a vancomycin-susceptible E. faecium isolate from the same blood culture. *J. Antimicrob. Chemother.* (2020) doi:10.1093/jac/dkaa532.

23. Wyres, K. L. *et al.* Genomic surveillance of antimicrobial resistant bacterial colonisation and infection in intensive care patients. *medRxiv* 2020.11.03.20224881 (2020) doi:10.1101/2020.11.03.20224881.

24. Sun, L. *et al.* Tandem amplification of the vanM gene cluster drives vancomycin resistance in vancomycin-variable enterococci. *J. Antimicrob. Chemother.* **75**, 283–291 (2020).

25. Li, N. *et al.* Horizontal transfer of vanA between probiotic Enterococcus faecium and Enterococcus faecalis in fermented soybean meal and in digestive tract of growing pigs. *J. Anim. Sci. Biotechnol.* **10**, 36 (2019).
